# Protective near-germline antibodies to malaria elicited by RTS,S/AS01_E_ vaccination in African infants

**DOI:** 10.64898/2026.10.01.755937

**Authors:** Emma Bloch, Fiona Devys, Fabien Cannac, Monika Jain, Katrina Broadbent, Carla Sánchez, Wen-Hsin Lee, Chenjerai Jairoce, Pedro Aide, Joseph J. Campo, Dídac Macià, Carlo Carolis, Ian A. Wilson, Andrew B. Ward, Rogerio Amino, Gemma Moncunill, Carlota Dobaño, Luis M. Molinos-Albert

## Abstract

Malaria is a leading cause of childhood mortality, yet infant immune responses to vaccination remain poorly understood. Here, we characterize 88 monoclonal antibodies (mAbs) directed against the *Plasmodium falciparum* circumsporozoite protein cloned from African infants immunized with the RTS,S/AS01_E_ malaria vaccine, identifying potent antiparasitic mAbs exhibiting molecular features resembling those observed in adults and minimal somatic hypermutation. Importantly, germline-reverted variants of four mAbs inducing sterile protection *in vivo* maintained high binding affinity and Fab-dependent parasite killing. Structural analyses revealed that a few amino acid replacements drive homotypic helical assemblies, fine-tuning parasite inhibition and protective efficacy. Thus, despite maturational constraints of early life, infants can generate potent antimalarial antibodies in a near-germline configuration, informing next-generation vaccine development for this vulnerable group.

## Introduction

Child mortality remains a major global health challenge. About 3.7 million children under one year of age die annually, and most of these are caused by infectious diseases (*1*). The optimization of immune interventions to control infections in this group requires a deeper understanding of the development and functional capacity of the immune system during early life. This is particularly urgent for malaria, a devastating disease that caused an estimated 282 million cases and more than 600,000 deaths in 2024, with children under 5 years accounting for 75% of mortality within the sub-Saharan African region (*2*).

Malaria is a parasitic mosquito-borne disease caused by *Plasmodium spp*., of which the deadliest is *Plasmodium falciparum* (*Pf*) (*2*). *Pf* is an apicomplexan parasite whose infective form, sporozoite (SPZ), is densely decorated by the circumsporozoite protein (*Pf*CSP) (*3*), which has a crucial role in the establishment of infection in the human host, and is a major target of neutralizing antibodies (*4*, *5*). The structure of *Pf*CSP in the 3D7 strain comprises a flexible, proteolytically processable N-terminal domain (NTD), containing the heparan sulfate binding site for hepatocyte attachment; a disordered, central repeat region (CRR) composed by 38 NANP (Asn-Ala-Asn-Pro), interspersed by 4 NVDP (Asn-Val-Asp-Pro) and 1 NPDP (Asn-Pro-Asp-Pro) tetrapeptide motifs; and a well-ordered conformational C-terminal domain (CTD), which is anchored to the parasite membrane through a glycosylphosphatidylinositol (GPI) (*6*, *7*). During a mosquito bite, SPZs are injected into the skin, subsequently entering the systemic circulation to arrest in the liver within minutes, where they extravasate to invade hepatocytes (*8*). Here, SPZs take about five days to develop into thousands of merozoites which exit the liver to enter in the circulation where they infect erythrocytes, causing malaria symptoms (*9*). The small number of SPZ injected by the mosquito (*10*), together with the different cellular barriers the parasite needs to cross to reach the liver, makes the pre-erythrocytic stage of *Pf* life cycle an optimal window for vaccine-induced antibodies to prevent malaria (*11*). Crucially, an effective pre-erythrocytic vaccine requires the sustained elicitation of antibodies capable of rapidly neutralize SPZ during their migration from the skin to the liver (*11*, *12*).

While vaccine candidates targeting different stages of *Pf* life cycle are under development, pre-erythrocytic vaccines are the most advanced. RTS,S/AS01_E_ (RTS,S, Mosquirix, developed by GlaxoSmithKline), a subunit vaccine targeting the 3D7 strain of *Pf*CSP, was the first malaria vaccine approved for use in African children (*13–15*). RTS,S is based on virus-like particles displaying a *Pf*CSP truncated form spanning 18.5 NANP motifs and the CTD, linked to the hepatitis B virus surface antigen (HBsAg) and formulated and administered with the AS01_E_ adjuvant (*16*). Together with R21/Matrix-M (developed by the University of Oxford’s Jenner Institute) (*17*, *18*), RTS,S has been integrated into immunization programmes under the WHO auspices in children older than 5 months living in malaria-endemic regions of Africa (*15*). While its implementation constitutes a milestone for global health, RTS,S presents important limitations, including modest vaccine efficacy and waning immunity (*14*, *19*), which demand further investigation to identify potent antibody responses to inform next-generation immunogens.

Immune responses in early life are generally weaker and shorter than in adults, resulting in higher susceptibility to infection. Hence, vaccination strategies require overcoming certain maturational constraints, including lower T-follicular helper cell frequencies, limited germinal center B-cell responses, and reduced immunoglobulin (Ig) class-switch recombination and somatic hypermutation (SHM) (*20*, *21*). Despite these limitations, increasing evidence suggest that infants can mount robust antibody responses to certain pathogens, as illustrated by the relatively fast development of broadly neutralizing antibodies against the human immunodeficiency virus (HIV-1) and respiratory syncytial virus (RSV) (*22*, *23*). Furthermore, repertoire sequencing analysis showed that B-cell clonal selection and diversification mechanisms are partially developed in infants upon malaria infection (*24*). The *Pf*CSP human B-cell antibody response has been widely investigated in adults (*25–33*) leading to significant discoveries, which culminated in the isolation of potent monoclonal antibodies (mAbs), some of them currently under clinical trials (*34*), and novel promising vaccine prototypes (*35–37*). However, the molecular attributes of the B-cell response, and whether similarly potent and protective *Pf*CSP antibodies are elicited in infants living in endemic areas have yet to be investigated. To address this knowledge gap, we characterized a collection of *Pf*CSP mAbs cloned from the circulating B cells of RTS,S-vaccinated Mozambican infants, and defined the ontogeny and structural features that drive *Pf*CSP recognition, SPZ neutralization, and protection against malaria in this vulnerable population.

## Results

### RTS,S elicits a polyclonal IgG *Pf*CSP^+^ B-cell response in Mozambican infants

MAL067 was an ancillary immunology study nested within the multicenter, randomized, observer-blind phase 3 clinical trial MAL055 (NCT00866619) designed to evaluate the efficacy, safety and immunogenicity of the RTS,S vaccine in young children aged 6 weeks to 17 months from seven African countries (*13*, *38*). During the primary vaccination phase RTS,S was administered at months (M)0, M1 and M2, and blood samples collected by M0 and M3, one month after the third dose. Extensive immuno-epidemiological investigations on MAL067 have been previously performed by our group and others (*39–43*).

To investigate the B-cell antibody response to *Pf*CSP in MAL067 participants after primary vaccination, we produced a recombinant protein (CSP18) containing 18.5 NANP repetitions and the CTD of the 3D7 *Pf*CSP strain, without the GPI, which was strongly recognized by benchmarked *Pf*CSP antibodies (*34*) (**fig. S1A and B**). In addition, we used a representative CRR peptide (NANP_10_) and a truncated CTD protein, to map the response towards both *Pf*CSP domains (**fig. S1A and B**). We selected six donors (d) [aged 9-weeks (d4, d5, d6, d9); six (d3) and eight (d8) months prior vaccination, **table S1**] from the Manhiça (Mozambique) site, developing high plasma IgG antibody titers to *Pf*CSP post-primary vaccination (M3), as determined by enzyme-linked immunosorbent assay (ELISA) (**Fig. 1A-B**). Peripheral blood B cells collected at M3 were stained with fluorescently labelled CSP18 and analyzed by flow cytometry (**Fig. 1C and fig. S1C**). Analysis of the Ig isotypes revealed that CSP18-reactive memory B cells (MBC, defined as CD19^+^, IgD^−^, and/or CD27^+^) were predominantly IgG (50-83%), while IgM ranged between 7% and 35%, and IgA represented a small fraction (**fig. S1D**). CSP18-reactive IgG^+^ B cells were detected at frequencies ranging from 0.41 to 1.4% (**Fig. 1C**), and were mainly resting (CD27^+^CD21^+^) and activated (CD27^+^CD21^−^) MBC (**fig. S1E**). From 408 CSP18^+^ B cells captured by single-cell flow cytometric sorting, 193 heavy and 332 light variable domain genes were amplified by reverse transcription polymerase chain reaction (RT-PCR). From the 182 matched antibody sequences, we produced 109 unique IgG mAbs by recombinant expression cloning (*44*). A total of 88 [81% (76 to 90%)] antibodies efficiently recognized CSP18 by ELISA (**Fig. 1D and E, fig. S2A-D and table S2**). Only 7 of the 88 CSP18 antibodies (8%) were part of clonally expanded B-cell lineages, which were found in all donors except d3 (**Fig. 1D**). ELISA analysis identified 33 antibodies out of the 88 binding to NANP_10_ [38% (11-60%)], whereas the remaining 55 antibodies recognized the CTD [62% (40-89%)] (**Fig. 1F and fig. S2A and E-H**). Thus, except for d5, a minority of the expressed CSP18 antibodies recognized the CRR of *Pf*CSP. Although antibodies similarly recognizing both CRR and CTD are rare (*26*), we identified one antibody with this profile (ISG9.108, **fig. S2A**). We conclude that RTS,S primary vaccination induced a polyclonal, IgG-switched *Pf*CSP-reactive B-cell response in Mozambican infants.

**Fig. 1.**
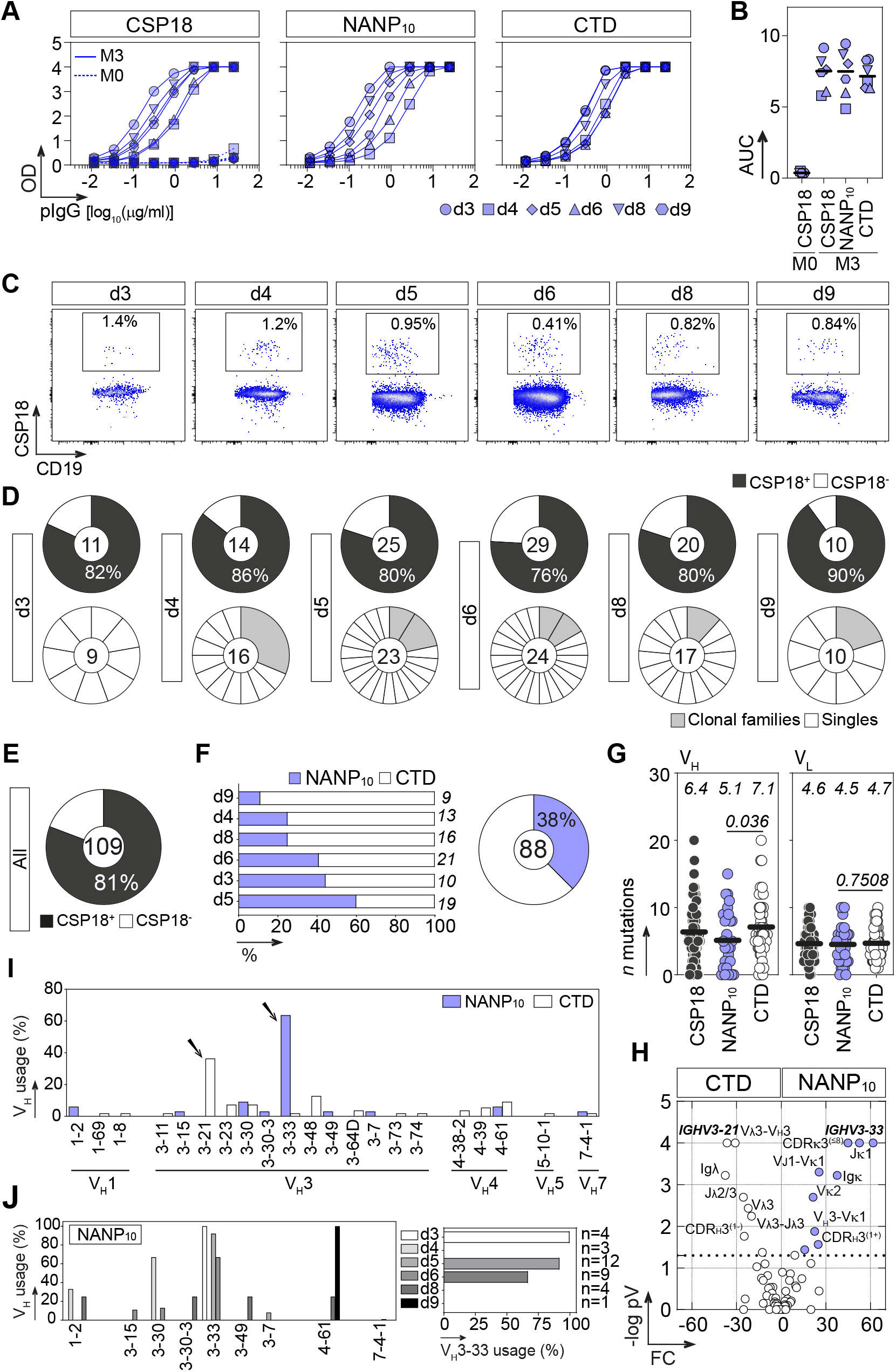
*Pf*CSP B-cell antibodies from RTS,S/AS01_E_ vaccinated African infants. (**A**) ELISA titration graphs showing the reactivity of purified plasma IgG from RTS,S vaccinated donors (d) against the CSP18 construct at month (M)0 (dashed line) and M3 (solid line), and NANP_10_ and CTD proteins at M3. (**B**) Dot plot comparing the area under the curve (AUC) values derived from the ELISA titration curves in (A). (**C**) Flow cytometry plots showing the percentage of circulating blood CSP18^+^ IgG B cells. (**D**) Individual pie charts showing the proportion of CSP18-reactive antibodies (top), and the distribution (bottom) of clonally expanded (grey) and unique (white) IgG B-cell clones with confirmed CSP18 specificity, with only one representative IgG antibody tested per clonal family, as presented in the top. The number of antibodies is indicated in the center of the pie chart, and the size of the grey slices is proportional to the number of clonal members. (**E**) Pie chart summarizing the proportion of CSP18-reactive antibodies. The number of antibodies tested is indicated in the center of the pie chart. (**F**) Bar graph (left) and pie chart (right) showing the proportion of IgG antibodies recognizing the NANP_10_ and CTD proteins per donor and in total, respectively. The number of antibodies per donor is indicated on the right of the bar graphs in italics. (**G**) Dot plots comparing the number of V_H_ and V_L_ gene mutations of all the CSP18 antibodies (n=88, black), or grouped by NANP_10_ (n=33, blue) and CTD (n=54, white) reactivity. The average number of mutations is indicated in italics. Groups were compared with the two-tailed Mann-Whitney test. *p* values are indicated. (**H**) Volcano plot analysis comparing the immunoglobulin gene repertoire of NANP_10_- and CTD-reactive IgG antibodies. (**I**) Bar graph comparing the frequency distribution of single V_H_ gene families expressed by NANP_10_ and CTD IgG antibodies. (**J**) Same as in (I) but for each donor (left), and bar graph showing the frequency of *IGHV3-33*-encoded (right), of NANP_10_ IgG antibodies.

### *Pf*CSP antibodies elicited during early life present molecular features resembling adults

Ig genes of CSP18 antibodies presented low levels of SHM (average 6.4[0-20] and 4.6[0-10] nucleotide mutations for IgH and IgL, respectively) (**Fig. 1G and fig. S3A**), suggesting the selection of high affinity precursors from the naïve B-cell repertoire. SHM rates were comparable to non-antigen enriched B cells from African infants (*24*), and substantially lower than RTS,S-induced *Pf*CSP^+^ B cells from malaria naive adults following a similar vaccination schedule (**fig. S3B**) (*31*). As reported in adults (*31*), CTD antibodies presented slightly higher IgH mutations (average 7.1[0-20] *versus* 5.1[0-15] nucleotide mutations, *p*=0.036, for CTD and CRR, respectively) (**Fig. 1G**). Analysis of the Ig molecular features revealed that CSP18 antibodies frequently used the *IGHV3* gene family, with a strong enrichment in *IGHV3-33* and *IGHV3-21* segments for CRR and CTD, respectively (**Fig. 1H and I and fig. S3C**), as similarly shown in adult *Pf*CSP responses (*27*, *28*, *31*). All CRR *IGHV3-33* antibodies were encoded by *IGHV3-33*01* or *IGHV3-33*06* alleles (**table S2**), both presenting a strictly conserved tryptophan residue at position 52 (W52) in the CDRH2, which critically drives germline (GL) reactivity and *in vitro* parasite inhibition (*45*). Most antibodies expressed the IgG1 subclass, and an average CDRH3 length of 13 amino acids (**fig. S3D and E**). In addition, CRR antibodies presented a higher frequency of positively charged CDRH3 amino acid residues, and a strong bias towards kappa light chain usage (82% *versus* 50% for CRR and CTD, respectively, *p*=0.0081) (**Fig. 1H and fig. S3D, F and G**). Notably, 71% of *IGHV3-33* CRR antibodies (n=21) were matched with an 8-amino acid-length CDRL3, most of them encoded by the *IGKV1-5* gene segment and, to a lesser extent, *IGKV3-11* and *IGKV3-15* (**fig. S3H and table S2**), being a hallmark of potent antibodies (*45*). Likewise, *IGHV3-21* CTD antibodies were frequently matched with *IGLV3-21* light chain (**Fig. S3I**) (*28*). We only identified *IGHV3-33* antibodies in three [d3, 100% (n=4); d5, 92% (n=12); d6, 67% (n=9)] out of the six donors (**Fig. 1J**), suggesting differential selection and expansion of B-cell clones or donor-dependent allelic variability (*46*). Interestingly, donors lacking *IGHV3-33* antibodies (d4, d8 and d9) presented the lowest proportion of CRR antibodies (**Fig. 1F**), supporting the selective advantage of *IGHV3-33* precursors in the evolution of CRR B-cell responses (*27*). Notably, *IGHV3-33*/*IGKV1-5* CRR antibodies presented selected replacement mutations commonly found in adults, which frequently involve position V50 in CDRH2 and, to a lesser extent, S31 of CDRH1 (**fig. S4**), both associated with antigen-driven selection mediating strong affinity for short NANP repetitions (*45*). In addition, we detected frequent amino acid substitutions (Y32F in the CDRH1, N56K and K64E in CDRH2, and Y94L in the CDRL3) known to promote inter-Fab homotypic contacts, high avidity binding, B-cell activation and improved *in vivo* protection (*45*, *47*) (**fig. S4**). Thus, *Pf*CSP B-cell antibodies elicited by RTS,S in these infants present molecular features resembling adults, and experienced refinement through antigen-driven selection, presumably in germinal centers, despite low somatic mutation rates.

### RTS,S elicits potent polyfunctional antibodies in infants

*Pf*CSP antibodies can provoke Fab-dependent cytotoxicity and the induction of a thread-like precipitate at the posterior and apical end of the SPZ (circumsporozoite precipitation reaction, CSPR) resulting in its elongation, subsequent motility loss, and parasite death (*48–52*). Both processes represent sequential stages compromising parasite fitness and downstream activities like cell invasion and traversal (*11*). Most of protection conferred by *Pf*CSP antibodies is related to SPZ neutralization during their migration in the skin and strongly correlates with Fab-dependent cytotoxicity (*53*). To broadly define the antiparasitic properties of CSP18 antibodies, we used *P. berghei* (*Pb*) SPZs knockout for the *Pb*CSP, expressing the full-length *Pf*CSP (PbPf). We first examined direct parasite killing (Fab-dependent cytotoxicity) in suspension (*53*). Interestingly, 27 out of 33 CRR antibodies were highly cytotoxic, compromising parasite viability below 25% when tested at 100 µg/ml. In striking contrast, CTD antibodies barely compromised SPZ viability (**Fig. 2A and fig. S5A**), in line with the limited exposure of the CTD on the surface of SPZ (*54*). Of note, 13 CRR antibodies exhibited half-maximal lethal effective concentrations (EC_50_) below 10 µg/ml, being as potent as the benchmarked antibody 317 (**Fig. 2B and fig. S5B and D**) (*34*). Simultaneous quantification of the CSPR revealed most CRR antibodies triggered a medium- or long-elongated SPZ phenotype, in contrast with CTD antibodies which induced a short or no CSPR (**Fig. 2C and fig. S5C and D**). CSPR values were strongly associated with lethal EC_50_, i.e., the most potent cytotoxic antibodies triggered a middle- and long-elongated CSPR phenotype (**Fig. 2D**), reinforcing that SPZ lethality is linked with *Pf*CSP shedding (*53*).

**Fig. 2.**
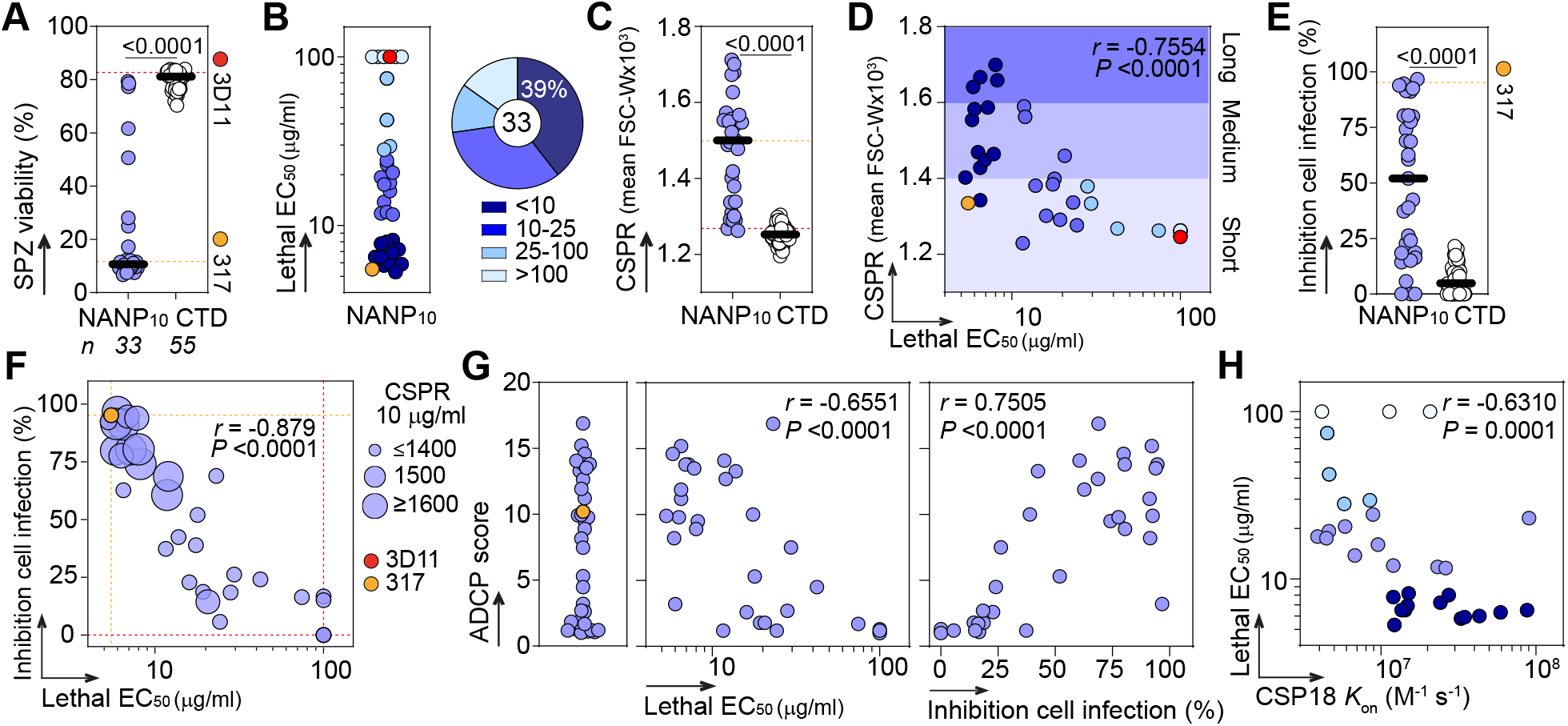
*In vitro* antiparasitic properties *Pf*CSP B-cell antibodies. (**A**) Dot plot comparing the Fab-dependent lethality induced in sporozoites (SPZ) by NANP_10_ and CTD antibodies tested at 100 µg/ml, after 45 min of incubation at 37°C, in duplicate in two independent experiments. (**B**) Dot plot (left) showing lethal half-maximal effective concentration (EC_50_) values induced in SPZ by NANP_10_ antibodies (n=3 independent experiments, see fig. S5C). The pie chart (right) shows the distribution of NANP_10_ antibodies by EC_50_ values (µg/ml) color coded in shades of blue, with darker shades indicating higher potency. (**C**) Dot plot comparing the circumsporozoite precipitation reaction (CSPR) induced in SPZs by NANP_10_ and CTD antibodies tested as in (A). (**D**) Correlation plot showing the induced CSPR phenotype, tested at 10 µg/ml versus the lethal EC_50_ values of NANP_10_ antibodies. CSPR is coded in shades of blue with darker shades indicating a more elongated phenotype (long-, medium-elongated, short round). Dots are color coded as in (B). (**E**) Dot plot comparing the percentage of SPZ inhibition of cell infection induced by NANP_10_ and CTD antibodies, after two hours of incubation at 37°C, tested at 3 µg/ml in 2-3 independent experiments (see fig. S6B). (**F**) Bubble plot showing the lethal EC_50_ values *versus* the percentage of SPZ inhibition of cell infection induced by NANP_10_ antibodies. The size of the bubbles is proportional to the elongated CSPR phenotype induced by antibodies at 10 µg/ml. (**G**) Dot plot (left) showing the antibody-dependent cellular phagocytosis (ADCP) score values of NANP_10_ antibodies tested in duplicate at 0.1 µg/ml, in at least two independent experiments. The correlation plots (right) show the ADCP scores *versus* the lethal EC_50_, and the percentage of inhibition of cell infection. (**H**) Correlation plot showing the CSP18 association constants (*K*_on_) *versus* the lethal EC_50_ values of NANP_10_ antibodies. Dots are color coded as in (B). The values for the cytotoxic and inhibitory *Pf* 317 antibody (orange), and the *Pb* 3D11 (red) negative control are indicated in colored dashed lines or dots. Groups were compared with the two- tailed Mann- Whitney test. *p* values are indicated. The two-sided Spearman rho correlation coefficients and corresponding *p* values are shown in (D), (F), (G) and (H).

Potent *Pf*CSP antibodies are also able to neutralize SPZs in the liver (*53*) by a cytotoxicity independent mechanism (*52*), probably associated with the inhibition of hepatocyte infection (*55*). We thus sought to examine the ability of the antibodies to inhibit PbPf SPZ infection of a hepatoma cell line HepG2 (*56*). To strictly detect cell infection inhibitory activities, antibodies were tested at non-cytotoxic concentrations [1 and 3 µg/ml, (**fig. S5D**), analyzed 48 h and 2 h post-inoculation, respectively]. At 3 µg/ml, CRR antibodies exhibited significantly superior inhibitory activities as compared with CTD antibodies (median inhibition 50.1% [0-97] *versus* 4.8% [0-21] for CRR and CTD, respectively), with a subset of 7 above 90% inhibition (**Fig. 2E and fig. S6A and B**). Expectedly, lowering the concentration to 1 µg/ml markedly reduced the inhibitory activity of CRR antibodies, with the top seven showing between 25% and 50% of inhibition (**fig. S6C**). Notably, highly cytotoxic and inhibitory NANP_10_ antibodies were strongly associated, and frequently induced a middle- and long-elongated CSPR phenotype, whereas weak antibodies induced a short or no CSPR (**Fig. 2F and fig. S6D**).

*Pf*CSP antibodies can mediate Fc-dependent effector functions, which can confer protection against clinical malaria (*57*), while improved FcyR binding can modestly increase the protective capacity of CIS43 antibody (*58*). To investigate the opsonization capacity, and ability of CRR antibodies to promote phagocytosis (antibody-dependent cellular phagocytosis, ADCP), we used an *in vitro* assay consisting of fluorescently-labelled *Pf*CSP-coated beads and the monocytic THP-1 line as effector cells. We found that 18 antibodies displayed strong ADCP scores, which were globally correlated with Fab-dependent lethality and cell infection inhibition (**Fig. 2G**). Here, high ADCP scores likely indicate an efficient avidity binding to *Pf*CSP which may translate into higher apparent affinity and stronger SPZ opsonization load, leading to highly potent cytotoxic and inhibitory activities. In fact, the association rates (*K*_on_), measured by surface plasmon resonance (SPR, **fig. S7 and table S3**) correlated with antiparasitic functions (**Fig. 2H and fig. S6E**), suggesting that fast *Pf*CSP binding determines an efficient SPZ neutralization, and subsequent inhibition of hepatocyte invasion. Altogether, our findings identify potent CRR antibodies elicited by RTS,S in infants, that neutralize SPZ through coordinated antiparasitic mechanisms.

### Potent antiparasitic CRR antibodies are encoded by *IGHV3-33* and other gene families

Despite being a public signature of potent CRR antibodies (*34*, *59*), we identified a few poorly cytotoxic *IGHV3-33* some of them lacking somatic mutations in both heavy (HC) and light (LC) chains (ISG5.154), or in the HC only (ISG5.147, ISG6.170) (**fig. S8**), indicating an intermediate maturational state, or that *IGHV3-33* usage does not necessarily determine potent neutralization. Notwithstanding, *IGHV3-33*antibodies tended to present lower lethal EC50s, and higher inhibitory and ADCP activities than those encoded by other gene segments, without reaching statistical significance, and induced a longer CSPR phenotype (**fig. S8A-D**). Interestingly, we identified three non-*IGHV3-33* antibodies (ISG4.171 and ISG8.151 [*IGHV1-2*]; ISG8.143 [*IGHV3-30-3*]) that presented lethal EC_50_ below 10 µg/ml, and moderate to strong inhibitory activities [57-98%] (**fig. S8E**), indicating that RTS,S also selects alternative, non-*IGHV3-33* B-cell antibodies with potent antiparasitic properties.

### Potent antiparasitic CRR antibodies frequently recognize short NANP repetitions and cross-react with different *Pf*CSP motifs

The repetitive and disordered structure of the CRR of *Pf*CSP confers multivalency and paratope adaptability, which may contribute to antibody binding promiscuity (*60*). Consequently, high-affinity CRR antibodies tend to cross-react with other *Pf*CSP motifs (*26*, *31*). To define the binding profiles associated with antiparasitic properties, we performed ELISA titration assays using a set of peptides covering the junctional KQPA and NPDP motifs (Jt_1_, KQPADGNPDPNANPN and Jt_2_, GNPDPNANPNVDPNA, respectively), a short minor repeat sequence containing the NVDP motif (M_1_, NANPNVDPNANPNV), the full minor repeat region (M_2_, NANPNVDPNANPNVDPNANPNVDP), and increased numbers of NANP repetitions (NP[NANP]_3-6_) (**Fig. 3A**).

**Fig. 3.**
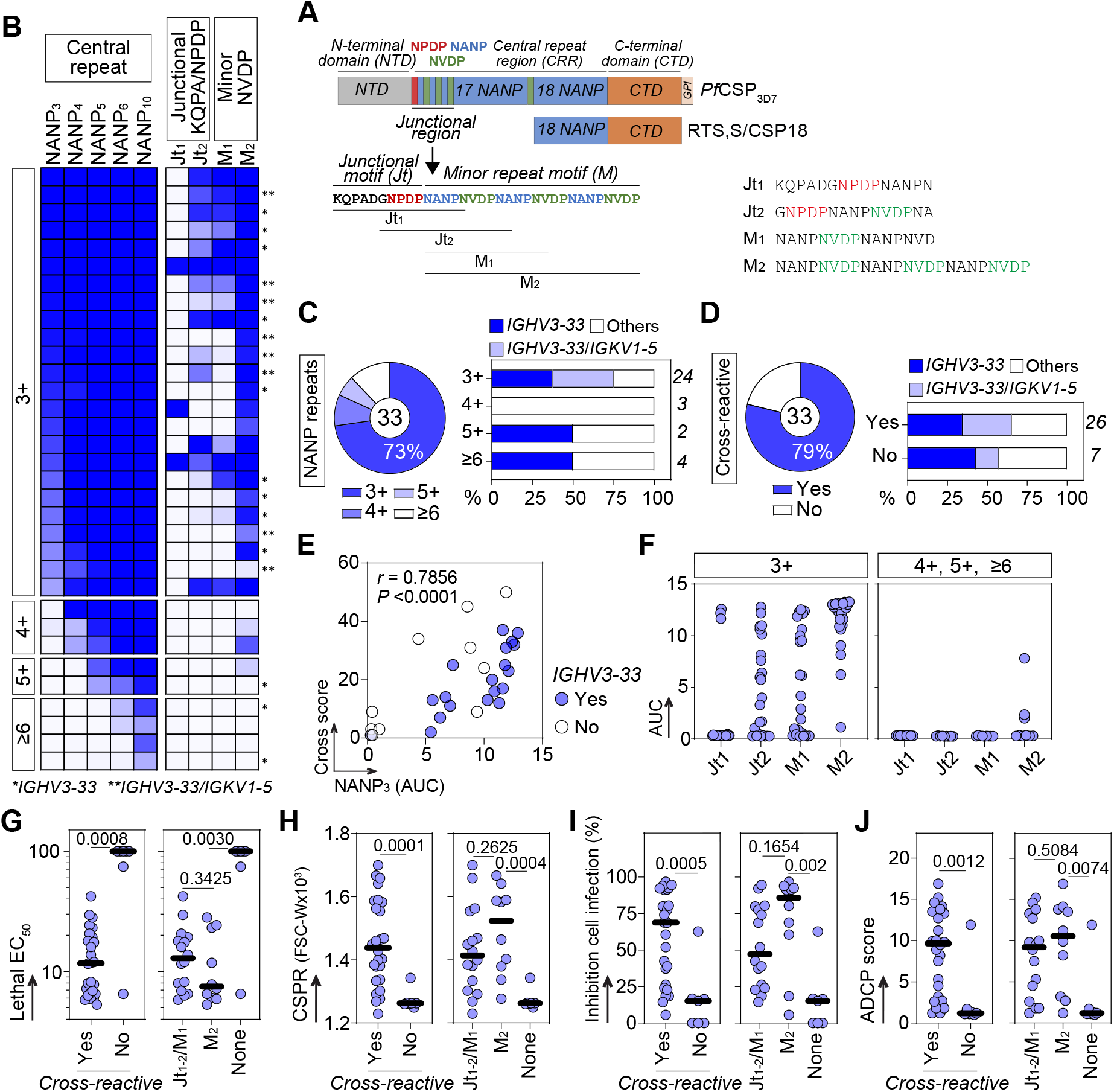
Reactivity profiles of CRR antibodies and *in vitro* antiparasitic properties. (**A**) Sequence diagram of the 3D7 *Pf*CSP protein. The detailed amino acid sequence of the junctional region is shown, with amino acid sequences of peptides Jt_1_, Jt_2_, M_1_ and M_2_ indicated. (**B**) Heatmap showing the reactivity of NANP_10_ antibodies against peptides comprising incremental NANP tetrapeptide repetitions (NANP_3-10_), the junctional KQPA/NPDP motifs (Jt_1_-Jt_2_) and the minor NVDP region of *Pf*CSP (M_1_-M_2_). The color-coded cells are expressed according to the ELISA area under the curve (AUC) values (see fig. S9B an table S2), with darker blue representing higher reactivity. The antibodies are grouped according with the recognition of at least three (3+), four (4+), five (5+) or six or more (≥6) NANP repetitions. The asterisks depict *IGHV3-33*-encoded antibodies paired (**) or not (*) with the *IGKV1-5* gene segment. (**C**) Graphs showing the proportion of NANP_10_ antibodies recognizing different NANP repetitions (pie chart, left), categorized by the Ig gene family used (bar graph, right). (**D**) Same as in (B) but for the proportion of cross-reactive antibodies. (**E**) Correlation plot showing the NANP_3_ reactivity versus the cross-reactive score. Color code depicts *IGHV3-33* usage. The two-sided Spearman rho correlation coefficient and corresponding *p* value are shown. (**F**) Dot plots showing the ELISA AUC values of NANP_10_ antibodies against peptides Jt_1-2_ and M_1-2_, categorized by their reactivity to different NANP repetitions. (**G**) Dot plots showing the lethal EC_50_ values of NANP_10_ antibodies categorized by cross-reactivity (left) or segregated by their cross-reactivity to KQPA/NPDP/NVDP short peptides (Jt_1-2_ and M_1_), the full NVDP minor repeat (M_2_) or none. (**H**) Same as in (G), but for the CSPR phenotype, measured at 10 µg/ml. (**I**) Same as in (G), but for the percentage of inhibition of cell infection, measured at 3 µg/ml. (**J**) Same as in (G), but for the ADCP score, measured at 0.1 µg/ml. Groups were compared with the two- tailed Mann- Whitney test. *p* values are indicated.

A large subset of 24 out of 33 CRR antibodies recognized a short NANP_3_ peptide, of which 18 are encoded by *IGHV3-33*, paired with *IGKV1-5* or other gene segments (**Fig. 3B-C and fig. S9A and B**). Analysis of the reactivity to Jt_1-2_ and M_1-2_ peptides revealed that 26 out of the 33 antibodies cross-recognize the full minor repeat region (M_2_), of which 16 exhibited distinct binding profiles predominantly to Jt_2_ and M_1_, indicating a preferential cross-recognition of NVDP and NPDP containing motifs (**Fig. 3B and D and fig. S9A, C and D**). Conversely, few antibodies recognize the Jt_1_ KQPA peptide (**Fig. 3B and fig. S9A and C**), as similarly reported in adults immunized with live SPZs (*26*). Both cross-reactive and non-cross-reactive groups included antibodies encoded by *IGHV3-33* **(Fig. 3D)**, supporting diverse antibody binding profiles associated to this gene segment (*26*, *60*). *IGHV3-33* cross-binders usually presented NVDP or NPDP binding profiles, whereas the three antibodies recognizing the Jt_1_ KQPA peptide were encoded by *IGHV3*-7 or *IGHV4-61* genes (**fig. S9D and E**). Interestingly, binding signals to NANP_3_ were strongly associated with antibody cross-reactivity scores (defined by the sum of the AUC values to jt_1_, jt_2_, M_1_ and M_2_ peptides), and the vast majority of cross-binders recognized at least three NANP repetitions (**Fig. 3E and F)**, suggesting that narrowing the response towards the NANP repeat core promotes the cross-recognition of similar motifs within the *Pf*CSP sequence.

Importantly, cross-binders presented significantly superior Fab- (lethality, CSPR, inhibition) and Fc- (ADCP) dependent antiparasitic properties *in vitro*, with antibodies exclusively recognizing the NVDP minor repeat (M2) exhibiting a non-significant trend towards more potent signals over those showing a KQPA/NPDP junctional binding profile **(Fig 3G-J)**. We also identified a NANP_3_ negative, non-cross-reactive antibody (ISG8.143), encoded by *IGHV3-30-3/IGKV1-5* gene segments, exhibiting a strong lethality and moderate inhibitory activities. We conclude that the majority of CRR antibodies strongly recognize short NANP repeats, facilitating the cross-recognition of structurally similar *Pf*CSP motifs not included in the RTS,S vaccine immunogen, leading to potent antiparasitic properties likely due to a higher apparent affinity binding and opsonization load.

### Near-germline CRR antibodies induce sterile protection *in vivo*

We evaluated the *in vivo* protective efficacy of CRR antibodies using C57BL/6 mice and a transgenic rodent-infective GFP-expressing PbPf SPZ (*53*). We selected four *IGHV3-33* CRR antibodies exhibiting a range of potent antiparasitic properties and diverse *Pf*CSP binding profiles (**fig S9F-G**). ISG6.149 was the most potent antibody of the whole collection, with a lethal EC_50_ of 5.2 µg/ml, and 92% invasion inhibition, followed by ISG5.143 and ISG5.163 (lethal EC_50_, 6.5 and 8 µg/ml; and 91% and 80% inhibition, respectively). ISG5.144 presented a slightly lower cytotoxicity and inhibition than its counterparts (12 µg/ml lethal EC_50_; and 69% inhibition). Of note, the four antibodies presented a near-GL configuration, carrying on average 5 [2-9] and 3 [1-6] nucleotide mutations in the HC and LC, respectively (**table S2**). Despite being poorly mutated, the antibodies were neither polyreactive nor bound to self-antigens, as determined by ELISA as well as by indirect immunofluorescence on Hep-2 cells, except ISG5.144 which weakly recognized dsDNA (**Fig. S10**). The four antibodies cross-recognize the full minor repeat region (peptide M2), whereas ISG5.144 and ISG5.163 also bind to the short NVDP/NPDP motifs (peptides Jt2 and M1, **fig. S9G**). As anticipated, all four antibodies bound to *Pf*CSP on the SPZ surface and induced an elongated CSPR phenotype with distinct patterns as quantified by flow cytometry and observed by confocal microscopy (**Fig 4A and B**). The three antibodies coming from the same donor (ISG5.143, ISG5.144 and ISG5.163) elicited a medium-long CSPR already at 10 µg/ml, while ISG6.149 induced a much shorter CSPR at the same concentration (**Fig. 4A**). Since the CSPR is dependent on parasite motility and viability, this difference is likely due to a higher affinity (**table S4**) and cytotoxic activity of ISG6.149 (**Fig. 4A and fig. S9F)**.

**Fig. 4.**
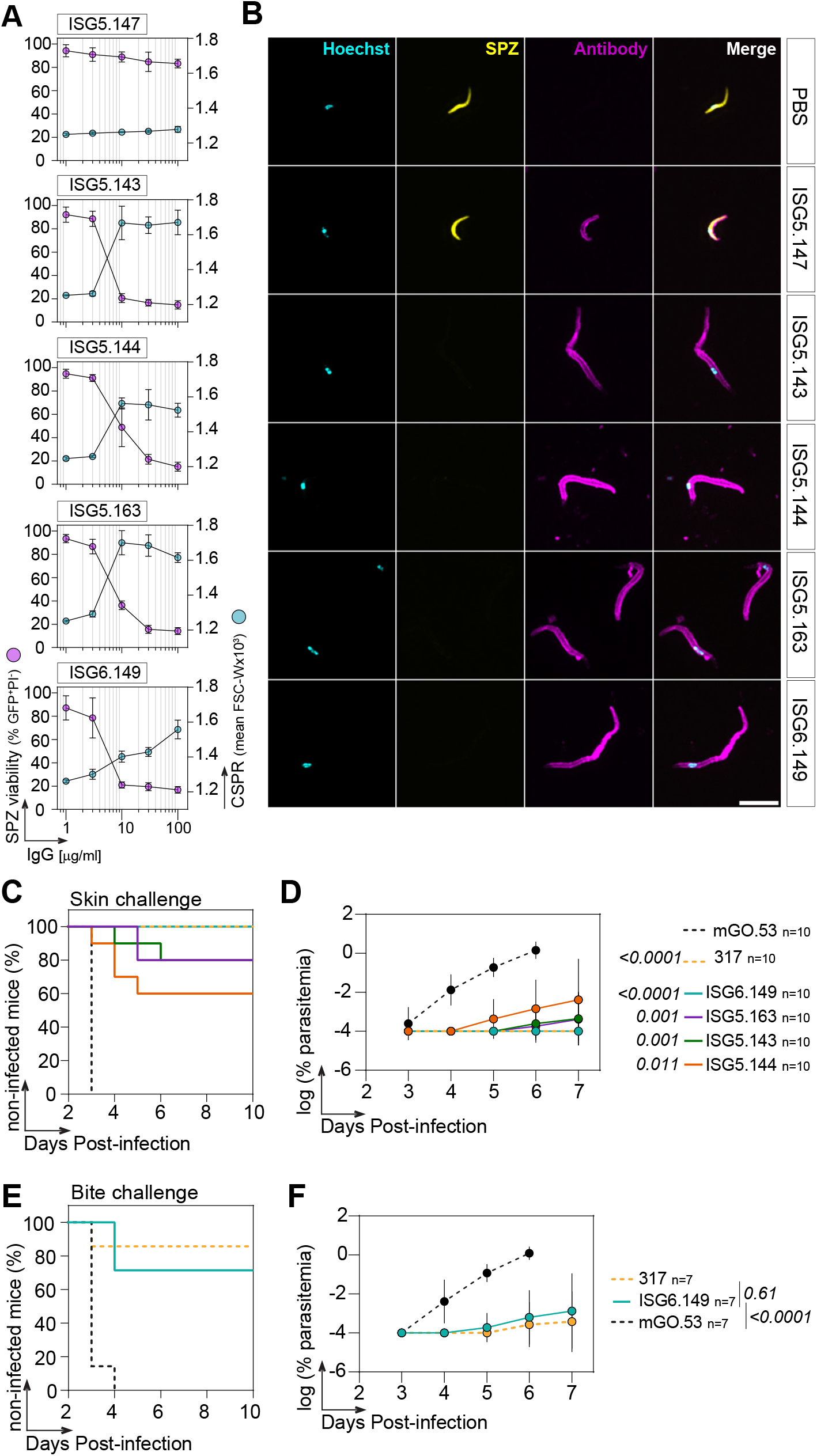
*In vivo* protective activity of CRR antibodies. (**A**) Titration graphs showing the SPZ viability (purple, left) and CSPR phenotype (cyan, right) after incubation of PbPf SPZ with the indicated antibody concentrations for 45 min at 37°C (n=3 independent experiments, data are represented as mean ± SEM. Error bars are indicated). (**B**) Representative confocal microscopy pictures showing distinct CSPR patterns induced in PbPf SPZ after incubation with 100 µg/ml of the indicated antibody for 45 min at 37°C. *Pf*CSP (pink) was revealed by incubating SPZ with Alexa Fluor™ 647-conjugated anti-human IgG secondary antibody. The scale white bar represents 10 µm. (**C**) Survival curves of mice challenged intradermally (footpad) with 5,000 PbPf SPZ, 24 h after passive immunization with 150 µg of the indicated antibodies (solid lines), a protective antibody control (317, dashed yellow line) and an isotype negative control (mGO.53, dashed black line). *p*-values were determined by comparing mGO.53 to the other antibodies using the Holm-Šídák test. (**D**) Graph showing the levels of parasitemia during the follow up days after PbPf SPZ challenge, for the indicated antibodies. (**E**) Same as in (C), but showing mice challenged via infected mosquito bites. *p*-values were determined by comparing mGO.53 and 317 to ISG6.149 using the Holm-Šídák test. (**F**) Same as in (D) but for mice challenged via infected mosquito bite.

Passively immunized mice were challenged by inoculation of SPZ in the skin (*51*). One day after intravenous transfer of a single dose of 150 µg of antibodies, mice were microinjected with 5000 PbPf SPZs in the footpad. For comparison, we included the protective benchmarked 317 antibody (*34*), and the isotype control mGO.53 (*61*). ISG5.143, ISG5.144 and ISG5.163 presented significant protective levels (80%, 60% and 80%, of non-infected mice, respectively), whereas 100% of mice treated with ISG6.149 remained parasitemia-free up to 10 days after infection (**Fig. 4C**). Notably, the mice treated with ISG5.143, ISG5.144 and ISG5.163 that became infected presented significantly lower parasitemia than those administered with mGO.53 (**Fig. 4D**). These results strengthen previous associations between *in vitro* and *in vivo* parasite neutralization (*53*), with ISG6.149 and ISG5.144 leading to 100% and 60% sterile protection, and displaying the highest and lowest cytotoxicity and invasion inhibition, respectively (**fig. S9F**).

To substantiate the highly protective capacity of ISG6.149 in a model resembling natural transmission, mice were challenged with bites of infected mosquitoes 24 h after receiving a single dose of 150 µg of the antibody. Passive transfer of ISG6.149 resulted in 70% of mice remaining free of parasitemia after 10 days post infection, showing non-significant differences with those treated with 317 (**Fig. 4E**). The percentage of parasitemia of ISG6.149-treated mice was significantly lower to those injected with mGO.53 (**Fig. 4F**). Thus, RTS,S can elicit protective antibodies in infants, with ISG6.149, despite its low mutation rate, exhibiting a strong *in vivo* efficacy against SPZ infection.

### Protective CRR antibodies utilize an extensive network of inter-Fab homotypic contacts

To define the molecular details of *Pf*CSP recognition by protective CRR antibodies, we determined the high-resolution structures of ISG5.143, ISG5.144, ISG5.163 and ISG6.149 in complex with the rsCSP protein (*62*), using cryo-electron microscopy (cryo-EM) (**Fig. 5A, fig. S11 and S12, and table S5**). Epitope-paratope interactions revealed a highly conserved binding pose for the four antibodies, recognizing the CRR in a classic type 1 β-turn conformation (*62*), involving *IGHV3-33* GL residues S31 and Y32 on CDRH1, as well as W52, Y52A and Y58 on the CDRH2 (**Fig. 5A**). The anchor point between the strictly conserved W52 and proline 6 (P6) of the NPNA core was identified, which is a distinctive feature of *IGHV3-33* CRR antibodies (*45*, *63*, *64*). LC-mediated binding occurred through the CDRL3 loops using predominantly conserved residues, N92 (Y92 in ISG5.143), S93 (N93 in ISG5.163), Y94 (L94 in ISG5.144) and W95 (F95 in ISG6.149) (**Fig. 5A**). All antibodies presented similar average buried surface areas (BSA) between the HC and LC and CRR, both per residue and across all residues (BSA: ISG5.143 = 1200 Å^2^; ISG 5.144 = 946 Å^2^; ISG 5.163 = 1116 Å^2^; ISG 6.149 = 1006 Å^2^), supporting a conserved binding mode (**Fig. 5B**). Most of the epitope (N_1_PNANPNA_8_) is sandwiched between the three CDRH loops, explaining HC interaction with most of the residues, in contrast to the LC which mostly interacts with Pro_2_ and Asn_3_ residues similarly between the four antibodies (**Fig. 5B**).

**Fig. 5.**
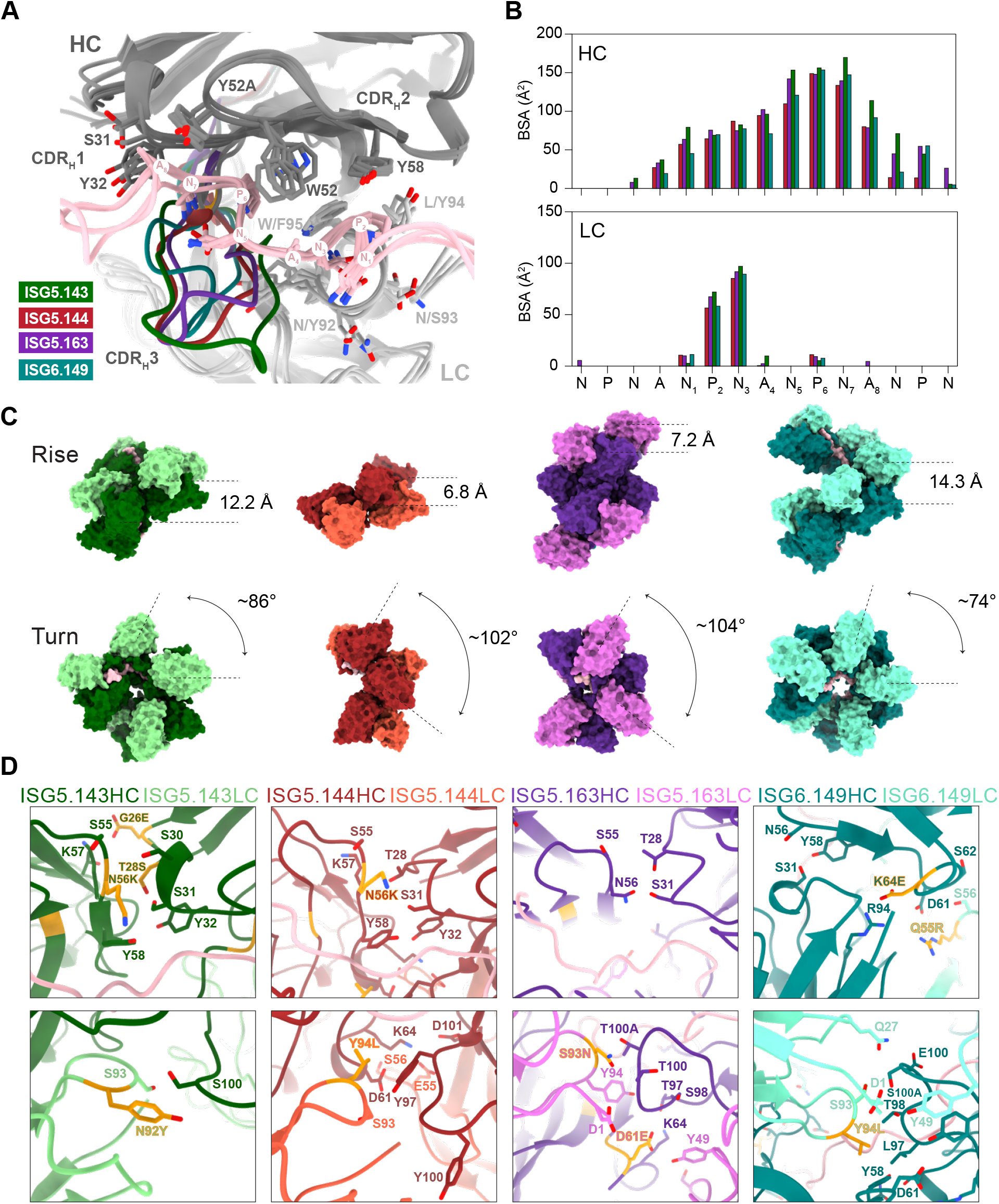
CryoEM structural analyses of ISG.5.143, ISG5.144, ISG5.163 and ISG6.149 antibodies. (**A**) Ribbon representation of the overlap of a single Fab heavy chain (HC, dark grey) and light chain (LC, light grey) bound to rsCSP (pink) for the four NANP_10_ antibodies, with key interacting residues shown as sticks. The NPNANPNA paratope residues are labelled N1 to A8. The CDRH3 of each Fab is highlighted with distinctive colors: ISG.5.143 (green), ISG5.144 (beige), ISG5.163 (purple), and ISG6.149 (teal). (**B**) Bar graph showing the buried surface area (BSA) contributions for the HC (top) and LC (bottom) by each residue within the full epitope of the antibodies, color coded as in (A). (**C**) Surface molecular representation of rsCSP in complex with the four Fabs color coded as in (A). Rise (top) and turn (bottom) helical parameters are indicated for each Fab. (**D**) Structural details of key homotypic interfaces for each Fab, between HC (dark colors, top) and HC-LC (dark and light colors, respectively, bottom). Residues shown as sticks are within 4Å of a residue in the adjacent Fab. Somatically mutated residues are indicated in orange. The axis positions and CDR loops definition follow the Kabat numbering system.

All antibodies formed spiral-like structures, stabilized through homotypic contacts between adjacent Fab molecules, as similarly reported for potent *IGHV3-33* antibodies from adults (*45*, *47*, *65*). The four complexes formed right-handed spirals, with inter-Fab rises ranging from 6.8 Å to 14.3 Å, whereas turn angles ranged from 74° to 104° (**Fig 5C and table S6**). ISG6.149 showed the lowest turn angle (74°) and highest rise (14.3Å), consistent with a narrow Fab packing around *Pf*CSP, while ISG5.144 presents a much gentler rise (6.8 Å) but broader span between Fabs (102°). Interestingly, ISG5.163 formed two stacked spirals, resembling that previously observed for 227 CRR antibody (*47*), but presenting a larger rise, instead of a flat disk-like shape adopted by the latter (**fig. S13**).

Inter-Fab homotypic contacts are mediated by GL-encoded and somatically mutated residues predominantly located at the HC-HC interface, involving the six CDRH loops, whereas the main LC contact points are located at the end of the CDRL2 and CDRL3 (**Fig. 5D and fig S14**). Additional contacts on the stacked spirals of ISG5.163 are located around CDRH1 and CDRH2, and mediated by GL residues (**fig. S13**). The four antibodies presented at least one amino acid replacement at the HC-HC interface, frequently introducing a charged residue such as Asp or Lys. Consistent with a higher mutational load (**table S2**), antibody ISG5.143 developed a rich network of salt bridges between adjacent HC and introduced LC N92Y to the HC-LC interface. In addition, the homotypic interface includes a mutated serine at HC position T28 (T28S), which is a recurring homotypic contact in *IGHV3-33* antibodies (*47*), as for ISG5.144 and ISG5.163 (**Fig. 5D**). Of note, ISG6.149 retained a full GL sequence in most of the homotypic interface, with HC K64E and LC Q55R replacements mediating contacts, as similarly observed for ISG5.163, which only presented amino acid replacements HC D61E and LC S93N at the HC-LC interface. Finally, ISG5.144 presents HC N56K and LC Y94L replacements mediating homotypic contacts, the latter being also observed in ISG6.149 (**Fig. 5D**), suggesting that selection of a less bulky residue than tyrosine at this position may contribute to the tight packing of homotypic structures. Thus, the four antibodies utilize an extensive network of homotypic contacts involving both GL-encoded residues and amino acid replacements, presumably selected by affinity maturation, to stabilize an extended helical spiral structure bound to *Pf*CSP.

### Germline-reverted variants of protective *IGHV3-33* antibodies similarly recognize the CRR as their mutated counterparts

Clonal selection of high affinity *IGHV3-33* B-cell precursors can determine the evolution of potent antibody responses to *Pf*CSP in adults (*27*). To investigate the structural and functional correlates of affinity maturation of protective antibodies, we sought to determine the role of somatic mutations by expressing the unmutated predicted GL versions of the HC (GL_HC), LC (GL_LC), or both (GL) (**fig. S15**). Reverted variants of ISG5.144, ISG5.163 and ISG6.149 exhibited strong binding to CSP18 by ELISA, whereas ISG5.143 completely lost antigen binding upon HC reversion (**Fig. 6A**). Co-crystal structures of a repeat peptide bound to ISG5.144GL, ISG5.163GL and ISG6.149GL (**tables S7 and S8**) revealed the recognition of a similar conformation, retaining the same hydrogen-bonding network, to that of the mutated counterparts (**Fig. 6B and fig. S16**), rationalizing the use of GL-encoded residues to mediate antigen contacts. Surprisingly, unlike the other antibodies, ISG6.149GL forms head-to-head inter-Fab contacts in crystals (**Fig. 6C**), which likely restricts the accommodation of a continuous, long-range helical spiral structure, as observed in the mutated form. Head-to-head interactions have been previously reported for *IGHV3-33* 1210 and *IGHV3-23* 1410 antibodies; however, the homotypic interfaces are asymmetric and are mediated in part by somatically mutated residues (*45*). More recently, symmetric and predominantly GL-encoded Fab-Fab interactions were described for *IGHV3-49* antibodies 7160 and 399 (*66*). In ISG6.149GL, the Fab-Fab interface is likewise symmetric, with CDRH1 of one Fab contacting CDRH3 and CDRL2 of the adjacent Fab, and the reciprocal contacts formed by neighboring Fab (**Fig. 6D**). Specifically, G26 and F27 in CDRH1 forms hydrogen bonds with S56 in CDRL2, while T28 forms a hydrogen bond with Y100_D_ in CDRH3 of the neighboring Fab, forming a reciprocal and symmetric Fab-Fab interface (**Fig. 6D**). To further evaluate the ability of ISG6.149 GL variants to form homotypic contacts and spirals, we solved the cryo-EM structures of rsCSP in complex with the three GL-reverted versions of the antibody (**fig. S12 and S17, and table S6**). Interestingly, ISG6.149GL_HC and ISG6.149GL_LC preserved the homotypic spiral conformation, although ISG6.149GL_HC was noticeably shorter (**fig. S17**). In addition, both variants overlap with the mature spiral with a root mean square deviation (RMSD) of 0.555 Å and 0.434 Å, respectively, and conserve most of their GL-encoded homotypic contacts. In striking contrast, the solved structure of the fully reverted ISG6.149GL complex did not form helical spirals, exhibiting a disordered structure bound to rsCSP resembling non-homotypic antibodies (**fig. S17**). These structures indicate that, even for antibodies showing a near-GL configuration, a few somatic mutations are required to stabilize the long-range helical spiral complexes.

**Fig. 6.**
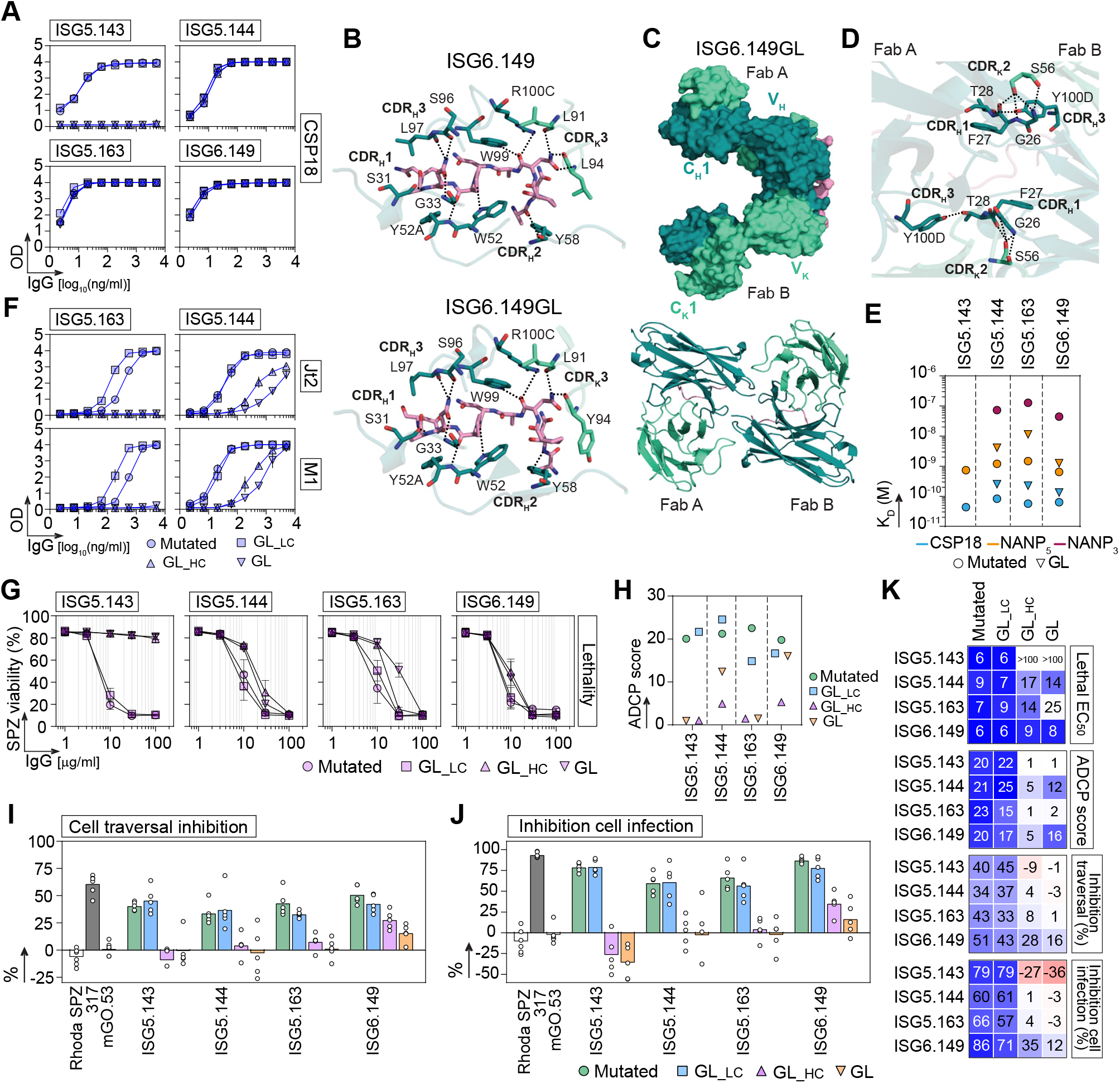
Structural and functional analyses of germline-reverted ISG.5.143, ISG5.144, ISG5.163 and ISG6.149 antibodies. (**A**) ELISA graphs comparing the reactivity of the mutated (circles), and light chain (GL_LC, square), heavy chain (GL_HC, up triangle) and fully (GL, down triangle) germline-reverted ISG.5.143, ISG5.144, ISG5.163 and ISG6.149 antibodies against the CSP18 protein. (**B**) Comparison of the crystal structures of ISG6.149 (top) and ISG6.149GL (bottom) Fabs bound to NPNA_3_ epitope. CDRL3 L94 amino acid replacement is reverted to Y94 in ISG6.149GL Fab, while preserving peptide engagement. (**C**) Surface (top) and cartoon (bottom) representations of head-to-head homotypic interactions between two ISG6.149GL Fab molecules. Fab A and Fab B are colored deep teal (HC) and cyan-green (LC). The NPNA_3_ peptide is shown as a pink cartoon. (**D**) Close-up view of the structural details of key homotypic Fab-Fab interfaces of ISG6.149GL, highlighting the symmetric inter-Fab contacts. Interacting residues are shown as sticks, and hydrogen bonds are indicated by black dashed lines. (**E**) Graph comparing the apparent affinity (*K_D_*) of the four mutated (circles) and fully GL-reverted (down triangle) antibodies against CSP18 (turquoise), NANP_5_ (orange) and NANP_3_ (red). Data represents the mean of two independent runs. (**F**) Same as in (A), but for ISG5.144 and ISG5.163 antibodies against the NPDP (Jt_2_) and short NVDP (M_1_) peptides. (**G**) Titration graphs showing SPZ viability after incubation of PbPf SPZ with the indicated antibody concentrations for 45 min at 37°C (n=3 independent experiments, data are represented as mean ± SEM. Error bars are indicated). Antibodies are coded as in (A). (**H**) Graph comparing the antibody-dependent cellular phagocytosis (ADCP) scores of mutated and GL-reverted antibodies. (**I**) Bar graph comparing the percentage values of cell traversal inhibition of mutated and GL-reverted antibodies. Average of four independent experiments is shown. (**J**) Same as in (I), but for the cell infection inhibition. (**K**) Heatmap summarizing the antiparasitic properties of the mutated and GL-reverted antibodies as determined in (G-J). Darker color indicates lower lethal EC_50_, or higher ADCP scores and inhibition of traversal and invasion.

### Germline-reverted CRR antibodies present high affinity and cytotoxic activities

We next determined the affinity constants of both fully-reverted and mutated antibodies to CSP18, NANP_5_ and NANP_3_ by SPR (**fig. S18 and table S4**). Remarkably, ISG5.144GL, ISG5.163GL and ISG6.149GL present picomolar (*K_D_*=134-252 pM) and nanomolar (*K_D_*=1-11 nM) affinities for CSP18 and NANP_5_, respectively, with *K_D_* values ranging from ∼2 to 8-fold higher than the mutated counterparts, mainly driven by increased dissociation rates (**Fig. 6E, fig. S18 and table S4**). Interestingly, ISG6.149 was minimally affected by GL-reversion, showing a *K_D_* increase of only 1.9- and 2.1-fold for CSP18 and NANP_5_, respectively (**table S4)**, supporting that GL-encoded features strongly contribute to binding affinity. Unlike mutated antibodies, inaccurate fitting of the kinetic curves of GL-reverted variants precluded the obtention of affinity constants for NANP_3_, showing a fast, weak binding. Combined with the observation that somatic mutations either on HC or LC seem to be sufficient to initiate helical spiral formation, this suggests that affinity maturation improves the recognition of short NANP motifs, through complex stabilization facilitated by tighter Fab-Fab interactions. Consistently, ISG5.163GL_HC and ISG5.144GL_HC variants decreased or completely abrogated the recognition of NPDP (Jt_2_) and short NVDP (M_1_) peptides, respectively (**Fig. 6F)**, supporting that HC somatic mutations facilitate cross-reactivity to the junctional region (*30*).

We next evaluated the *in vitro* Fab-dependent lethality, reasoning that the strong affinity binding of GL-reverted antibodies would still impact SPZ motility, ultimately provoking parasite death (*11*). Interestingly, ISG5.144GL, ISG5.163GL and ISG6.149GL presented potent or moderate lethality values (average EC_50_=16 [8-25] µg/ml) (**Fig. 6G and K**). As expected, mutated antibodies presented improved parasite killing, with 1.5 to 3.3-fold lower EC_50_ than their reverted counterparts, mostly contributed by HC amino acid replacements (**Fig. 6G and K**). Notably, ISG6.149GL preserved a potent EC_50_ of 8 µg/ml, suggesting that ISG6.149 originated from a highly cytotoxic B-cell precursor. In line with the role of HC somatic mutations, ISG5.143GL and ISG5.143GL_HC completely lacked cytotoxic activities (**Fig. 6G and K**). Similarly, both ISG5.143 and ISG5.163 lost their ADCP capacity upon HC reversion, in contrast to ISG5.144GL and ISG6.149GL which preserved moderate scores (**Fig. 6H and K**). Surprisingly, ISG5.144GL_HC and ISG6.149GL_HC presented decreased ADCP scores as compared with the fully reverted versions (**Fig. 6H and K**), suggesting that LC amino acid replacements alone may interfere with parasite opsonization. Collectively, our findings point to the selection of cytotoxic, high-affinity precursors by RTS,S immunization. Affinity maturation incorporates amino acid replacements that alter the steric and structural properties of the antibodies, allowing the formation of homotypic helical assemblies, in a fine-tuning effect resulting in increased reactivity breadth and SPZ lethality.

### Somatic mutations drive potent inhibition of SPZ cell invasion and traversal

Protective *Pf*CSP antibodies usually present high affinity binding, but not necessarily potent cytotoxicity, suggesting that additional properties may determine protective efficacy (*53*, *67*). To decouple Fab-dependent cytotoxicity from other antiparasitic properties, we next evaluated the capacity of GL-reverted antibodies to inhibit SPZ cell traversal and invasion under non-cytotoxic concentrations. Interestingly, GL-reversion of the HC substantially decreased the ability to hinder inhibitory activities, with only ISG6.149 preserving residual levels of inhibition (**Fig. 6I-K**). Thus, the molecular and structural changes driven by somatic mutations lead to efficient inhibitory activities, which may ultimately contribute to protective efficacy.

## Discussion

Understanding protective immunity in vulnerable populations is imperative to guide the development of next-generation vaccines. While *Pf*CSP B-cell responses have been investigated in adults (*25–27*, *31*, *33*), the molecular and functional attributes of vaccine-induced antibodies in early childhood have remained largely elusive. We addressed this gap by characterizing a collection of *Pf*CSP human antibodies from Mozambican infants, defining the hallmarks of potent B-cell responses to malaria during the early stages of life. RTS,S elicited protective antibodies in a near-GL configuration, extending proof of principle that efficient *Pf*CSP B-cell responses rapidly develop in African children. Most important as a conceptual advance, RTS,S selected high-affinity, cytotoxic *Pf*CSP B-cell antibody precursors, which are likely present in the infant naïve repertoire. Our findings indicate that the partially developed B-cell diversification mechanisms associated with early childhood (*20*) do not preclude the elicitation of protective antibodies, supporting ongoing vaccination programmes in Africa, and immunogen development for young children.

An effective vaccine with broad coverage should elicit immune responses with shared elements among population groups (*68*). Most of the CRR antibodies isolated here were encoded by *IGHV3-33*, paired with an 8-amino acid-long CDRL3 preferentially using the *IGKV1-5* gene segment, which are likewise highly selected in adults due to an optimal architecture for high affinity binding (*27*, *31*, *33*, *45*). Importantly, we identified selected somatic mutations among different CRR antibodies, frequently observed in malaria-naïve American or European adults immunized with RTS,S or live SPZs (i.e., H.V50I, H.N56K, H.K64E, K.Y94L, K.S93N) (*45*, *47*), respectively. These findings support reproducible antigen-driven selection traits across different age groups and ethnicities, suggesting that many of the structural solutions for *Pf*CSP recognition observed in adults are available in infants.

High-affinity antibodies can be induced early without affinity maturation (*69*). A key finding of the current study is that protective CRR antibodies exhibited minimal SHM loads, with antigen contacts mediated by GL-encoded residues only. This would explain the strong affinity of GL-reverted variants, and reflects classical hapten studies showing that initial high affinity B cells tend to accumulate less mutations than low affinity clones, due to more relaxed positive selection mechanisms because an affinity ceiling has been reached (*70–72*). This is reminiscent to that reported for viruses like RSV (*23*), hepatitis C virus (*73*), cytomegalovirus (*74*) or SARS-CoV-2 (*75*); and bacteria such as *Staphylococcus aureus* (*76*). The data expand the concept of “innate-like scaffolds” recognizing specific structural motifs (*77*), and extend this to a repetitive region of a eukaryotic parasite. Our findings also build upon previous observations in malaria naïve adults inoculated with PfSPZ-CVac (*27*), by showing that clonal selection of high-affinity B-cell precursors orchestrates protective antibody responses. Our discoveries suggest that high affinity B-cell precursors are likely available in the infant naïve repertoire, and that these cells could be expanded by *Pf*CSP-based immunization strategies.

Despite being scarce, somatic mutations were sufficient to shape the structural and functional evolution of protective antibodies. Antigen-driven selection led to an evolutionary pathway towards long-range helical ordering stabilized by homotypic interactions (*45*, *47*, *65*), in a fine-tuning effect that led to increased binding affinity to the NANP core, broader junctional reactivity, and efficient inhibition of parasite cell traversal and invasion. It is likely that the formation of stable antibody-*Pf*CSP helical complexes induces a rigid conformational state that modifies the intrinsically disordered and flexible properties of the CRR (*78*), hindering *Pf*CSP function (*54*, *56*, *79*), ultimately affecting parasite arrest in the liver and hepatocyte infection. While a full understanding on how CRR antibodies target SPZ *in vivo* merits further investigation, our findings suggest that few amino acid replacements differentially modulate distinct mechanisms of action of *Pf*CSP antibodies.

Our structural studies confirmed the wide range of helical spiral *Pf*CSP conformations induced by *IGHV3-33* antibodies (*45*, *47*, *65*, *80*). Importantly, analysis of GL-reverted variants revealed that ISG6.149GL_HC and ISG6.149GL_LC are able to maintain a helical spiral conformation despite reversion of some amino acid residues promoting homotypic contacts. Contrarily, the fully reverted ISG6.149GL is no longer able to form a stable homotypic spiral, although it does present head-to-head inter-Fab contacts in crystals, resembling the structural configuration reported for *IGHV3-33* antibody 1210 (*45*). These findings indicate that ISG6.149-like precursors may be predisposed to form inter-Fab contacts, potentially explaining its superior NANP_5_ affinity and lethality, and subsequent maturation leading to a highly protective antibody. In contrast, GL-reverted ISG5.144 and ISG5.163 lacked inter-Fab contacts but preserved high-affinity binding and moderate lethality, whereas ISG5.143 completely lost antigen binding upon reversion, illustrating that not all *IGHV3-33* B-cell precursors are intrinsically strong binders. Hence, GL features may act in concert with antibody sequence attributes as a result of V(D)J recombination, such as CDRH3 length and amino acid composition. ISG5.143 has an elongated CDRH3 with four charged residues, which may promote steric hindrance or electrostatic repulsion, precluding antigen contacts for the GL variant. Consistently, ISG5.143 required more maturation than ISG5.144, ISG5.163 and ISG6.149, resulting in a rich network of salt bridges mediating inter-Fab homotypic contacts, leading to a highly potent antibody, suggesting different pathways in the infant immune system for developing protective responses.

Although we show that RTS,S can induce protective antibody B-cell responses in infants, different host- and vaccine-dependent factor should be considered for future vaccines aiming to elicit potent and sustained antibody levels in plasma. It has been recently reported that the repetitive structure of the CRR induces a strong BCR cross-linking, leading to fast, short-lived effectors producing high antibody levels, at the expense of generating long-lived memory B cells and plasma cells (*81*, *82*). Moreover, a feedback effect induced by high affinity primary responses may restrict germinal center participation of naïve B cells upon recall (*83*, *84*). These factors could explain both the rapid decay in CRR antibody levels after primary vaccination and the weak responses observed after boosting in RTS,S vaccine trials (*14*, *19*, *43*). Low valency antigens tend to be more stringent by increasing the B-cell activation threshold, selectively activating strong binding precursors (*85*). Hence, priority should be given to CRR shortened versions capable of engaging high-affinity B-cell clones, while bypassing BCR hypercross-linking and apoptosis (*86*). This was supported by promising studies formulating NANP_5_-based immunogens in tobacco mosaic virus (*36*) or *Helicobacter pylori* apoferritin-based nanoparticles (*35*), which induced high-affinity, long-lived antibody responses in preclinical models. Our analyses of GL-reverted antibodies revealed that ISG5.144GL, ISG5.163GL, and ISG6.149GL recognize NANP_5_ with nanomolar affinity, indicating that naïve B-cells could be efficiently expanded by shortened CRR immunogens in humans, further supporting those vaccine strategies.

Despite a high binding strength, CTD antibodies exhibited poor *in vitro* inhibition of hepatocyte invasion and a limited ability to kill SPZ, consistent with the inaccessibility of this region on the parasite surface, providing further evidence of weak neutralizing properties (*28*, *54*, *87*). Recent studies reported that CTD plasma antibodies from RTS,S vaccinees can mediate complement deposition and Fc-dependent effector functions, which might contribute to parasite inhibition (*88*, *89*), although similar experiments using live SPZs remain to be performed. RTS,S primary vaccination led to increased and sustained plasma antibody levels and better booster responses to CTD, which have recently been associated with lower clinical malaria incidence (*43*). However, although CTD antibodies could be a marker of the overall strength of antiparasitic responses or present an adjunctive effect to those targeting the CRR (*90*), the mechanisms underlying their association with reduced malaria risk (*39*, *43*, *91*) remain to be demonstrated. The preferential elicitation of high affinity, non-protective CTD responses could outcompete B-cell selection, and subsequent induction of protective antibodies. This is particularly important in early childhood, where a poorly developed follicular dendritic cell and stromal cell networks (*92–94*), together with the sudden exposure to environmental antigens, tend to restrict long-lived plasma cell survival due to high competition for bone marrow niches (*95*). Future studies will need to demonstrate the CTD antibody functions associated with protection in order to evaluate the benefit of inducing or suppressing the humoral response against this region.

A limitation of this study is that we only analyze *Pf*CSP antibodies from six Mozambican infants, and thus the molecular parameters described may not be representative of the demographic populations from other African regions. In addition, we did not detect *IGHV3-33 Pf*CSP antibodies in donors d4, d8 and d9, pointing to inter-individual variability in the use of Ig gene segments, as previously suggested (*27*, *45*). Although we obtained a low number of CRR antibodies from these donors, this observation is consistent with a recently reported homozygous deletion of the *IGHV3-33* gene in about 40% of individuals in African groups (*96*). Considering that Ig gene variation can compromise humoral responses to pathogens (*97*–*100*), it will be crucial to determine the extent to which this occurs with *Pf*CSP responses upon infection or vaccination. Furthermore, highly potent antibodies using alternative gene families to *IGHV3-33* should be thoroughly investigated. Of note, we identified *IGHV1-2*-encoded ISG4.171 and ISG8.151, elicited in donors d4 and d8, respectively, as a new class of highly cytotoxic, inhibitory antibodies.

This study represents a substantial advance in understanding the molecular mechanisms of protective antibody responses in early life. The identification of near-GL, high-affinity, and protective antibodies, such as ISG6.149, elicited early after primary RTS,S vaccination, will inform next-generation vaccine development, including GL-targeting approaches. Moreover, engineered, long-acting versions of the antibodies described here could provide protective immunity, offering potential developability advantages due to limited somatic mutation loads (*101*). Beyond malaria, our discoveries provide an improved framework for the discovery of protective antibodies to guide next-generation interventions against complex pathogens in early life.

## Methods

### Human samples

Stored plasma and peripheral blood mononuclear cells (PBMC) samples from RTS,S/AS01_E_ vaccinated children were previously obtained from a random subset recruited in Manhiça, Mozambique as part of the immunology MAL067 study ancillary to the phase 3 randomized clinical trial MAL055 (ClinicalTrials.gov identifier: NCT00866619) (*13*). The study protocol of MAL067 was approved by the Ethics Committees from Spain (Comitè Ètic d’Investigació Clínica, Hospital Clínic Research Ethics Committee, Barcelona, 2008/4622) and Mozambique (Comité Nacional de Bioética para Saúde, Manhiça, 332/CNBS/18). Written informed consent was obtained from parents or guardians of all participants before recruitment. PBMC were isolated from venous blood samples by density gradient centrifugation using Ficoll-Paque (Merck), cryopreserved in heat-inactivated fetal bovine serum (HI-FBS) (Thermo Fisher Scientific) with 10 % dimethyl sulfoxide (DMSO) (Merck), and stored in liquid nitrogen.

### Serum IgG purification

Human IgG antibodies were purified from donors’ sera by affinity chromatography using Protein G Sepharose® 4 Fast Flow (GE Healthcare). Purified serum antibodies were dialyzed against PBS using Slide-A-Lyzer® Cassettes (30 K MWCO, Thermo Fisher Scientific). Samples were aliquoted and stored at −80 °C.

### Cell lines

Freestyle 293-F suspension cells (Thermo Fisher Scientific) were maintained in FreeStyle Expression Medium (Thermo Fisher Scientific) and kept under constant shaking at 150 rpm. HepG2 hepatoma cell line (ATCC) was grown in Dulbecco’s modified Eagle’s medium (Thermo Fisher Scientific) with 10% heat inactivated fetal calf sera (FCS, Panbiotech) and non-essential amino acids (Sigma Aldrich). The THP1 promonocytic cell line (ATCC) was maintained in a cell density between 1×10^5^ and 1×10^6^ in completed RPMI 1640 medium (Thermo Fisher Scientific), supplemented with 10% heat-inactivated fetal bovine serum (HI-FBS), Penicillin/Streptomycin, and L-glutamine. All cell lines were cultured at 37°C, with 5% CO2.

### Antigens and antibody controls

Codon-optimized nucleotide fragments encoding recombinant CSP proteins from *Plasmodium falciparum* 3D7 strain (PF3D7_0304600.1) CSPFL_20-373_, CSP18_200-373_ and CTD_273-373_ were synthetized and cloned into pcDNA3.1/Zeo^(+)^ expression vector (Thermo Fisher Scientific). All constructs included Kozak and Ig◻ leader sequences at the N-terminus and AviTag and 10xHis C-terminal tags. Human *Pf*CSP IgG antibody controls are: 311, 317 (*62*), 663 (*25*), 1210 (*45*); CIS43(*30*); L9 (*102*); 1512 and 236 (*103*). Non-*Pf* antibodies include polyreactive ED38 (*104*) and isotype IgG control mGO53 (*61*). Recombinant proteins and antibodies were produced by transient transfection of exponentially growing Freestyle 293-F suspension cells (Thermo Fisher Scientific) using the polyethylenimine (PEI) precipitation method as previously described (*105*). Proteins and antibodies were purified from culture supernatants by high-performance chromatography using the Ni Sepharose Excel Resin or Protein G Sepharose 4 Fast Flow (GE Healthcare), respectively, according to the manufacturer’s instructions, dialyzed against PBS using Slide-A-Lyzer dialysis cassettes (Thermo Fisher Scientific), quantified using NanoDrop One instrument (Thermo Fisher Scientific), and controlled for purity by SDS– polyacrylamide gel electrophoresis (PAGE) using NuPAGE 3 to 12% Bis-Tris gels (Life Technologies) as previously described (*105*). AviTagged CSP18 was subsequently purified by size exclusion chromatography using a Superdex 75 increase 10/300 GL column on an ÄKTA pure 25 system (GE Healthcare), and biotinylated using the Enzymatic Protein Biotinylation Kit (Sigma-Aldrich). Peptides NANP_3-6_, Jt_1_, Jt_2_, M_1_ and M_2_ were synthetized by Genscript. The monoclonal antibody 3D11 was obtained through BEI Resources, NIAID, NIH: Hybridoma 3D11 Anti-Plasmodium berghei 44-kDa Sporozoite Surface Protein, MRA-100, and purified by affinity chromatography using protein G beads, as described above. Recombinant rsCSP protein was purified as previously described (*47*). Briefly, a twin-strep rsCSP construct was expressed in E.coli BL21(DE3) (New England Biolabs) overnight at 18°C, purified with StrepTactinXT resin (IBA Lifesciences) and eluted in TBS. The eluate was then cleared on an S200increase column (Cytiva).

### Single B-cell sorting and expression-cloning of antibodies

Peripheral blood B cells were isolated from PBMCs by magnetic-activated cell sorting using CD19 MicroBeads (Milteny Biotec). CD19^+^ B cells were then stained with LIVE/DEAD fixable dead cell stain kit (Thermo Fisher Scientific) and incubated for 30 min at 4°C with 1 µg of biotinylated CSP18 protein. Cells were washed once with 1% FBS-PBS (FACS buffer), and incubated for 30 min at 4°C with a cocktail of mouse anti-human antibodies: CD19 A700 (HIB19, BD Biosciences, San Jose, CA), CD21 BV421 (B-ly4, BD Biosciences), CD27 PE-CF594 (M-T271, BD Biosciences), IgD APC-H7 (IA6-2), IgM BV605 (G20-127, BD Biosciences), IgG BV786 (G18-145, BD Biosciences), IgA FITC (IS11-8E10, Miltenyi Biotec, Bergisch Gladbach, Germany), and streptavidin R-PE conjugate (Invitrogen, Thermo Fisher Scientific). Stained cells were washed and resuspended in 200 µl of FACS buffer. Single IgG^+^ CSP^+^ B cells were sorted into 96-well plates using a FACS Aria II SORP sorter (Becton Dickinson). Following a lymphocyte and single cell gating, dead cells were excluded. B-cell immunophenotyping analyses were performed using FlowJo software (v10.8.1, FlowJo LLC, Ashland, OR). Single-cell cDNA synthesis using SuperScript IV reverse transcriptase (Thermo Fisher Scientific) followed by nested-PCR amplifications of IgH, Igκ, and Igλ genes, and sequence analysis for Ig gene features were performed as previously described (*106*). Purified digested PCR products were cloned into human Igγ1-, Igκ-, or Igλ-expressing vectors. Recombinant antibodies were produced by transient co-transfection of Freestyle 293-F suspension cells (Thermo Fisher Scientific) using the polyethylenimine (PEI)-precipitation method as previously described (*107*). Recombinant human IgG antibodies were purified by protein G affinity chromatography (Protein G Sepharose 4 Fast Flow, GE Healthcare). Purified antibodies were dialyzed against PBS. The GL variants of ISG5.143, ISG5.144, ISG5.163 and ISG6.149 antibodies were obtained by reverting the V and J segments of the HC and LC to their predicted GL versions, while retaining the V(D)J junction. Synthetic genes were ordered to GeneArt (Thermo Fisher Scientific), cloned into human Igγ1- or Igκ-expressing vectors, and the recombinant antibodies were produced as described above.

### ELISAs

ELISAs to measure binding to recombinant proteins were performed in the ISGlobal Immune response and biomarkers Core Facility as previously described (*108*). High-binding 96-well ELISA plates (Costar, Corning) were coated overnight with 125 ng/well of purified proteins. After washings, plates were blocked 2 hours with 2% bovine serum albumin (BSA), 1mM EDTA, 0.05% Tween 20-PBS (Blocking buffer), washed, and incubated with serial dilutions of purified plasma IgG or monoclonal antibodies in PBS for 2 h. After washings, plates were revealed by incubation for 1 h with goat HRP-conjugated anti-human IgG antibody (Immunology Jackson ImmunoReseach, 0.8 µg/ml final) in blocking buffer, and by adding 100 µl of HRP chromogenic substrate (ABTS solution, Euromedex), after washing steps. After 1 h incubation, optical densities were measured at 405 nm (OD_405nm_). For peptide ELISA, binding of antibodies to *Pf*CSP linear peptides was tested as previously described (*109*). High-binding 96-well ELISA plates (Costar, Corning) were coated overnight with 625 ng/well of peptides. After washings, plates were blocked 2 hours with 3% non-fat milk, 5% sucrose, 1% Tween 20-PBS, washed, and incubated with serial dilutions of purified antibodies in 1% BSA, 0.1% Tween-20-PBS for 2 hours. After washings, plates were revealed as described above. Binding of purified antibodies to biotinylated NANP_3_ peptide was tested by coating plates with 125 ng/well of purified streptavidin (BD Pharmingen). After washings, plates were blocked 2 hours with blocking buffer, washed, and incubated with 10 µg/ml of NANP_3_ for two hours, to subsequently proceed as described above. Experiments were performed using BioTek 405 TS™ microplate washer (Agilent Technologies) and Infinite M Plex microplate absorbance reader (Tecan Männedorf). Antibodies were tested in duplicate in at least two independent experiments, which included mGO53 as negative control, and benchmarked *Pf*CSP antibodies as positive controls. Polyreactivity ELISA was performed as previously described (*110*). High-binding 96-well ELISA plates were coated overnight with 500 ng/well of purified double stranded (ds)-DNA, KLH, Lysozyme, C1q and Thyroglobulin, and 250 ng/well of insulin (Sigma-Aldrich, Saint-Louis, MO), in PBS. After blocking and washing steps, performed as described above, purified IgG antibodies were tested in duplicate at 4 µg/ml and 3 consecutive 1:4 dilutions in PBS. Control antibodies, mGO53 (negative), and ED38 (high positive) were included in each experiment. ELISA binding was developed as described above.

### HEp-2 cell IFA

Binding of human *Pf*CSP and control monoclonal IgG antibodies (mGO.53 and ED38) to HEp-2 cell-expressing autoantigens were analyzed at 100 µg/ml by indirect immunofluorescence assay (IFA) (ANA HEp-2 AeskuSlides®, Aesku.Diagnostics) following the manufacturer’s instructions. IFA sections were examined using the fluorescence microscope IXplore IX73 (Olympus), and pictures were taken at magnification x 40 with 5000 ms-acquisition using CellSens software (Standard 4.4 version, Olympus).

### Surface plasmon resonance

Surface plasmon resonance (SPR)-based technology was performed in the Protein Technologies Unit of the Center of Genomic Regulation (CRG) using a Biacore T200 instrument (Biacore Cytiva) to assess the kinetics of interactions between antibodies and CSP18, NANP_5_ and NANP_3_. Purified antibodies were captured onto a CM5 sensor chip using Human antibody Capture Kit (Cytiva). Binding measurements were performed at 25 °C using HEPES-buffered saline (10 mM HEPES, 150 mM NaCl, 0.05–0.1% Tween-20) as running buffer. For initial screening, IgG antibodies (3 µg ml⁻¹) were captured over flow cell 4 (10 µl min⁻¹, 30 s; 20 s stabilization). For single-cycle kinetics, CSP18 was injected over flow cell 3 and 4 at 1 and 10 nM (60 µl min⁻¹; 100 s association, 600 s dissociation), and data were collected at 10 Hz. Surfaces were regenerated with two 60 s pulses at 50 µl min⁻¹ of 3 M MgCl₂ followed by 30 s stabilization. For refined analysis, antibodies were captured under shorter contact times (12 s; 60 s stabilization), and CSP18 was injected at 0.3–5 nM (90 µl min⁻¹; 100 s association, 1200 s dissociation). For multi-cycle kinetics (MCK), antibodies were captured at 3 µg ml⁻¹ (CSP18, 12 s) or 3 - 5 µg ml⁻¹ (NANP_3_ and NANP_5_, 60 s), at 10 µl min⁻¹, followed by 60 s stabilization. CSP18 (0.013–9 nM) and peptides (1.4 nM–1 µM) were injected at 90 µl min⁻¹ (120 s association, 240 s dissociation). Surfaces were regenerated with three 60 s pulses of 3 M MgCl₂ followed by 60 s stabilization. All data were subtracted and analyzed using Biacore T200 Evaluation Software v3.2.1. Kinetic parameters were obtained by global fitting to a 1:1 Langmuir binding model, with refractive index (RI) set to constant.

### Mice and mosquitoes

For the *in vivo* protection assays, 3-week-old female C57BL/6JRj mice (Janvier Lab) were used. For mosquitoes infection, 3-week-old female RjOrl:Swiss mice (Janvier Lab) were used. Mice were housed in the Center for Production and Infection of Anopheles (CEPIA) animal facility at Institut Pasteur, accredited by the French Ministry of Agriculture for rodents experiments. Work on animals was performed in compliance with the French and European regulations on care and protection of laboratory animals. All experiments were approved by the Ethics Committee #089 and registered under the references MESR01324, dap210057 (APAFIS #32989-2021091516594748 v1) and dap210060 (APAFIS#32422-2021071317049057 v2). *Anopheles stephensi* (SDA 500 strain) mosquitoes were reared at the CEPIA and are maintained at 21°C with 80% humidity and sucrose.

### Parasites and mosquito infection

A transgenic *P.berghei* ANKA strain expressing the full-length of *P. falciparum* 3D7 CSP and GFP under the HSP70 promoter (PbPf) was used (*67*). Swiss mice were infected by intraperitoneal injection of infected red blood cells. When parasitemia, determined by flow cytometry (Cytoflex), reached ≥ 2% and gametocytemia ≥ 0.2%, mice were anesthetized with 90 mg/kg Ketamin and 9 mg/kg Xylazin Rompun 2% and used for mosquito blood feeding, performed by the CEPIA. Mosquitoes were allowed to feed for 20 minutes at 21°C, with mice rotated every 5 min. A naïve feeding was performed one week later and mosquitoes were maintained with sucrose for 3-4 weeks.

### Sporozoites obtention

For *in vitro* assays and skin challenge, sporozoites were isolated from mosquito salivary glands 18-26 days after infectious blood feeding. Mosquitoes were anesthetized on ice and washed with 70% ethanol followed by PBS. Salivary glands were collected in PBS (supplemented with 2X penicillin, streptomycin and neomycin for HepG2 infection experiments), disrupted and filtered through 35µm filters to isolate sporozoites, which were kept on ice until use.

### *In vitro* sporozoite cytotoxicity assay in suspension

Fab-dependent cytotoxicity and Circumsporozoite Precipitation Reaction (CSPR) were assessed by using PbPf sporozoites as described previously (*53*). Around 10,000 PbPf sporozoites were incubated at 37°C for 45 min with different concentrations of *Pf*CSP antibodies in the presence of 10% heat inactivated fetal calf sera (FCS, Panbiotech). Samples were next transferred on ice, incubated for 10 min with 2.5 µg/mL propidium iodide (PI, Invitrogen) and diluted 10 times with cold PBS prior to acquisition on a CytoFLEX S flow cytometer (Beckman Coulter). At least 5,000 events in the sporozoite gate were acquired. Viability was defined as the percentage of GFP^+^PI^−^ sporozoites to the sum of GFP^+^PI^−^, GFP^+^PI^+^ and GFP^−^PI^+^ sporozoites. The CSPR seen in the presence of *Pf*CSP antibodies was determined by the mean intensity of the forward scatter-width signal (FSC-W), which was observed to indicate the length of the sporozoites. Data were analyzed using the CytExpert 2.0 software (Beckman Coulter). For each experiment, 317 and 3D11 antibodies were included as positive and negative controls, respectively. To preclude eventual killing due to acquisition timing, wells without antibodies were acquired before and after those containing the antibodies. Data were expressed as a killing percentage after normalization using 3D11 control.

### CSPR confocal microscopy

PbPf sporozoites were incubated with 100 µg/mL antibodies for 45 min at 37°C in the presence of 10% heat inactivated FCS (Panbiotech). PBS, a cytotoxic antibody (317) and a non-cytotoxic antibody (ISG5.147) were used as controls. Sporozoites were fixed with 2% PFA, washed with PBS by centrifugation (3 min, 500 x g) and blocked overnight at 4°C in 1% BSA/PBS. Samples were then incubated with Alexa Fluor 647-conjugated goat anti-human IgG (1:500) for 1 h at room temperature in the dark, followed by Hoechst 33342 (diluted 1/500) for 10 min. After washing, samples were imaged using a spinning-disk confocal microscope (PerkinElmer UltraView ERS, 100x objective). Images were acquired with Velocity software and analyzed using Fiji.

### HepG2 cell inhibition assays

The ability of CSP18 antibodies to inhibit PbPf SPZ infection of the hepatoma cell line HepG2 was evaluated as described previously (*56*). 40,000 HepG2 cells were plated in a 96 wells plate containing growth media supplemented with 2X penicillin, Streptomycin and Neomycin one day prior infection. PbPf sporozoites were incubated with different concentrations of *Pf*CSP antibodies on the HepG2 cell monolayer at a 1:4 sporozoite-to-cell ratio. After 2 or 48 h at 37°C, 5% CO2 and 10% O2, cells were trypsinized, diluted in 2% FCS/PBS and filtered through 35 µm filters before being analyzed using a CytoFLEX S flow cytometer (Beckman Coulter). The percentage of infected cells was determined based on GFP-positive events within the HepG2 gate, with the same volume acquired for each well. For cell traversal assays, dextran tetramethylrhodamine (ThermoFisher Scientific) was added in the media at 1 mg/mL and cell traversal was quantified as the percentage of wounded cells based on dextran-positive events within the HepG2 gate. Data were expressed as an inhibition percentage after normalization using 3D11 control.

### Opsonic phagocytosis assay (ADCP)

ADCP activity was assessed using fluorescent beads coupled with CSPFL and the human monocyte THP-1 cell line, as described previously (*88*). Amine-modified fluorescent latex beads (Sigma-Aldrich) were washed twice with 400 µl of 0.1% SDS–PBS, centrifuged at 1500g for 5 min, and incubated in 400 µl of 8% glutaraldehyde on a shaker overnight at room temperature. After two washes with 600 µl of 0.1% SDS–PBS, CSP-FL was added at 0.5 mg/ml to the beads and incubated on a shaker for 4 h at room temperature. Coupled beads were subsequently centrifuged at 1500 g for 5 min and incubated with 600 µl of 4% diethanolamine on a shaker for 30 min at room temperature. Beads were then centrifuged at 1500 g for 5 min, and incubated with 600 µl of 10 mg/ml BSA and 0.1% SDS–PBS on a shaker overnight at room temperature. CSPFL-coupled beads were finally kept in a sonicating water bath for 30 min and adjusted to 7.5 × 10⁷ beads/ml in 10 mg/ml BSA, 5% glycerol, 0.1% sodium azide and 0.1% SDS–PBS. Beads were stored at 4°C until use.

Sterile 96-well U-bottom plates (Corning) were blocked with 1% BSA-PBS and incubated overnight at 37°C. Plates were washed three times with 300 µl/well of 0.05% Tween 20–PBS, and 20 µL/well of CSPFL-coupled beads were subsequently added. After three washes with 1% BSA-PBS, 50 µL/well of antibodies at 0.1 µg/mL in 1% BSA-PBS were added to the plate and incubated for 1 h at room temperature. Plates were then washed three times and 7.5×10⁴ THP-1 cells/well were added to the plate and incubated with opsonized beads for 40 min at 37°C. Phagocytosis was stopped by centrifugation at 500 g for 7 min at 4°C. Cells were then fixed with 150 µL/well of 2% paraformaldehyde for 30 minutes at 4°C. Fixed cells were then centrifuged at 500 g for 7 minutes at room temperature, and resuspended in 100 µL of PBS, before being analyzed in a CytoFLEX (Beckman Coulter) flow cytometer. Experiments were performed in duplicate, including positive (L9, CIS43, 311) and negative (mGO.53) controls in at least two independent experiments. The data were analyzed with FlowJo 10 (v10.8.1, FlowJo LLC, Ashland, OR). ADCP scores were calculated by dividing the fluorescence signals (% positive cells x geometric MFI positive cells) given by antibodies, by the one of the negative control antibody, mGO.53.

### Skin challenge

Antibodies (150 µg in 150µL PBS) were transferred intravenously into 4-week-old female C57BL/6JRj mice (Janvier Lab). The protective antibody 317 and the isotype IgG control mGO.53 were used as positive and negative controls. To quantity serum antibody levels 24 h after transfer, blood was collected in Microvette 200 Serum Gel tubes (Sarstedt) and centrifuged for 5 min at 10,000xg. Mice were challenged with 5,000 sporozoites into the footpad skin using a NanoFil syringue (35G, Word Precision Instruments) 24h after antibody administration. Parasitemia was followed 3-10 days post-infection by scoring GFP^+^ events on 250,000-500,000 gated erythrocytes by flow cytometry (CytoFLEX S, Beckman Coulter).

### Mosquito bite challenge

Antibodies (150 µg in 150 µL PBS) were transferred intravenously into 4-week-old female C57BL/6JRj mice (Janvier Lab). The protective antibody 317 and the isotype IgG control mGO53 were used as positive and negative controls, respectively. To quantity serum antibody levels 24 h after transfer, blood was collected in Microvette 200 Serum Gel tubes (Sarstedt), centrifuged for 5 min at 10,000xg. Mice were anesthetized with 6 mg/kg Xylazine Rompun 2% and 58 mg/mL ketamine 24 h following antibody administration. Nine infected (salivary glands GFP^+^) mosquitoes were allowed to feed on mice for 20 min at 21°C with animals rotated every 5 min. Following feeding, mosquito abdomens were inspected to confirm blood meal. Parasitemia was followed 3-10 days post-infection by scoring GFP^+^ events on 250,000-500,000 gated erythrocytes by flow cytometry (CytoFLEX S, Beckman Coulter).

### Cryo-EM Fab production, sample preparation and data collection

IgH and IgK genes were cloned into human Igγ1- and Igκ-expressing vectors, respectively, and expressed through the proprietary TurboCHO system (Genscript) or in Expi293 cells (in-house expression). For in-house purification, after 5 days of expression, secreted IgG were captured from the cell media onto Protein G resin and eluted with 0.1M glycine. Enriched IgG was further purified by size-exclusion chromatography (SEC) in TBS on an S200increase column (Cytiva). IgG antibodies were digested to Fab fragments with papain incubation for 4h at 37°C before being purified by size-exclusion chromatography. Purified Fab fragments and rsCSP were mixed overnight at a 10:1 molar ratio and then loaded on an S200 increase column (Cytiva) in TBS. rsCSP-antibody complex fractions were isolated, concentrated with a 30.000 MWCO concentrator to the smallest volume achievable, resulting in working concentrations of at least 1 mg/ml. Samples from SEC-purified complexes were loaded onto a Quantifoil 1.2/1.3 grid glow discharged at 25 mAmp for 15 sec. Samples were blotted using a Vitrobot Mark IV (ThermoFisher) at 4°C and 100% humidity and flash-frozen in liquid ethane.

Data for Fab-rsCSP complexes were collected on a Glacios II transmission electron microscope (ThermoFisher) equipped with a Falcon IV direct electron detector (ThermoFisher) at a pixel size of 0.725 Å^2^ (ISG5.143, ISG5.144, ISG5.163 and ISG6.149), or a Falcon 4(i) direct electron detector (ThermoFisher), at a pixel size of 0.718 Å^2^ (ISG6.149GL-reverted variants). Automated data collection (EPU, ThermoFisher) was set-up to collect 1500-2000 movies based on particle density. After alignment with Patch Motion Correction and CTF correction with Patch CTF in CryoSPARC Live (*111*), particles were picked with a blob picker. Binned particles were subject to an initial round of 2D classification to remove junk particles and classes displaying single, distinct complexes with high-resolution features were selected for further processing. The selected particles were brought to 3D Ab-initio and of the best 3D model was then refined with 3D refinement. Maps were then further processed with Global CTF refinement, NU Refinement and finally Local Refinement with Rotation Search extent of 2° and Shift search extent of 1 Å. No symmetry was applied at any stage to any of the complexes.

### Cryo-EM protein model building and refinement

An initial model of the Fab fragments was generated with Modelangelo (*112*) on the sharpened maps. Coot (*113*) was used for manual building, followed with iterations of real-space refined with Phenix refine (*114*) until no Ramachandran outliers remained subsisted and the clash scores, geometry scores and model fits criteria were optimized. Structural figures were made using USCF ChimeraX (*115*). Buried surface area was measured with the command *measure buriedarea [HC+LC] with [Epitope]*.

### Protein expression, purification, and preparation of Fab-peptide complexes for crystallization

Fab fragments of ISG5.163, ISG5.144, and ISG6.149 and their fully germline-reverted versions were transiently expressed in ExpiCHO cells and purified using a HiTrap Protein G HP column (GE Healthcare), followed by size-exclusion chromatography on a Superdex 200 16/90 column (GE Healthcare) equilibrated in Tris-buffered saline (TBS; 50 mM Tris, pH 7.8, 140 mM NaCl). The CSP-derived peptides were synthesized with N-terminal acetylation and C-terminal amidation. The peptide sequences of long and short repeat regions were Ac-NANPNANPNANPNANPNANPNANP-NH₂ (NANP_6_) and Ac-NPNANPNANPNA-NH₂ (NPNA_3_). Innopep Inc. synthesized peptides with a purity of over 98%. Purified Fabs were concentrated to 10 mg/mL in TBS using 10 kDa Amicon Ultra centrifugal filters and incubated with the peptide at a 1:5 Fab-to-peptide molar ratio overnight at 4°C before crystallization.

### X-ray crystallization of Fab-peptide complexes

Crystallization screening was performed using a high-throughput CrystalMation robotic system (Rigaku) at The Scripps Research Institute (TSRI). Crystals were grown using the sitting-drop vapor diffusion method, with 35 μL reservoir solutions and 0.2 μL drops consisting of 0.1 μL protein solution mixed with 0.1 μL precipitant solution. Fab ISG5.163 in complex with NANP₆ crystallized in a solution containing 0.1 M sodium citrate, 20% (v/v) 2-propanol, 10% (v/v) ethylene glycol, and 20% (w/v) PEG 4000. Fab ISG5.144 in complex with the NANP₆ peptide crystallized in 0.17 M ammonium sulfate, 15% (v/v) glycerol, and 25.5% (w/v) PEG 4000. Fab ISG6.149 in complex with the NPNA₃ peptide crystallized in 0.1 M Tris (pH 7.0) and 50% (v/v) PEG 200. The corresponding germline-reverted variants were crystallized under similar conditions. Fab ISG5.163_GL in complex with the NANP₆ peptide crystallized in 0.2 M ammonium sulfate and 20% (w/v) PEG 3350. Fab ISG5.144_GL in complex with a NANP_6_ peptide crystallized in 0.2 M sodium chloride, 1.0 M sodium citrate, 0.1 M Tris (pH 7.0), and 10% (v/v) ethylene glycol. Fab ISG6.149_GL in complex with the NPNA₃ peptide crystallized in 0.2 M potassium citrate, 10% (v/v) ethylene glycol, and 20% (w/v) PEG 3350. All crystals were grown at 293 K. Crystals obtained from conditions lacking a cryoprotectant were cryoprotected by soaking in the corresponding mother liquor supplemented with 10% (v/v) ethylene glycol before data collection.

### X-ray data collection and structure determination

X-ray diffraction data for matured and germline-reverted Fabs were collected at beamline BL12-1 of the Stanford Synchrotron Radiation Lightsource (SSRL). Diffraction data were processed and scaled using HKL-2000 (*116*). The structures were solved by molecular replacement using PHASER (*117*), with AlphaFold2-generated homology models (*118*) as search templates. Initial model building was performed in Coot (*113*) followed by iterative refinement using phenix.refine (*114*). Peptide molecules were manually positioned into Fo–Fc electron density maps and subsequently refined through iterative cycles in phenix.refine. Fab residues were numbered according to the Kabat nomenclature.

### Statistical analyses

Comparison of the different immunoglobulin gene features between groups were assayed using two-sided 2×2 or 2×5 Fisher’s Exact tests with SISA online tools (http://www.quantitativeskills.com/sisa). The number of somatic mutations, and antiparasitic properties were compared across groups of antibodies using two-tailed Mann-Whitney test. Volcano plot comparing Ig gene features (n=239 parameters) of *Pf*CSP B cells was performed using GraphPad Prism software (v10.1.0 GraphPad Prism Inc.). The y axis indicates the statistics expressed as -log_10_ (p-values) and the x axis represents the differences between the group means for each parameter. Spearman rank correlation coefficients were calculated with two-tailed p-values with 95% confidence intervals. For the survival curves, comparisons were tested using the Holm-Šídák multiple comparisons test. Test Statistics are described in figure captions and all data were plotted using GraphPad Prism (v10.1.0 GraphPad Prism Inc.).

## ACKNOWLEDGEMENTS

We are grateful to the volunteers and their families, and the clinical, field, and laboratory teams at the research institutions. We thank the MAL067 Vaccine Immunology Consortium investigators and working groups and GSK for their support in 2008–09, for developing the study plan and for transferring custodianship of the MAL067 samples to ISGlobal. GSK was provided the opportunity to review a preliminary version of this manuscript for factual accuracy, but the authors are solely responsible for final content and interpretation. We are grateful to the Malaria Immunology Group (ISGlobal), and especially Laura Puyol and Diana Barrios for technical assistance. We also thank the Cytometry Platform of the Institut de Investigacions Biomèdiques August Pi I Sunyer (IDIBAPS) for technical support. We thank the team of the Center for Production and Infection of Anopheles (CEPIA, C2RA, Institut Pasteur) for providing and infecting mosquitoes, the staff of Central Animal Facility (C2RA, Institut Pasteur) for animal care, and the team of the Photonic BioImaging platform (UTechS PBI, C2RT, Institut Pasteur) for providing access to the spinning-disk microscope and support. We acknowledge the Malaria Infection and Immunity team (Institut Pasteur) and specially Susanna Celli and Pauline Formaglio for technical assistance for the antibody passive transfer. Expression Ig vectors were kindly provided by Hugo Mouquet (Institut Pasteur). The monoclonal antibody 3D11 was obtained through BEI Resources, NIAID, NIH: Hybridoma 3D11 Anti-Plasmodium berghei 44-kDa Sporozoite Surface Protein, MRA-100, contributed by Victor Nussenzweig. Use of the Stanford Synchrotron Radiation Lightsource, SLAC National Accelerator Laboratory, is supported by the U.S. Department of Energy, Office of Science, Office of Basic Energy Sciences under Contract No. DE-AC02-76SF00515.

## Author contributions

E.B., F.D., F.C., M.J., K.B., R.A., and L.M.M-A. designed, performed and analyzed experiments. W.L. performed cryo-EM data collection and processing. C.S. and D.M. designed and performed bioinformatic analyses. C.C., I.A.W., A.B.W., R.A., G.M., C.D., and L.M.M-A. supervised experiments. P.A. organized volunteer recruitment, clinical follow up, and provided clinical data. C.J., J.J.C., G.M., and C.D. organized sample collection and processing. J.J.C., G.M., and C.D. conceived and developed the MAL067 study framework. L.M.M-A. conceived and coordinated the study, and wrote the manuscript. E.B., F.D., F.C., R.A., M.J., I.A.W., A.B.W., G.M., C.D., and L.M.M-A. wrote, reviewed and edited the manuscript. All authors approved the final version of the manuscript.

## Data and Materials Availability

the authors declare that all data needed to evaluate the conclusions in this study are present in the manuscript and/or the supplementary materials. The cryo-EM density maps and models of ISG5.143, ISG5.144, ISG5.163, ISG6.149, ISG6.149GL_LC and ISG6.149GL_HC have been deposited in the Protein Data Bank (PDB) and Electron Microscopy Data Bank (EMDB) under accession codes 38XB (EMD-79206), 38XC (EMD-79207), 38XA (EMD-79205), 38XD (EMD-79208), 38XE (EMD-79209) and 38XF (EMD-79210), respectively. The crystal structure factors and atomic coordinates of ISG6.149, ISG5.144, ISG5.163, ISG6.149GL, ISG5.144GL, and ISG5.163GL have been deposited into the PDB under accession codes 38VS, 38VP, 38VO, 38VR, 38VQ, and 38VN, respectively.

