## Supplementary Materials for "Protective near-germline antibodies to malaria elicited by RTS,S/AS01_E_ vaccination in African infants"

**The PDF file includes:**
Figs. S1 to S18
Tables S1 to S8

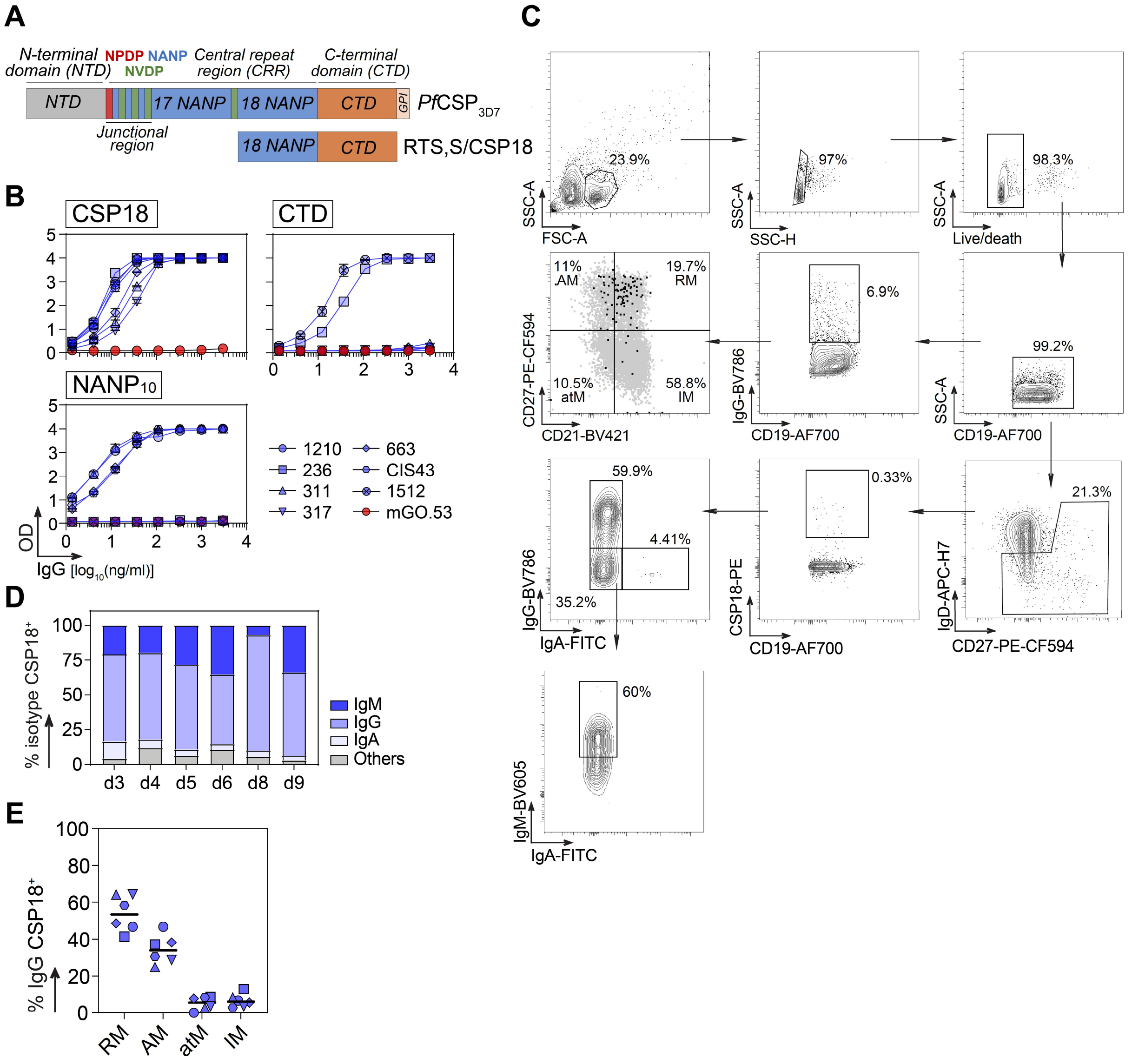


**Figure S1**. **CSP18 antigenicity and memory B cells from infants immunized with RTS,S/AS01$_E$**. (**A**) Sequence diagram of the *Pf*CSP protein (top) and the RTS,S/CSP18 protein construct (bottom). (**B**) ELISA titration graphs showing the reactivity of benchmarked *Pf*CSP antibodies (blue symbols), and a negative control (mGO.53, red circles) to CSP18, NANP$_{10}$ and CTD proteins. (**C**) Representative flow cytograms showing the gating strategy used to identify circulating blood CSP18-reactive IgG$^+$ B cells prior to single cell sorting, and memory B-cell (MBC) analysis. (**D**) Bar graph showing the isotype distribution of CSP18-reactive MBC for each donor (d). (**E**) Graph showing the percentage of CSP18-reactive IgG$^+$ MBC according with their phenotype as defined in (C). RM, resting memory, AM, activated memory, atM, atypical memory, IM, intermediate memory.

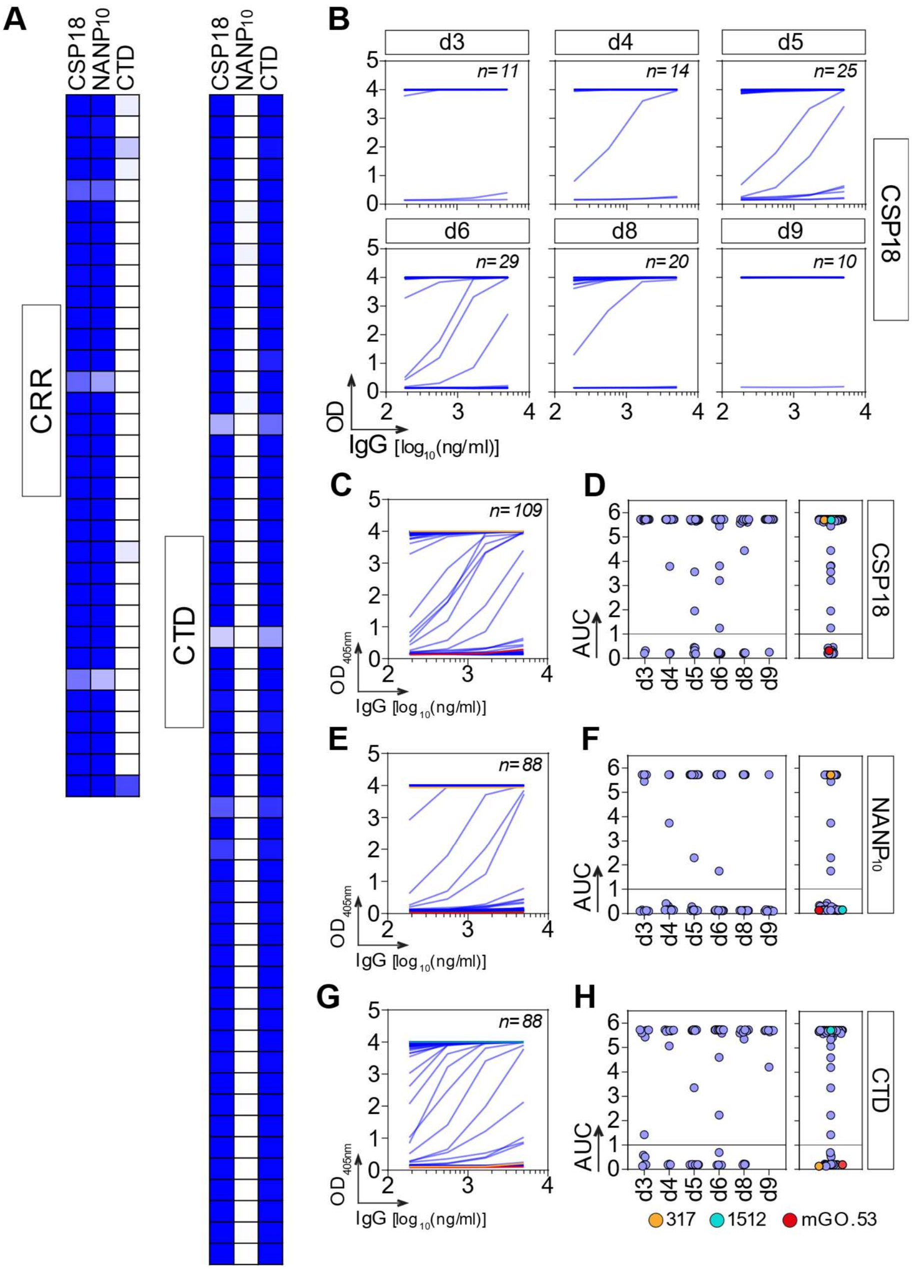


**Figure S2**. **Reactivity of CSP18 IgG B-cell antibodies**. (**A**) Heatmap showing the reactivity of CSP18-reactive antibodies (*n*=88) against CSP18, $NANP_{10}$ and CTD proteins, with color-coded cells according to the values shown in Table S2. The heatmap is segregated according to antibody specificity to the central repeat region (CRR, left) or the C-terminal domain (CTD, right). (**B**) Representative ELISA titration graphs showing the reactivity of antibodies cloned from $CSP18^+$ B cells. The number of tested antibodies is indicated in italics. (**C**) Same as in (B) but for all cloned antibodies together. (**D**) Dot plots showing the area under the curve calculated from graphs in (C), segregated by donors (left) or together (right). (**E-F**) Same as in (C-D), respectively, but for reactivity against the $NANP_{10}$ protein. (**G-H**) Same as in (E-F), respectively, but for reactivity against the CTD protein. The number of antibodies are indicated in italics. Positive controls for $NANP_{10}$ (317, orange) and CTD (1512, cyan), and negative isotype control (mGO.53, red) are indicated.

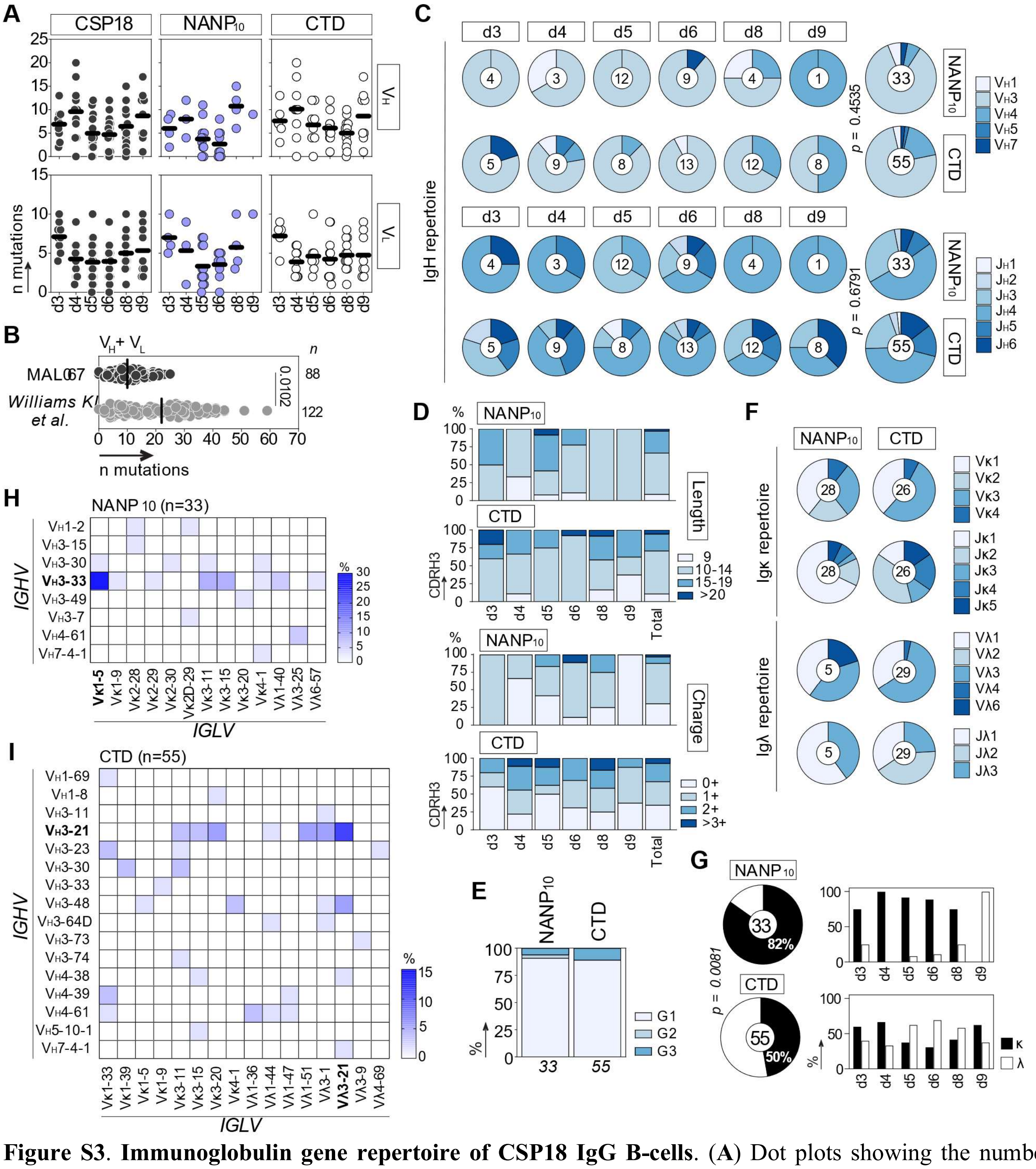


**Figure S3**. **Immunoglobulin gene repertoire of CSP18 IgG B-cells**. (**A**) Dot plots showing the number of nucleotide mutations in $V_H$ (top) and $V_L$ (bottom) genes of all the CSP18 B-cell antibodies (n=88, black), or grouped by $NANP_{10}$ (n=33, blue) and C-terminal domain (CTD, n=54, white) reactivity. Averaged mutation per donor is indicated by the solid black lines. (**B**) Dot plot comparing the number of somatic mutations in $V_H$+$V_L$ genes of all CSP18 antibodies from our collection, and *Pf*CSP antibodies cloned from RTS,S-vaccinated adults (data from *Williams KL et al, (2024), Nat. Med. 30:117-129, Doi: 10.1038/s41591-023-02659-z*). Groups were compared with the two- tailed Mann- Whitney test. *p* value is indicated. (**C**) Pie charts comparing the distribution of $V_H$ and $J_H$ gene usage of $NANP_{10}$ and CTD B-cell antibodies. The number of antibody sequences analyzed is indicated in the center of each pie chart. (**D**) Bar graph showing the distribution of amino acid length (top) and total positive charges (bottom) of the CDRH3 of $NANP_{10}$ and CTD B-cell antibodies. (**E**) Bar graph showing the distribution of IgG subclass of $NANP_{10}$ and CTD B-cell antibodies. (**F**) Same as in (C) but for the Igk and Igλ repertoire. (**G**) Pie chart (left) and bar graphs (right) comparing the distribution of kappa vs lambda Ig chain usage between $NANP_{10}$ and CTD B-cell antibodies. (**H**) Heatmap showing the frequency of $NANP_{10}$ B-cell antibodies encoded by the indicated IGHV and IGHL pairs. (**I**) Same as in (H) but for CTD B-cell antibodies. Groups were compared using 2x5 Fisher's Exact test in (C) and (G).

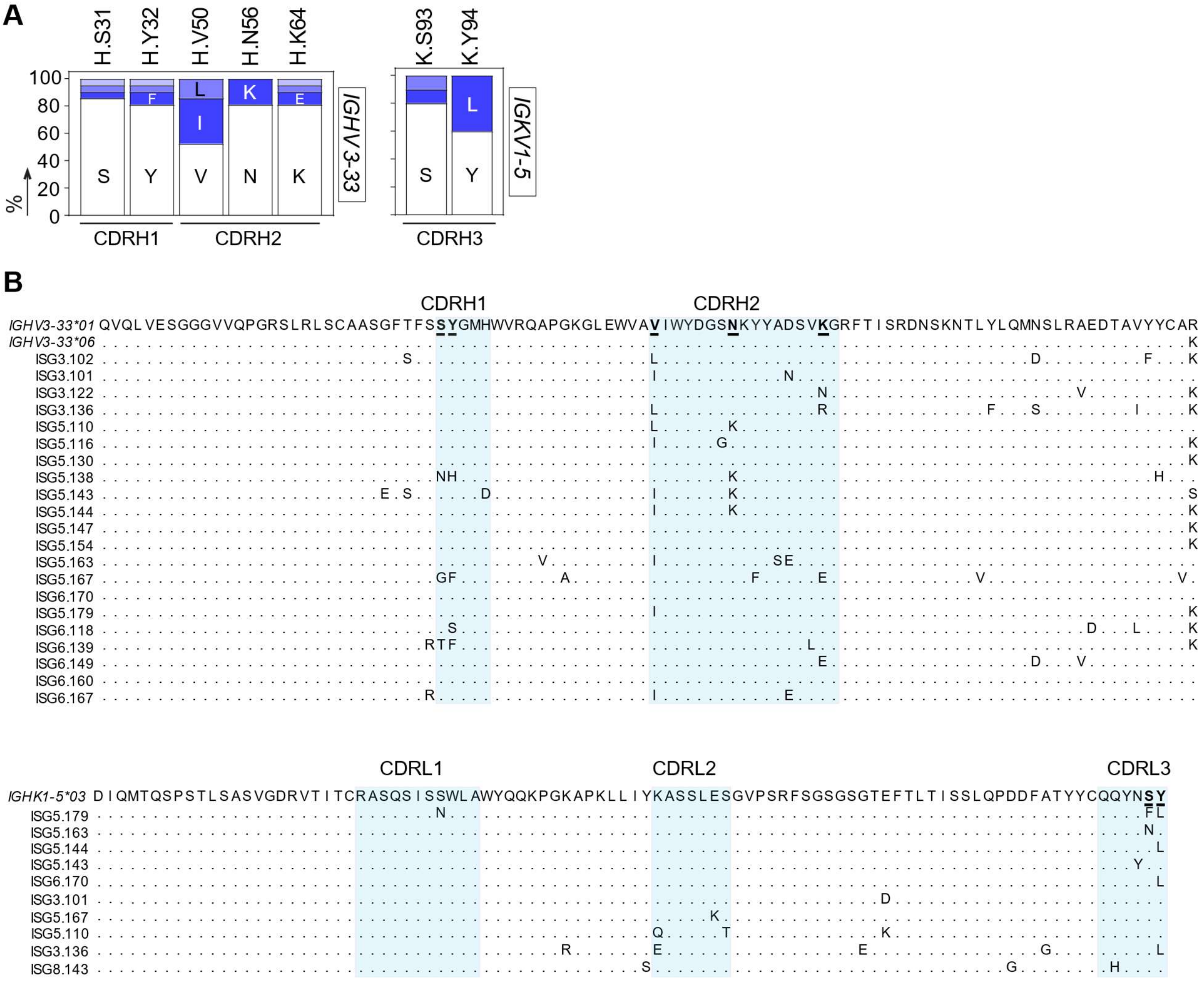


**Figure S4**. **Amino acid replacements and alignment of *IGHV3-33/IGK1-5* NANP$_{10}$ antibodies.** (**A**) Bar graphs showing the proportion of amino acid replacements at the indicated positions of *IGHV3-33* (left) and *IGK1-5* (right) gene segments of NANP$_{10}$ antibodies. Amino acids of unmutated genes are indicated in white and replacements are indicated in shades of blue. (**B**) Amino acid sequence alignment of *IGHV3-33* (top) and *IGK1-5* (bottom) gene segment alleles of NANP$_{10}$ antibodies.

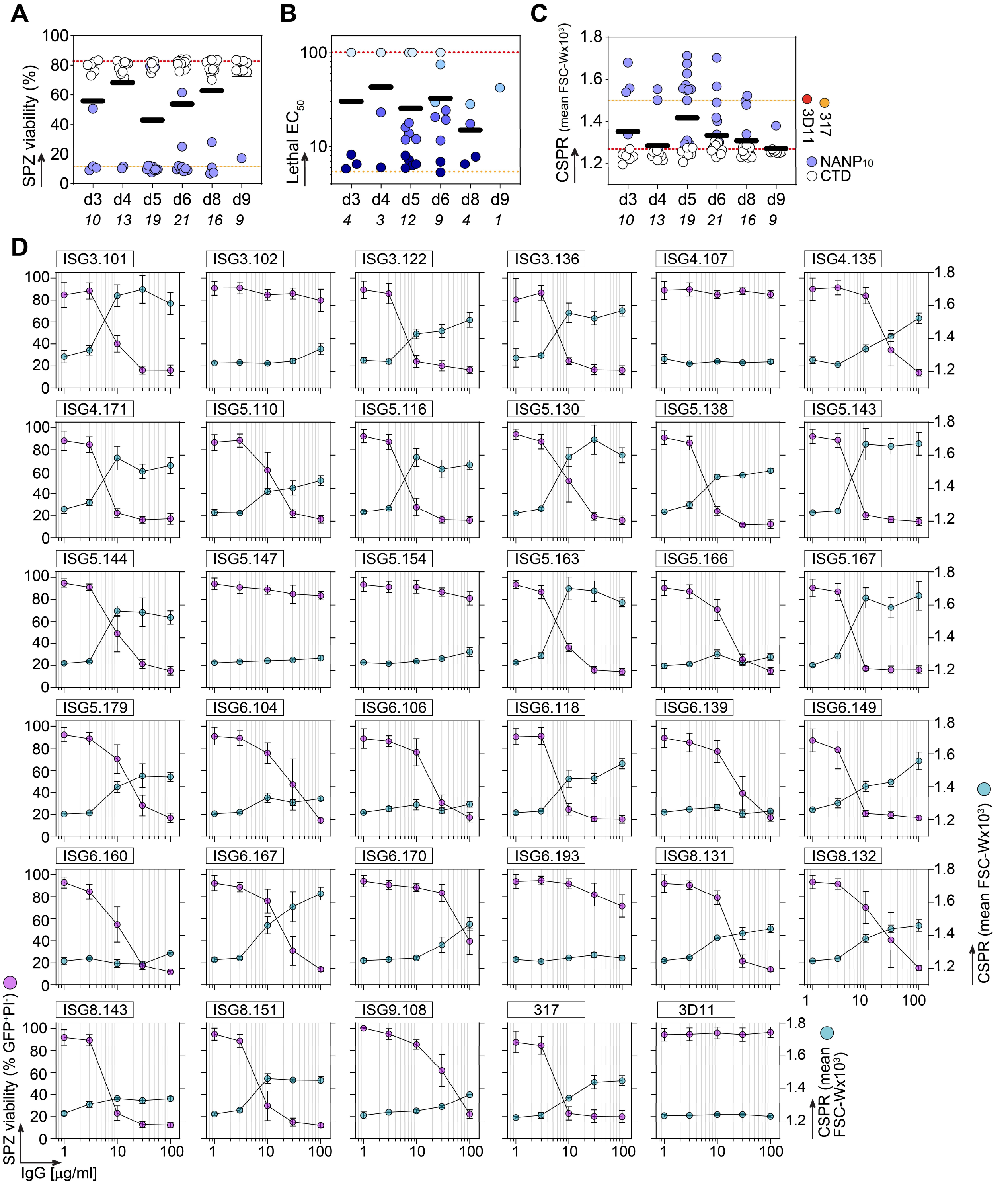


**Figure S5. *In vitro* antiparasitic properties of CSP18 antibodies**. (**A**) Dot plot comparing the Fab-dependent lethality induced in sporozoites (SPZ) by $NANP_{10}$ and C-terminal domain (CTD) antibodies at 100 µg/ml, after 45 min of incubation at 37ºC, segregated by donor (d), tested in duplicate in two independent experiments. (**B**)

Dot plot showing the lethal half-maximal effective concentration ($EC_{50}$) values induced in SPZ by $NANP_{10}$ antibodies (n=3 independent experiments, see [D]). $EC_{50}$ values color coded in shades of blue, with darker shades indicating higher potency. (**C**) Same as in (A), but for the circumsporozoite precipitation reaction (CSPR). (**D**) Graph depicting the distinct *in vitro* lethality (purple) and CSPR (cyan) induced in SPZ by $NANP_{10}$ antibodies after 45 min of incubation at 37ºC (n = 3 independent experiments, data are presented as mean ± SEM).

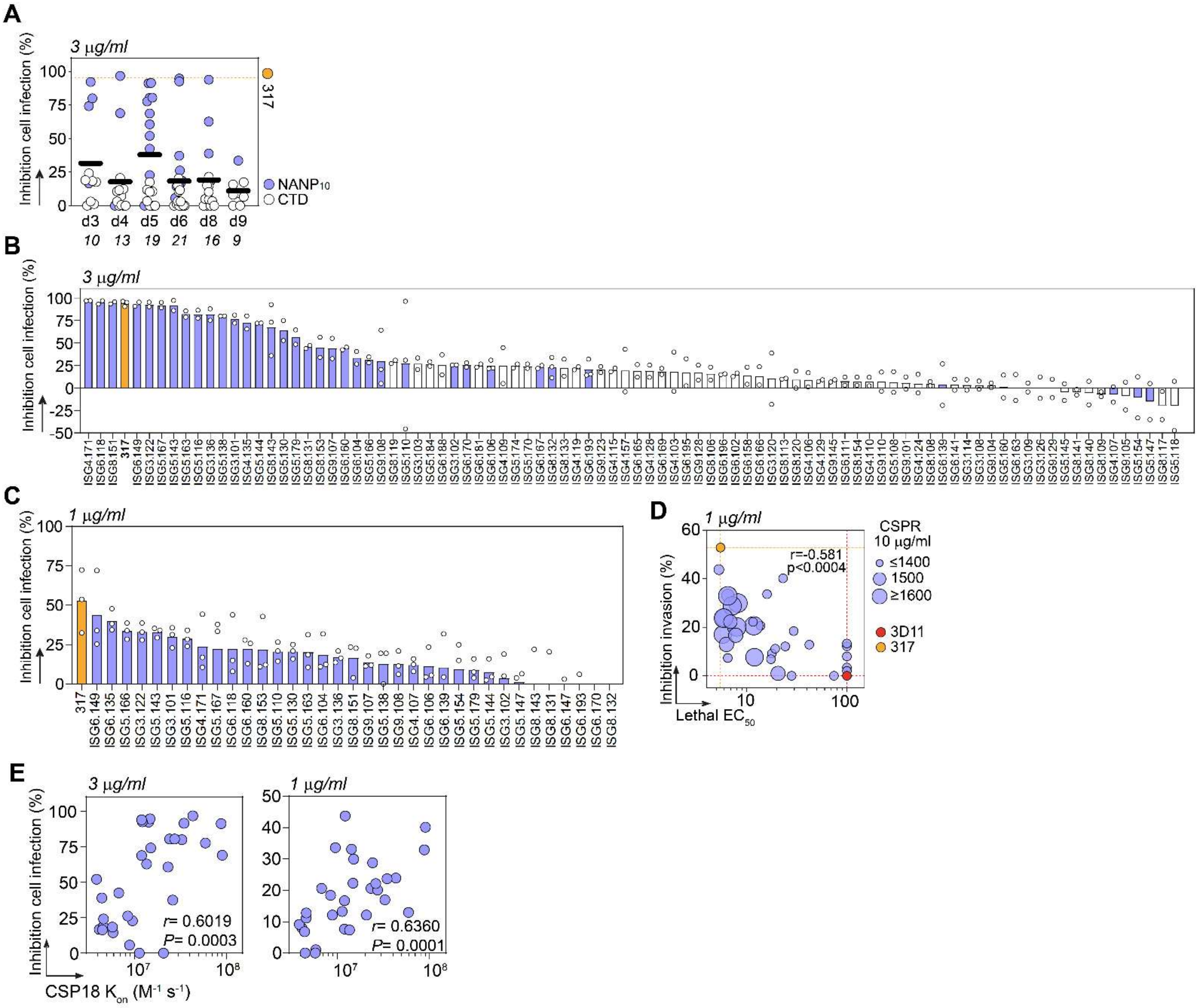


**Figure S6.** ***In vitro*** **inhibition of hepatocyte invasion by CSP18 antibodies**. (**A**) Dot plot comparing the percentage of sporozoite (SPZ) inhibition to invade HepG2 cells induced by $NANP_{10}$ (blue) and CTD (white) antibodies, after two hours of incubation at 37ºC, tested at 3 µg/ml in 2-3 independent experiments, segregated by donor. (**B**) Bar graphs showing the percentage of SPZ inhibition to invade HepG2 cells induced by $NANP_{10}$ (blue) and CTD (white) antibodies for each antibody tested at 3 µg/ml in 2-3 independent experiments. Each dot represents the value of an independent experiment. (**C**) Same as in (B), but for $NANP_{10}$ antibodies after 48h of incubation at 37 ºC, tested at 1 µg/ml in three independent experiments. (**D**) Bubble plot showing the lethal half-maximal effective concentration ($EC_{50}$) values versus the percentage of inhibition of HepG2 invasion induced by $NANP_{10}$ antibodies after 48 h of incubation at 37ºC, tested at 1 µg/ml. The size of the bubbles is proportional to the elongated CSPR phenotype induced by the antibodies at 10 µg/ml. (**E**) Correlation plots showing the association constant ($K_{on}$) to CSP18 versus the percentage of inhibition of HepG2 invasion of $NANP_{10}$ antibodies tested at 3 (left) and 1 (right) µg/ml. The values for the *Pf* inhibitory 317 (orange), and the *Pb* 3D11 (red) negative control antibodies are indicated in colored dashed lines or dots. The two-sided Spearman rho correlation coefficients and corresponding *p* values are shown in (D) and (E).

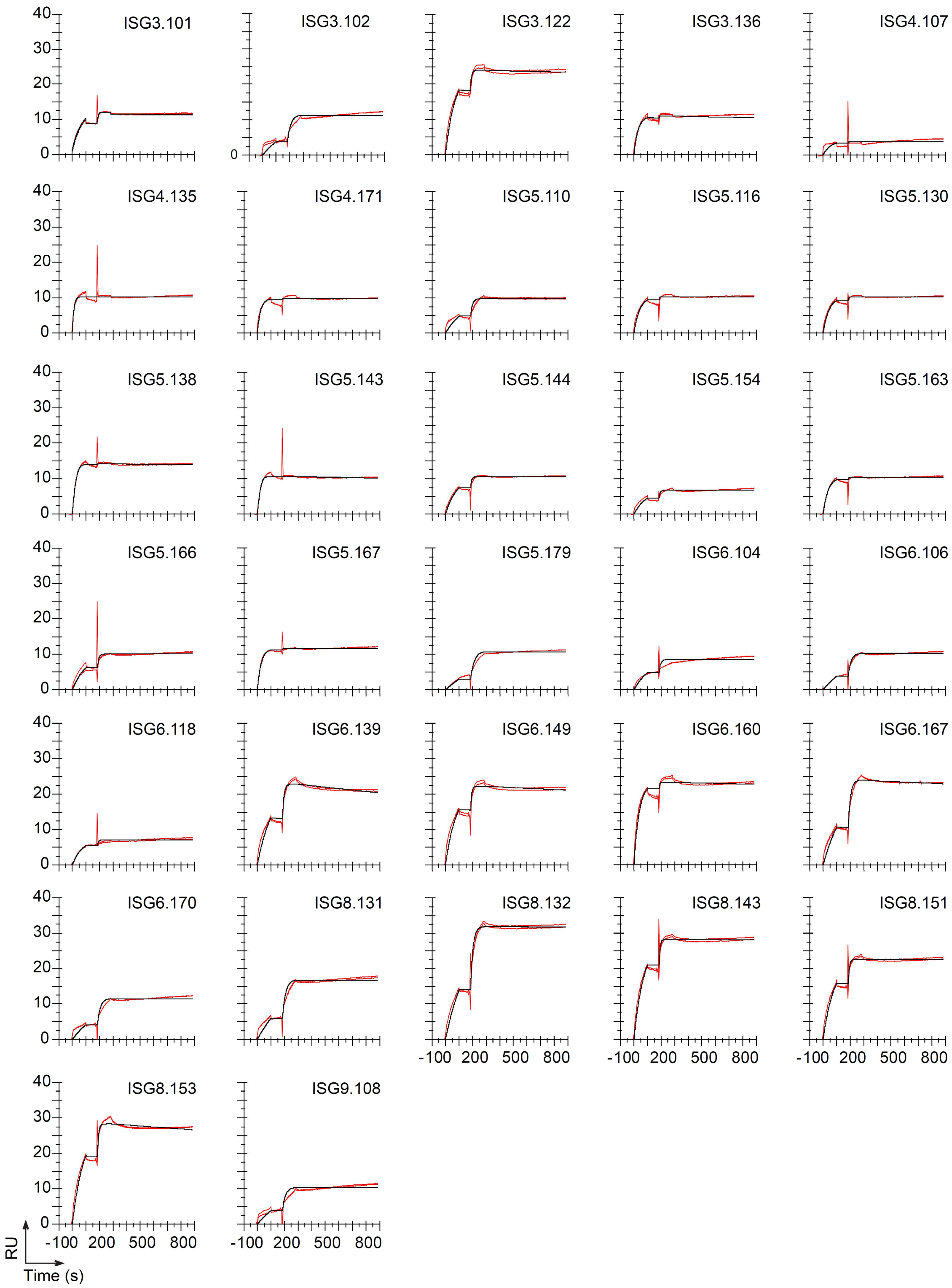


**Figure S7. Surface plasmon resonance (SPR) of NANP$_{10}$ antibodies.** Sensorgrams showing the single-cycle kinetics (SCK) of NANP$_{10}$ antibodies binding to CSP18.

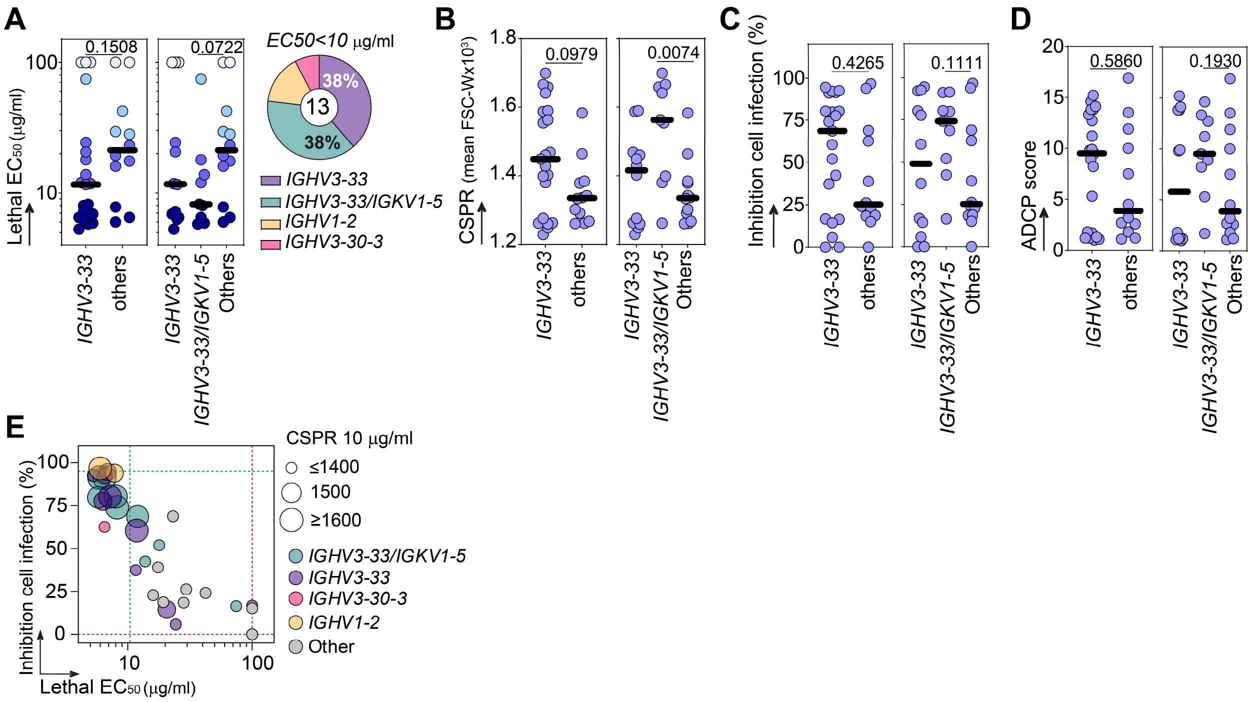


**Figure S8**. ***In vitro* antiparasitic properties of *IGHV3-33* $NANP_{10}$ antibodies**. (**A**) Dot plots comparing the lethal half-maximal effective concentration ($EC_{50}$) values induced in sporozoites(SPZ) by $NANP_{10}$ antibodies (n=3 independent experiments) between those encoded by *IGHV3-33* (left), matched *IGHV3-33/IGKV1-5* (right), and other gene segments. The dots are coded in shades of blue, with darker shades indicating higher lethal $EC_{50}$ potency, as indicated in Fig 2B. The pie chart shows the distribution of $NANP_{10}$ antibodies with $EC_{50}$ values below 10 µg/ml, according with the color coded *IGHV* gene family used. (**B**) Dot plots as in (A) but for the CSPR measured at 10 µg/ml. (**C**) Dot plots as in (A) but for the percentages of inhibition of cell infection, measured at 3 µg/ml. (**D**) Dot plots as in (A) but for the ADCP score. (**E**) Bubble plot showing the lethal $EC_{50}$ values versus the percentage of inhibition of cell infection induced by $NANP_{10}$ antibodies, with color coded *IGHV/IGHL* gene families. The size of the bubbles is proportional to the elongated CSPR phenotype induced by the antibodies at 10 µg/ml.

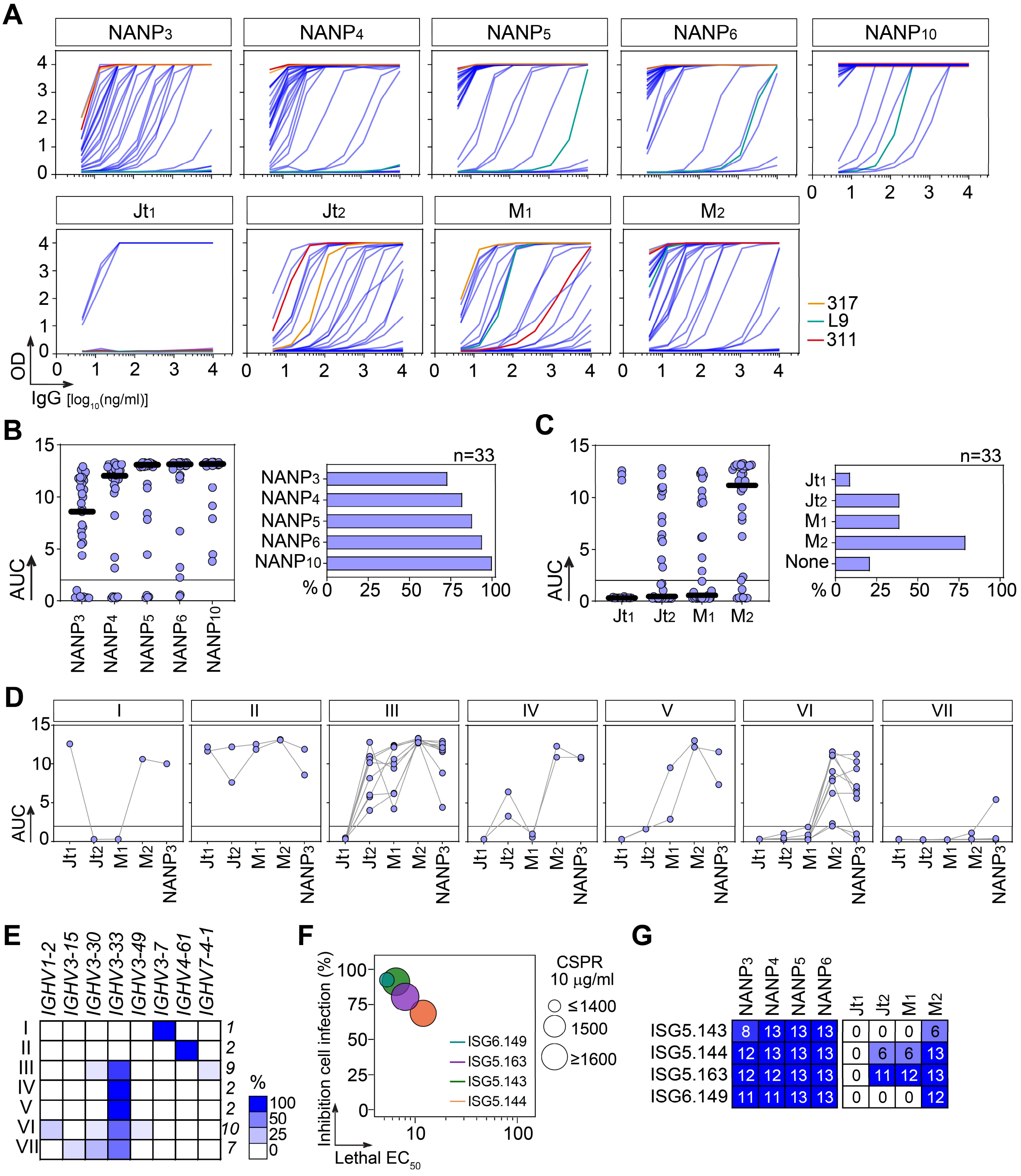


**Figure S9**. **Binding profiles of central repeat region (CRR) antibodies**. (**A**) Representative ELISA titration graphs of three independent experiments tested in duplicate, showing the reactivity of NANP$_{10}$ antibodies against peptides with incremental NANP repetitions (NANP$_{3-10}$, top) and peptides Jt$_1$, Jt$_2$, M$_1$ and M$_2$ (bottom). Central (311 and 317) and minor (L9) repeat control antibodies are shown. (**B**) Dot plot (left) showing the area under the curve (AUC) calculated from the ELISA graphs in (B) against NANP$_{3-10}$. The bar graph shows the proportion of antibodies recognizing each peptide. (**C**) Same as in (B) but for peptides Jt$_1$, Jt$_2$, M$_1$ and M$_2$. (**D**) Graphs showing the ELISA AUC of antibodies against the indicated peptides, grouped by different binding profiles (I-VII). (**E**)

Heatmap showing the proportion of *IGHV* gene segments used by the antibodies grouped by binding profiles I-VII from (E). (**F**) Bubble plot showing the lethal half-maximal effective concentration ($EC_{50}$) values *versus* the percentage of inhibition of cell infection induced by antibodies ISG5.143, ISG5.144, ISG5.163 and ISG6.149. The size of the bubbles is proportional to the elongated CSPR phenotype induced by the antibodies at 10 µg/ml. (**G**) Heatmap showing the ELISA AUC values of ISG5.143, ISG5.144, ISG5.163 and ISG6.149 against $NANP_{3-6}$ and $Jt_1$, $Jt_2$, $M_1$ and $M_2$ peptides. The color-coded cells are expressed according to the ELISA area under the curve (AUC) values with darker colors representing higher reactivity.

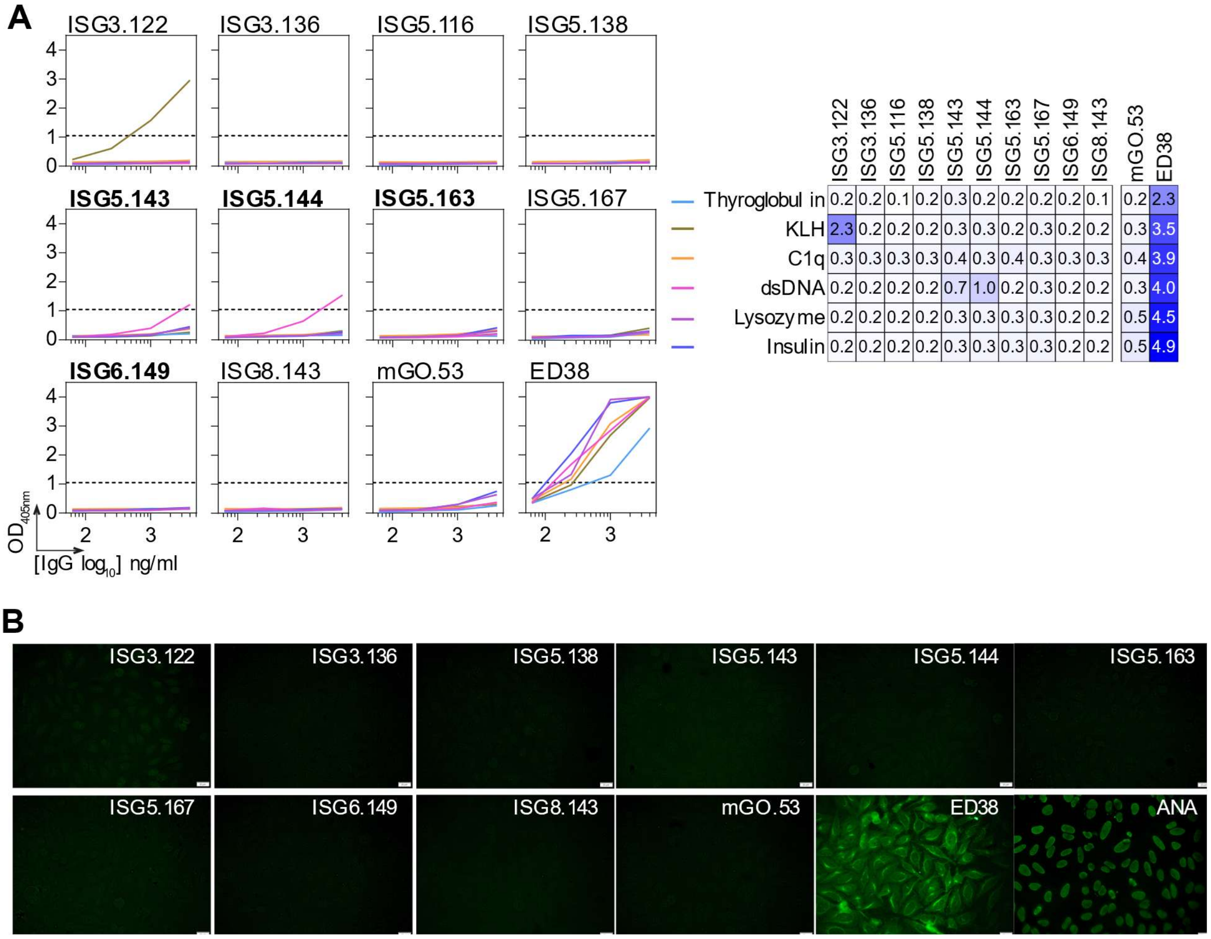


**Figure S10**. **Off-target binding characteristics of central repeat region (CRR) antibodies**. (**A**) Heatmap (right) showing the ELISA reactivity of selected CRR antibodies against a panel of structurally unrelated antigens. Polyreactive (ED38) and non-polyreactive (mGO.53) controls are included. Color values are proportional to the reactivity level measured as the area under the curve (AUC) of the ELISA graphs (left). (**B**) Microscopic images showing the reactivity of the antibodies in (A) to Hep2-expressing self-antigens assayed by indirect immunofluorescence assay. The negative (mGO.53) and positive (ED38, and kit's ANA) controls are included. The scale bars represent 40 μm.

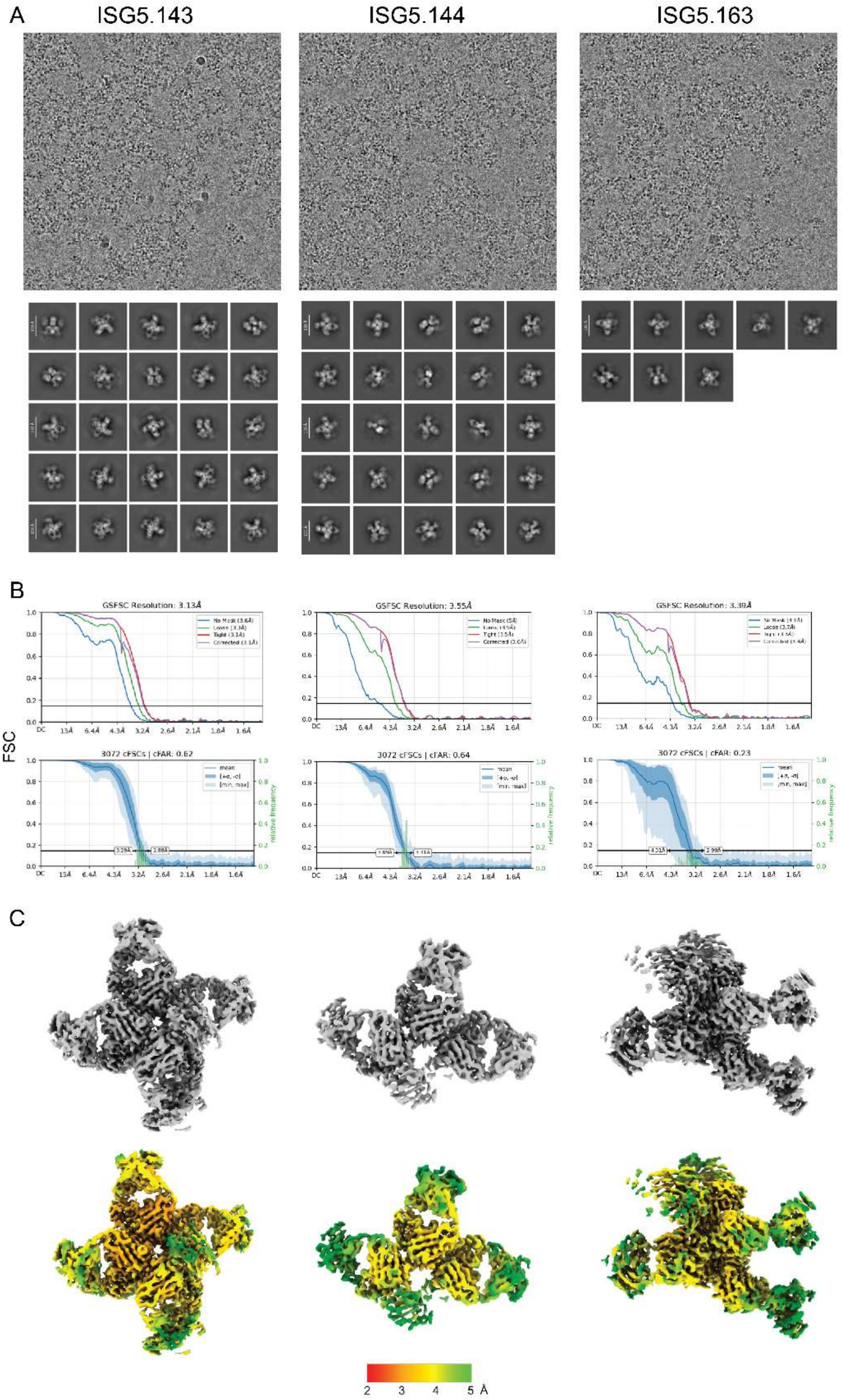


**Figure S11**. **CryoEM data processing of rsCSP protein in complex with ISG5.143, ISG5.144 and ISG5.163 Fabs**. (**A**) Representative electron micrograph (top), with a 3 Å low-pass filter for better contrast, and 2D classes (bottom). (**B**) Gold-standard Fourier-shell correlation (FSC) and cFAR FSC. (**C**) Overall map 3D volume, uncolored (top) and colored by local resolution (bottom).

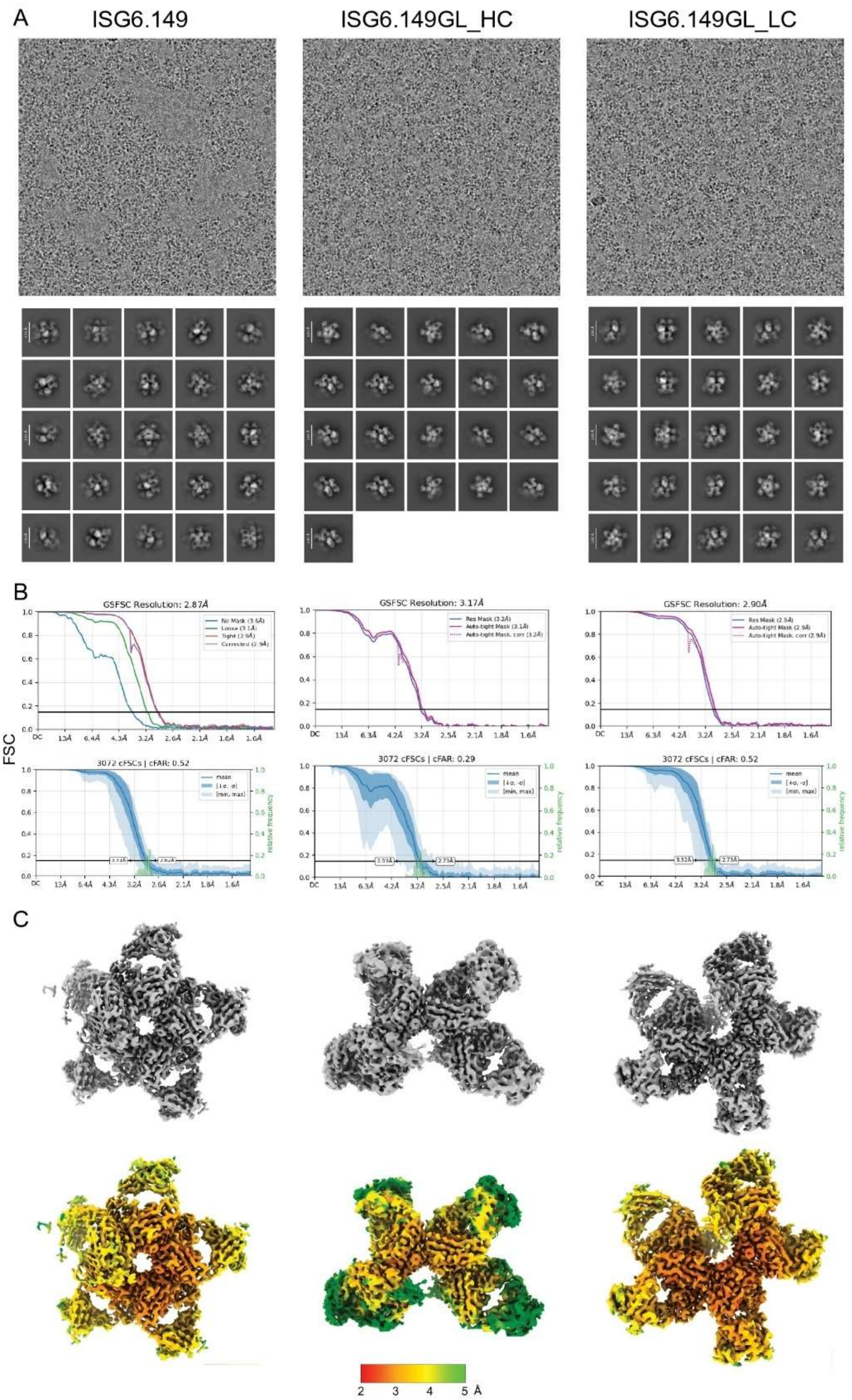


**Figure S12**. **CryoEM data processing of rsCSP protein in complex with ISG6.149, ISG6.149GL_HC and ISG6.149GL_LC Fabs**. (**A**) Representative electron micrograph (top), with a 3 Å low-pass filter for better contrast, and 2D classes (bottom). (**B**) Gold-standard fourier-shell correlation (FSC) and cFAR FSC. (**C**) Overall map 3D volume, uncolored (top) and colored by local resolution (bottom).

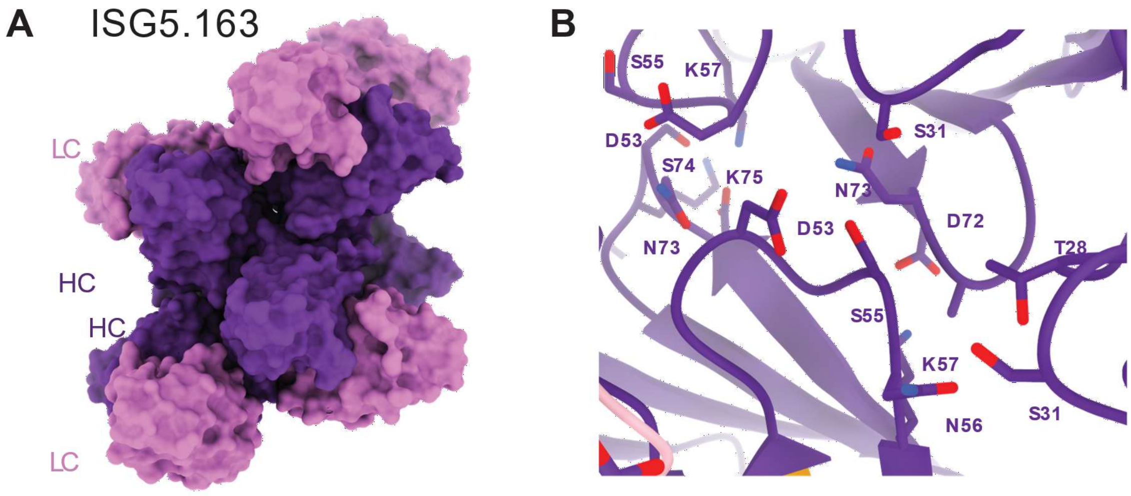


**Figure S13. Stacking of ISG5.163 homotypic spirals**. (**A**) Surface molecular representation of two spirals of ISG5.163 Fabs, mirroring each other and interacting by the heavy chain (HC). (**B**) Structural details of the interface between HC Fabs. Contacting residues are shown as sticks with heteroatoms colored (blue: N, red: O). Color key: purple (HC) and pink (light chain, LC).

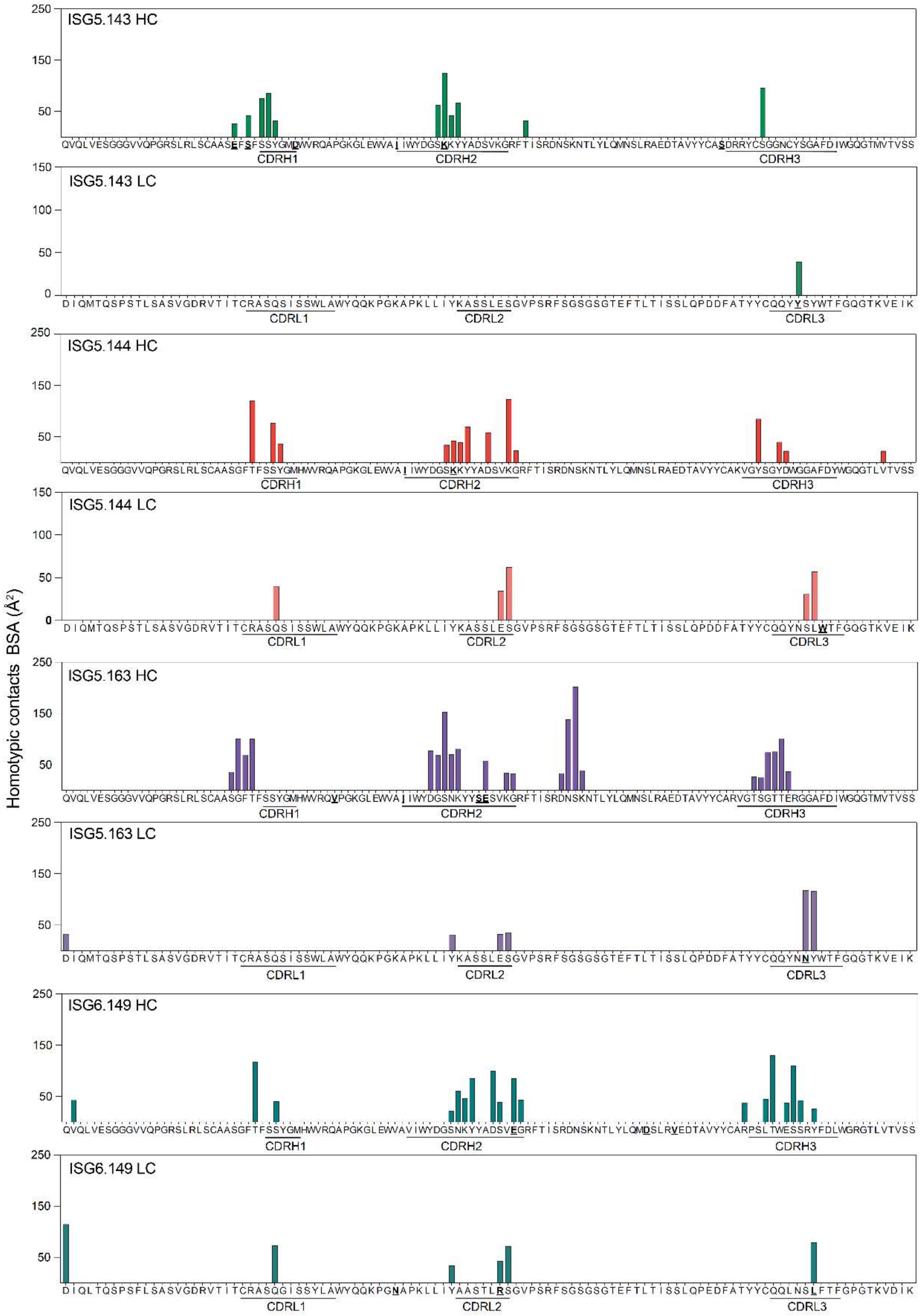


**Figure S14**. **Homotypic interfaces of ISG5.143, ISG5.144, ISG5.163 and ISG6.149**. Bar graphs showing per residue BSA contributions to Fab-Fab homotypic interfaces with rsCSP. Somatically mutated residues are highlighted in bold and underlined. HC, heavy chain. LC, light chain.

```
ISG5.143
                                              CDRH1                            CDRH2
ISG5.143GL_HC   QVQLVESGGGVVQPGRSLRLSCAASGFTFSSYGMHWVRQAPGKGLEWVAVIWYDGSNKYYADS
ISG5.143H       .........................E.S......D..............I......K......
                CDRH2                              CDRH3
ISG5.143GL_HC   VKGRFTISRDNSKNTLYLQMNSLRAEDTAVYYCARDRRYCSGGNCYSGAFDIWGQGTMVTVSS
ISG5.143H       ..................................S............................
                                        CDRK1                          CDRK2
ISG5.143GL_LC   DIQMTQSPSTLSASVGDRVTITCRASQSISSWLAWYQQKPGKAPKLLIYKASSLESGVPSRF
ISG5.143L       ..............................................................
                                           CDRK3
ISG5.143GL_LC   SGSGSGTEFTLTISSLQPDDFATYYCQQYNSYWTFGQGTKVEIK
ISG5.143L       .............................Y..............

ISG5.144
                                              CDRH1                            CDRH2
ISG5.144GL_HC   QVQLVESGGGVVQPGRSLRLSCAASGFTFSSYGMHWVRQAPGKGLEWVAVIWYDGSNKYYAD
ISG5.144H       .................................................I......K.....
                CDRH2                                    CDRH3
ISG5.144GL_HC   SVKGRFTISRDNSKNTLYLQMNSLRAEDTAVYYCAKVGYSGYDWGGAFDYWGQGTLVTVSS
ISG5.144H       .............................................................
                                        CDRK1                          CDRK2
ISG5.144GL_LC   DIQMTQSPSTLSASVGDRVTITCRASQSISSWLAWYQQKPGKAPKLLIYKASSLESGVPSRF
ISG5.144L       ..............................................................
                                           CDRK3
ISG5.144GL_LC   SGSGSGTEFTLTISSLQPDDFATYYCQQYNSYWTFGQGTKVEIK
ISG5.144L       ...............................L............

ISG5.163
                                              CDRH1                            CDRH2
ISG5.163GL_HC   QVQLVESGGGVVQPGRSLRLSCAASGFTFSSYGMHWVRQAPGKGLEWVAVIWYDGSNKYYADS
ISG5.163H       .......................................V.........I..........SE.
                CDRH2                              CDRH3
ISG5.163GL_HC   VKGRFTISRDNSKNTLYLQMNSLRAEDTAVYYCARVGTSGTTERGGAFDIWGQGTMVTVSS
ISG5.163H       .............................................................
                                        CDRK1                          CDRK2
ISG5.163GL_LC   DIQMTQSPSTLSASVGDRVTITCRASQSISSWLAWYQQKPGKAPKLLIYKASSLESGVPS
ISG5.163L       ............................................................
                                             CDRK3
ISG5.163GL_LC   RFSGSGSGTEFTLTISSLQPDDFATYYCQQYNSYWTFGQGTKVEIK
ISG5.163L       ................................N.............

ISG6.149
                                              CDRH1                            CDRH2
ISG6.149GL_HC   QVQLVESGGGVVQPGRSLRLSCAASGFTFSSYGMHWVRQAPGKGLEWVAVIWYDGSNKYYAD
ISG6.149H       ..............................................................
                CDRH2                                CDRH3
ISG6.149GL_HC   SVKGRFTISRDNSKNTLYLQMNSLRAEDTAVYYCARPSLTWESSRYFDLWGRGTLVTVSS
ISG6.149H       ..E..................D...V..................................
                                        CDRK1                          CDRK2
ISG6.149GL_LC   DIQLTQSPSFLSASVGDRVTITCRASQGISSYLAWYQQKPGKAPKLLIYAASTLQSGVPSRF
ISG6.149L       .........................................N............R.......
                                           CDRK3
ISG6.149GL_LC   SGSGSGTEFTLTISSLQPEDFATYYCQQLNSYFTFGPGTKVDIK
ISG6.149L       ...............................L............
```

**Figure S15**. **Germline-reverted variants of ISG5.143, ISG5.144, ISG5.163 and ISG6.149.** Amino acid sequence alignment of the heavy and light chains and inferred germline-reverted variant.

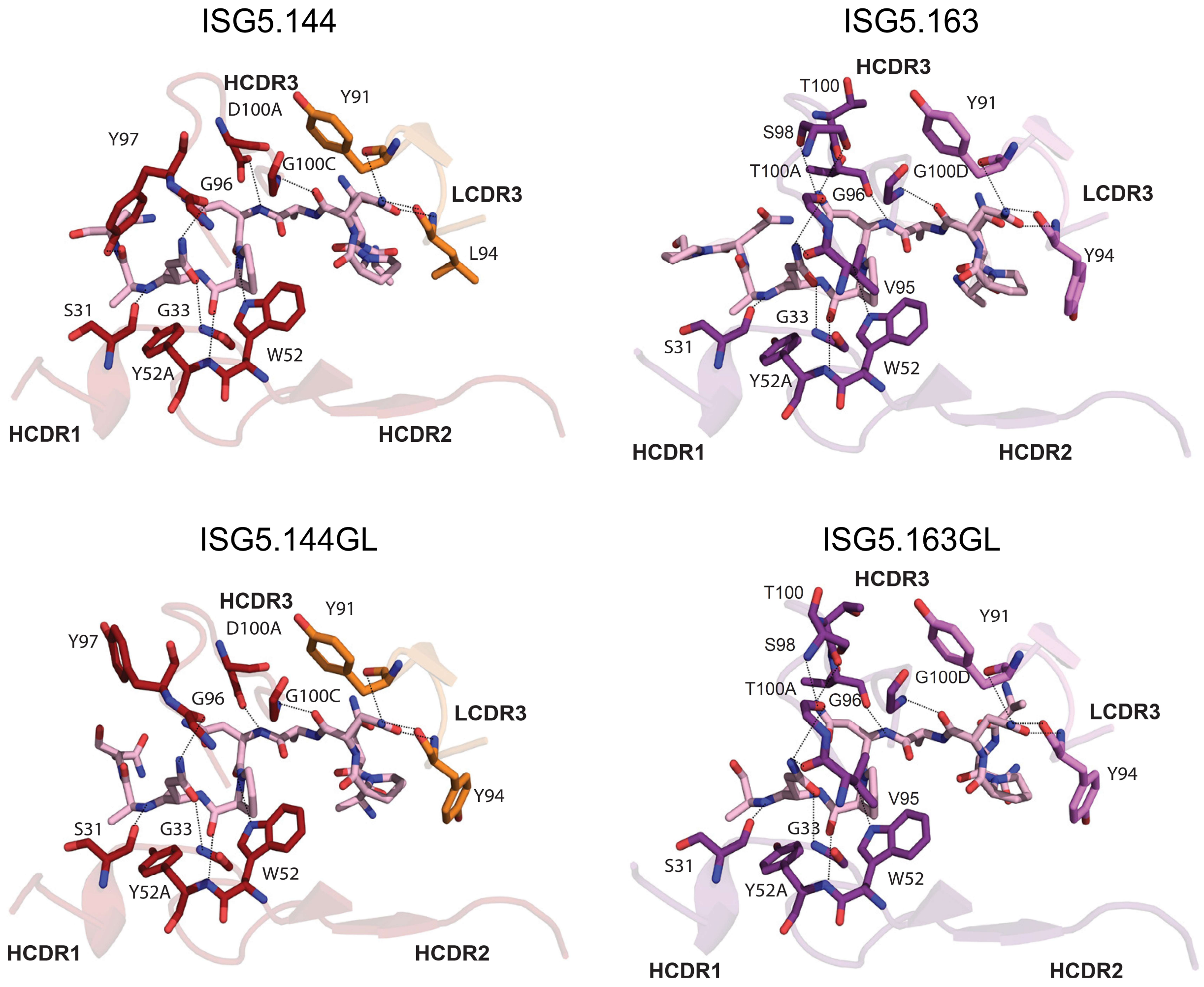


**Figure S16**. **Co-crystal structures of mutated and germline-reverted variants of ISG5.144 and ISG5.163 Fabs**. Comparison of crystal structures of matured (top) and GL-reverted variants (bottom) Fabs bound to NPNA epitope.

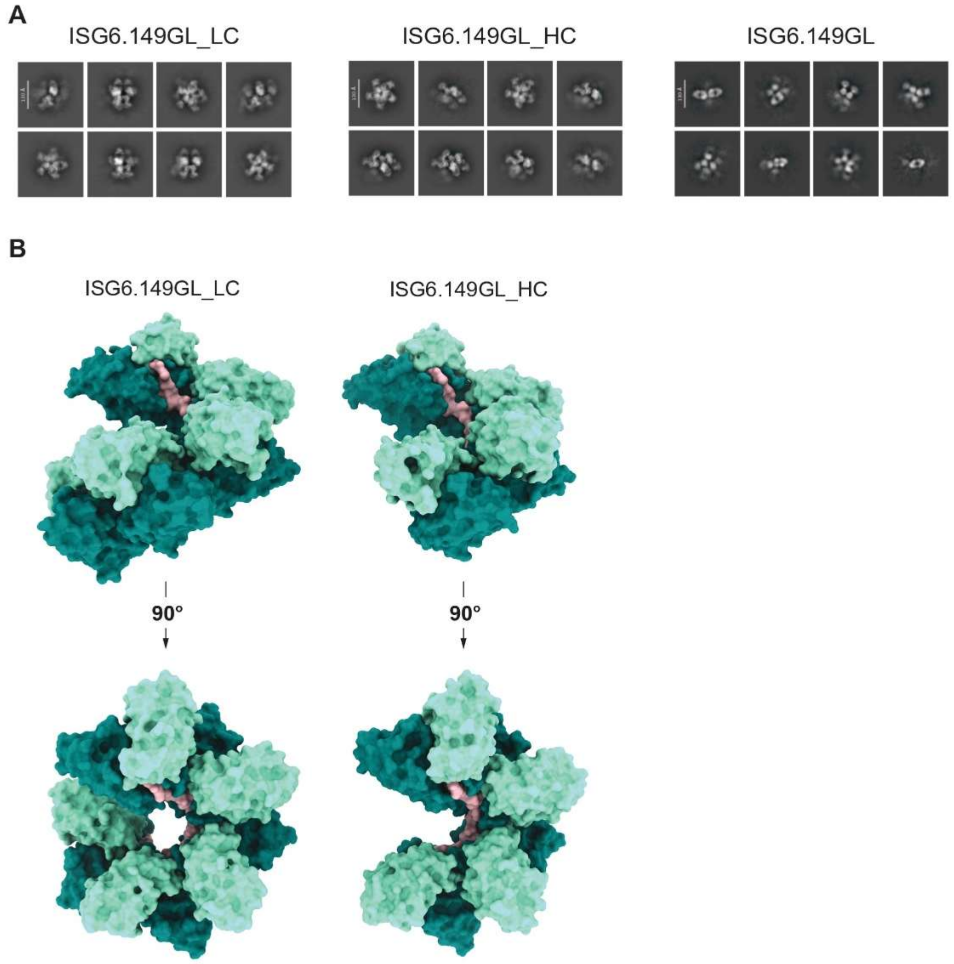


**Figure S17**. **Cryo-EM analysis of GL-reverted variants of ISG6.149**. (**A**) Representative 2D classes of ISG6.149GL, ISG6.149GL_HC and ISG6.149GL_LC Fabs in complex with rsCSP protein. (**B**) Surface molecular representation of rsCSP (pink) in complex with the indicated Fabs, with heavy and light chains colored in teal and aquamarine, respectively. ISG6.149GL did not form homotypic spirals, and 3D reconstruction was not obtained.

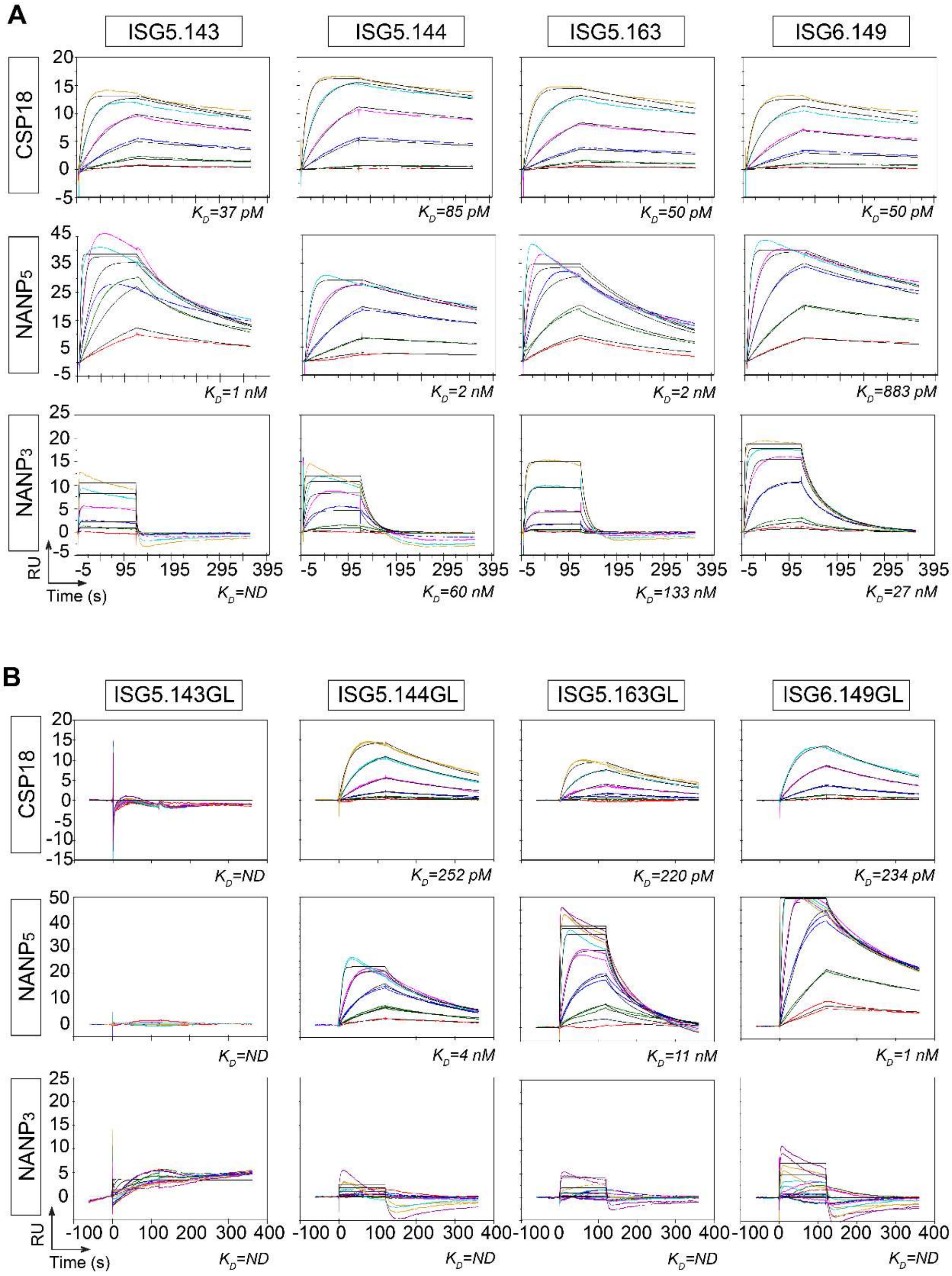


**Figure S18**. **Surface plasmon resonance (SPR) of mutated and germline-reverted antibodies**. Representative multi-cycle kinetics (MCK) sensorgrams of mutated (**A**) and germline-reverted variants (**B**) of ISG5.143, ISG5.144, ISG5.163 and ISG6.149 antibodies, bound to CSP18, $NANP_5$ and $NANP_3$.

**Table S1.** Characteristics of RTS,S vaccinated donors

| Donor ID | Age* | | Gender | Site | CSP18 IgG ELISA (AUC) | | CSP18-reactive IgG$^+$ B cells (%) | n CSP18 mAbs |
|---|---|---|---|---|---|---|---|---|
| | Weeks | Months | | | M0 | M3 | | |
| d3 | 27 | 6 | M | Manhiça | 0.4 | 9.1 | 1.4 | 9 |
| d4 | 9 | 2 | M | Manhiça | 0.5 | 5.8 | 1.2 | 13 |
| d5 | 9 | 2 | F | Manhiça | 0.3 | 7.6 | 0.95 | 16 |
| d6 | 9 | 2 | F | Manhiça | 0.4 | 6.1 | 0.41 | 21 |
| d8 | 38 | 8 | M | Manhiça | 0.3 | 8.2 | 0.82 | 10 |
| d9 | 9 | 2 | F | Manhiça | 0.4 | 7.4 | 0.84 | 19 |

### **Table S2.** Immunoglobulin gene repertoire and reactivity of *Pf*CSP memory B-cell antibodies from RTS,S vaccinated donors

| mAb | Donor | IgH VH | IgH DH | IgH JH | IgH Subtype | CDRH3 Sequence (aa) | CDRH3 Length | CDRH3 (+) | CDRH3 (-) | VH MUT | SHM (%) | IgL | IgL VL | IgL JL | CDRL3 Sequence (aa) | CDRL3 Length | CDRL3 (+) | CDRL3 (-) | VL MUT | SHM (%) | Clonotype members (N) | *Pf*CSP screening (AUC) CSP18 | *Pf*CSP screening (AUC) NANP10 | *Pf*CSP screening (AUC) CTD | *Pf*CSP peptide mapping (AUC) Jt1 | Jt2 | M1 | M2 | NANP3 | NANP4 | NANP5 | NANP6 | NANP10 | Lethality viability 100 µg/ml (%) | Lethality EC50 (µg/ml) | CSPR 100 µg/ml | CSPR 10 µg/ml | Inhibition cell infection (%) 3 µg/ml | Inhibition cell infection (%) 1 µg/ml | ADCP score |
|---|---|---|---|---|---|---|---|---|---|---|---|---|---|---|---|---|---|---|---|---|---|---|---|---|---|---|---|---|---|---|---|---|---|---|---|---|---|---|---|---|
| ISG3.101 | d3 | 3-33*01 | 3-22*01 | 4*02 | y1 | AIDNYDRSDYSTFDY | 15 | 1 | 4 | 2 | 0,7 | κ | 1-5*03 | 1*01 | QQYNSYWT | 8 | 0 | 0 | 6 | 2,3 | 1 | 5,7 | 5,7 | 0,6 | 0,3215 | 6,411 | 0,578 | 12,3 | 10,74 | 11,72 | 12,88 | 13,14 | 13,2 | 9,0 | 8,2 | 1712 | 1659 | 74,2 | 29,9 | 9,5 |
| ISG3.102 | d3 | 3-33*06 | 3-9*01 | 6*03 | y1 | DHDYFAPNYMDV | 12 | 1 | 3 | 8 | 2,7 | κ | 3-11*01 | 2*01 | QQRSNWPPFT | 10 | 1 | 0 | 7 | 2,6 | 1 | 5,7 | 5,5 | 0,2 | 0,343 | 0,272 | 0,2852 | 0,2947 | 0,3294 | 0,2768 | 0,3948 | 3,237 | 9,2 | 50,6 | 100 | 1365 | 1249 | 16,7 | 8,0 | 1,1 |
| ISG3.103 | d3 | 3-21*01 | 3-9*01 | 3*02 | y1 | DFGTELRHFDWSDALDI | 17 | 2 | 5 | 7 | 2,4 | κ | 3-20*01 | 4*01 | QQYGSSLT | 8 | 0 | 0 | 4 | 1,5 | 1 | 5,7 | 0,2 | 5,7 | ND | ND | ND | ND | ND | ND | ND | ND | ND | 75,9 | ND | 1295 | ND | 17,7 | ND | ND |
| ISG3.109 | d3 | 3-23*04 | 7-27*01 | 3*02 | y1 | NTLSGNWGPDAFDI | 14 | 0 | 2 | 13 | 4,5 | κ | 3-11*01 | 4*01 | QQRSRRPT | 8 | 3 | 0 | 8 | 3,0 | 1 | 5,7 | 0,1 | 5,7 | ND | ND | ND | ND | ND | ND | ND | ND | ND | 82,4 | ND | 1268 | ND | 1,3 | ND | ND |
| ISG3.114 | d3 | 3-21*01 | 3-3*01 | 6*03 | y1 | GDSFWSGYSNSRVYYSCMDV | 20 | 1 | 2 | 3 | 1,0 | κ | 3-20*01 | 1*01 | QQYGSSPPWT | 10 | 0 | 0 | 7 | 2,6 | 1 | 5,7 | 0,1 | 5,4 | ND | ND | ND | ND | ND | ND | ND | ND | ND | 80,6 | ND | 1293 | ND | 3,1 | ND | ND |
| ISG3.120 | d3 | 7-4-1*02 | 4-23*01 | 2*01 | y1 | DYGGNSEDWYFDL | 13 | 0 | 4 | 9 | 3,0 | λ | 3-21*03 | 1*01 | QVWDSSSDHGV | 11 | 1 | 2 | 8 | 2,8 | 1 | 5,7 | 0,1 | 5,6 | ND | ND | ND | ND | ND | ND | ND | ND | ND | 80,1 | ND | 1195 | ND | 18,4 | ND | ND |
| ISG3.122 | d3 | 3-33*06 | 6-19*01 | 4*02 | y1 | DGRGLSGVAGTGFFDF | 16 | 1 | 2 | 5 | 1,7 | λ | 1-40*01 | 1*01 | QSYDSSLGAFV | 11 | 0 | 1 | 5 | 1,7 | 1 | 5,7 | 5,7 | 1,4 | 0,3397 | 3,99 | 6,145 | 12,99 | 12,13 | 12,25 | 13,11 | 13,11 | 13,1 | 11,9 | 6,5 | 1568 | 1428 | 92,2 | 33,1 | 15,2 |
| ISG3.126 | d3 | 3-21*01 | 7-27*01 | 5*02 | y1 | DPSSGNWGAWFDP | 13 | 0 | 2 | 6 | 2,0 | λ | 3-21*02 | 2*01 | QVWDTTSDHPGV | 12 | 1 | 2 | 9 | 3,2 | 1 | 5,7 | 0,1 | 5,7 | ND | ND | ND | ND | ND | ND | ND | ND | ND | 83,3 | ND | 1316 | ND | 0 | ND | ND |
| ISG3.136 | d3 | 3-33*06 | 1-7*01 | 4*02 | y1 | VGEGTAERGGAFDY | 14 | 1 | 3 | 9 | 3,1 | κ | 1-5*03 | 1*01 | QQYNSLWT | 8 | 0 | 0 | 10 | 3,8 | 1 | 5,7 | 5,7 | 0,5 | 0,3448 | 8,144 | 10,61 | 13,12 | 12,57 | 12,01 | 13,2 | 13,24 | 13,1 | 10,7 | 5,8 | 1494 | 1554 | 80,0 | 17,0 | 14,6 |
| ISG4.103 | d4 | 4-38-2*01 | 3-22*01 | 5*02 | y1 | AITMVIVVITELHNWFDP | 18 | 1 | 2 | 10 | 3,5 | κ | 3-15*01 | 5*01 | QQYNSLWT | 9 | 0 | 0 | 2 | 1,9 | 1 | 5,7 | 0,3 | 5,7 | ND | ND | ND | ND | ND | ND | ND | ND | ND | 71,9 | ND | 1283 | ND | 19,1 | ND | ND |
| ISG4.106 | d4 | 3-11*04 | 3-22*01 | 4*02 | y1 | GRTNMMVHDTYFDY | 14 | 2 | 2 | 13 | 4,4 | λ | 3-1*01 | 2*01 | QAWDSSTYVV | 10 | 0 | 1 | 6 | 2,2 | 1 | 5,7 | 0,2 | 5,7 | ND | ND | ND | ND | ND | ND | ND | ND | ND | 81,4 | ND | 1230 | ND | 7,2 | ND | ND |
| ISG4.107 | d4 | 3-30*02 | 6-19*01 | 4*02 | y1 | DGGWYYFDY | 9 | 0 | 2 | 8 | 2,7 | κ | 4-1*01 | 2*03 | QQCYSSPYS | 9 | 0 | 0 | 6 | 2,1 | 1 | 3,8 | 3,7 | 0,2 | 0,337 | 0,269 | 0,274 | 0,3062 | 0,3138 | 0,2827 | 0,291 | 2,254 | 4,5 | 77,3 | 100 | 1265 | 1261 | 0 | 12,1 | 1,1 |
| ISG4.109 | d4 | 1-69*18 | 1-7*01 | 4*02 | y1 | GRRGWNFPRYFDY | 13 | 3 | 1 | 10 | 3,4 | κ | 1-33*01 | 5*01 | QQYDNLPSIT | 10 | 0 | 1 | 3 | 1,1 | 5 | 5,7 | 0,4 | 5,7 | ND | ND | ND | ND | ND | ND | ND | ND | ND | 83,0 | ND | 1270 | ND | 21,6 | ND | ND |
| ISG4.110 | d4 | 3-30*02 | 7-27*01 | 3*02 | y1 | NWVEALDI | 8 | 0 | 2 | 17 | 5,8 | κ | 1-39*01 | 2*01 | QQSYSSRPYT | 10 | 1 | 0 | 6 | 2,3 | 1 | 5,7 | 0,2 | 5,7 | ND | ND | ND | ND | ND | ND | ND | ND | ND | 78,0 | ND | 1226 | ND | 0 | ND | ND |
| ISG4.119 | d4 | 3-64D*09 | 5-18*01 | 5*02 | y1 | SATDSFSWYNWFDP | 14 | 0 | 2 | 7 | 2,4 | λ | 1-44*01 | 3*02 | AAWDDSLNGWV | 11 | 0 | 2 | 2 | 0,7 | 1 | 5,7 | 0,2 | 5,7 | ND | ND | ND | ND | ND | ND | ND | ND | ND | 74,1 | ND | 1261 | ND | 12,4 | ND | ND |
| ISG4.124 | d4 | 3-23*04 | 5-18*01 | 4*02 | y1 | GRGYSYGVFDY | 13 | 1 | 1 | 10 | 3,4 | κ | 1-33*01 | 2*01 | QQYDNLPYT | 9 | 0 | 1 | 5 | 1,9 | 1 | 5,7 | 0,1 | 5,7 | ND | ND | ND | ND | ND | ND | ND | ND | ND | 75,6 | ND | 1290 | ND | 0 | ND | ND |
| ISG4.128 | d4 | 3-23*04 | 5-18*01 | 4*02 | y1 | GLGYSYGRFDS | 13 | 1 | 1 | 20 | 6,8 | κ | 1-33*01 | 2*01 | QQYDNLPYT | 9 | 0 | 1 | 5 | 1,9 | 1 | 5,7 | 0,1 | 5,7 | ND | ND | ND | ND | ND | ND | ND | ND | ND | 81,1 | ND | 1306 | ND | 11,6 | ND | ND |
| ISG4.129 | d4 | 5-10-1*01 | 6-13*01 | 5*02 | y1 | HISSSSWVHSSWFDP | 15 | 2 | 1 | 4 | 1,4 | κ | 3-15*01 | 2*01 | QQYNNWPPEMYT | 12 | 0 | 1 | 2 | 0,7 | 1 | 5,7 | 0,2 | 5,1 | ND | ND | ND | ND | ND | ND | ND | ND | ND | 81,4 | ND | 1307 | ND | 3,1 | ND | ND |
| ISG4.135 | d4 | 3-30*02 | 6-19*01 | 4*02 | y1 | VGDGTVAGHFDY | 12 | 1 | 2 | 12 | 4,1 | κ | 2-30*01 | 2*01 | MQGTHWPPT | 9 | 1 | 0 | 1 | 0,4 | 1 | 5,7 | 5,7 | 0,2 | 0,3232 | 0,3209 | 0,2763 | 7,818 | 0,405 | 4,183 | 8,403 | 12,08 | 13,1 | 11,8 | 23,1 | 1588 | 1337 | 68,9 | 40,1 | 16,9 |
| ISG4.157 | d4 | 3-64D*09 | 1-1*01 | 6*02 | y1 | GELERRGYYYNGMDV | 15 | 2 | 3 | 0 | 0,0 | λ | 3-1*01 | 2*01 | QAWDSSTLGVV | 11 | 0 | 1 | 4 | 1,4 | 1 | 5,7 | 0,1 | 5,7 | ND | ND | ND | ND | ND | ND | ND | ND | ND | 82,0 | ND | 1229 | ND | 20,6 | ND | ND |
| ISG4.171 | d4 | 1-2*06 | 6-19*01 | 5*02 | y1 | DVWGGSSGYPFDP | 13 | 0 | 2 | 4 | 1,4 | κ | 2D-29*01 | 1*01 | MQSIQLPRT | 9 | 1 | 0 | 9 | 3,2 | 1 | 5,7 | 5,7 | 0,2 | 0,3106 | 0,2827 | 0,2661 | 2,366 | 0,3167 | 0,4153 | 7,838 | 11,67 | 13,1 | 10,3 | 6,0 | 1480 | 1583 | 96,7 | 23,9 | 3,2 |
| ISG5.108 | d5 | 3-21*01 | 1-7*01 | 4*02 | y3 | PSIMFNWNYYFDY | 13 | 0 | 1 | 4 | 1,4 | κ | 3-15*02 | 5*01 | QQYDNWPPIT | 10 | 0 | 1 | 4 | 1,5 | 1 | 5,7 | 0,3 | 5,7 | ND | ND | ND | ND | ND | ND | ND | ND | ND | 74,8 | ND | 1275 | ND | 0,5 | ND | ND |
| ISG5.110 | d5 | 3-33*01 | 3/OR15-3a*01 | 3*02 | y3 | VRSGPANLVRDWTGAFDI | 18 | 2 | 2 | 5 | 1,7 | κ | 1-5*03 | 1*01 | QQYNSYWT | 8 | 0 | 0 | 7 | 2,6 | 1 | 5,7 | 5,7 | 0,2 | 0,3364 | 1,696 | 2,92 | 12,2 | 11,64 | 13,13 | 13,22 | 13,25 | 13,3 | 10,3 | 13,8 | 1513 | 1381 | 42,4 | 20,6 | 13,3 |
| ISG5.112 | d5 | 3-21*01 | 3-3*01 | 4*02 | y1 | AGNFWSGPYDY | 11 | 0 | 1 | 0 | 0,0 | κ | 3-15*02 | 1*01 | QQYNNWPWT | 9 | 0 | 0 | 2 | 0,8 | 1 | 1,9 | 0,1 | 3,4 | ND | ND | ND | ND | ND | ND | ND | ND | ND | ND | ND | ND | ND | ND | ND | ND |
| ISG5.116 | d5 | 3-33*06 | 2-2*01 | 3*02 | y1 | VGDGGFCSGTYCHGDAFDI | 19 | 1 | 3 | 3 | 1,0 | κ | 3-15*02 | 1*01 | QQYNNWWT | 8 | 0 | 0 | 3 | 1,1 | 1 | 5,6 | 5,7 | 0,2 | 0,3279 | 0,4664 | 0,8733 | 9,045 | 7,09 | 12,68 | 13,11 | 13,21 | 13,3 | 9,7 | 7,2 | 1590 | 1587 | 80,3 | 28,8 | 13,8 |
| ISG5.118 | d5 | 3-73*02 | 2-15*01 | 5*02 | y1 | AVVGQDWFGP | 10 | 0 | 1 | 12 | 4,0 | λ | 3-9*01 | 2*01 | QVWDSSTHVV | 10 | 1 | 1 | 5 | 1,8 | 1 | 5,7 | 0,1 | 5,7 | ND | ND | ND | ND | ND | ND | ND | ND | ND | 82,2 | ND | 1254 | ND | 0 | ND | ND |
| ISG5.130 | d5 | 3-33*06 | 2-2*01 | 3*01 | y1 | VGDGGYCSSTSCHGDAFDL | 19 | 1 | 3 | 0 | 0,0 | κ | 3-15*02 | 1*01 | QQYNNWWT | 8 | 0 | 0 | 7 | 2,7 | 3 | 5,7 | 5,7 | 0,2 | 0,3381 | 1,053 | 1,915 | 11,02 | 6,699 | 11,89 | 12,98 | 13,12 | 13,2 | 9,2 | 11,8 | 1651 | 1590 | 60,6 | 20,6 | 14,1 |
| ISG5.138 | d5 | 3-33*01 | 1-7*01 | 3*02 | y1 | VCNEACQGKGTTSLAFDI | 18 | 1 | 2 | 5 | 1,7 | κ | 3-15*02 | 1*01 | QQYNNWWT | 8 | 0 | 0 | 6 | 2,3 | 1 | 5,7 | 5,7 | 0,2 | 0,3343 | 10,14 | 9,981 | 12,68 | 12,37 | 13,12 | 13,22 | 13,26 | 13,1 | 7,5 | 6,3 | 1630 | 1469 | 77,6 | 12,9 | 9,8 |
| ISG5.143 | d5 | 3-33*01 | 2-15*01 | 3*02 | y1 | DRRYCSGGNCYSGAFDI | 17 | 2 | 2 | 9 | 3,0 | κ | 1-5*03 | 1*01 | QQYYSYWT | 8 | 0 | 0 | 1 | 0,4 | 1 | 5,7 | 5,7 | 0,2 | 0,3254 | 0,2958 | 0,2751 | 6,23 | 6,241 | 12,61 | 13,08 | 13,17 | 13,3 | 9,6 | 6,5 | 1741 | 1667 | 91,2 | 32,9 | 11,2 |
| ISG5.144 | d5 | 3-33*06 | 5-12*01 | 4*02 | y1 | VGYSGYDWGGAFDY | 14 | 0 | 2 | 2 | 0,7 | κ | 1-5*03 | 1*01 | QQYNSLWT | 8 | 0 | 0 | 6 | 2,3 | 1 | 5,7 | 5,7 | 0,2 | 0,3369 | 5,98 | 6,196 | 12,74 | 11,83 | 12,54 | 13,15 | 13,1 | 13,2 | 9,3 | 12,0 | 1530 | 1562 | 68,7 | 7,7 | 12,7 |
| ISG5.145 | d5 | 3-21*01 | 6-19*01 | 4*02 | y1 | RWGIAVRYFDY | 11 | 2 | 1 | 8 | 2,7 | λ | 3-21*03 | 2*01 | QVWDSSSDHPDVV | 13 | 1 | 3 | 4 | 1,4 | 1 | 5,7 | 0,2 | 5,7 | ND | ND | ND | ND | ND | ND | ND | ND | ND | 77,1 | ND | 1273 | ND | 0 | ND | ND |
| ISG5.147 | d5 | 3-33*06 | 2-15*01 | 4*02 | y1 | GGQWGSGEFDY | 11 | 0 | 2 | 0 | 0,0 | κ | 4-1*01 | 1*01 | QQYYSTPWT | 9 | 0 | 0 | 1 | 0,4 | 1 | 3,6 | 2,3 | 0,2 | 0,3369 | 0,2553 | 0,3479 | 0,2916 | 0,3029 | 0,4069 | 0,5419 | 0,5957 | 3,8 | 79,4 | 100 | 1310 | 1263 | 0 | 3,4 | 1,3 |
| ISG5.154 | d5 | 3-33*06 | 6-13*01 | 4*02 | y1 | VGSSSWYYFDY | 11 | 0 | 1 | 0 | 0,0 | λ | 6-57*01 | 3*02 | QSYDWV | 6 | 0 | 1 | 0 | 0,0 | 1 | 5,7 | 5,7 | 0,2 | 0,3197 | 0,2588 | 0,2846 | 0,2854 | 0,3158 | 0,3197 | 4,453 | 6,71 | 10,9 | 78,6 | 100 | 1286 | 1260 | 0 | 13,3 | 1,0 |
| ISG5.160 | d5 | 3-21*01 | 2-2*01 | 3*02 | y1 | DRGRYCSTTVCYDAFDI | 17 | 2 | 3 | 6 | 2,0 | κ | 3-20*01 | 2*01 | QQYGSSRYTT | 10 | 1 | 0 | 4 | 1,5 | 1 | 5,7 | 0,1 | 5,7 | ND | ND | ND | ND | ND | ND | ND | ND | ND | 82,1 | ND | 1256 | ND | 0 | ND | ND |
| ISG5.163 | d5 | 3-33*01 | 1-1*01 | 3*02 | y1 | VGTSGTTERGGAFDI | 15 | 1 | 2 | 4 | 1,4 | κ | 1-5*03 | 1*01 | QQYNNYWT | 8 | 0 | 0 | 2 | 0,8 | 1 | 5,7 | 5,7 | 0,2 | 0,3269 | 10,61 | 12,29 | 13,01 | 12,9 | 12,36 | 13 | 12,99 | 13,4 | 11,5 | 8,0 | 1680 | 1700 | 80,4 | 20,1 | 8,9 |
| ISG5.166 | d5 | 3-7*01 | N/A | 4*02 | y1 | YLNY | 4 | 0 | 0 | 4 | 1,4 | κ | 2D-29*02 | 5*01 | MQSIQLPIT | 9 | 0 | 0 | 2 | 0,7 | 1 | 5,7 | 5,7 | 0,2 | 12,57 | 0,2839 | 0,3118 | 10,62 | 10 | 12,04 | 13,12 | 13,12 | 13,3 | 12,0 | 16,0 | 1302 | 1302 | 22,8 | 33,6 | 2,6 |
| ISG5.167 | d5 | 3-33*01 | 4-17*01 | 3*02 | y1 | VQADNDYVAFDI | 12 | 0 | 3 | 11 | 3,7 | κ | 1-5*03 | 1*01 | QQYNSYWT | 8 | 0 | 0 | 3 | 1,2 | 2 | 5,7 | 5,7 | 0,2 | 0,3112 | 0,2743 | 0,3052 | 11,17 | 11,29 | 11,62 | 12,97 | 13,04 | 13,2 | 12,2 | 5,9 | 1400 | 1641 | 91,5 | 23,7 | 8,2 |
| ISG5.170 | d5 | 4-39*07 | 6-19*01 | 1*01 | y1 | EEREILAGTVMFSRYFQH | 18 | 3 | 3 | 12 | 4,0 | λ | 1-47*01 | 2*01 | AAWDDSLSGVV | 11 | 0 | 2 | 3 | 1,0 | 1 | 5,7 | 0,1 | 5,7 | ND | ND | ND | ND | ND | ND | ND | ND | ND | 78,3 | ND | 1316 | ND | 14,9 | ND | ND |
| ISG5.174 | d5 | 3-48*01 | 6-13*01 | 4*02 | y1 | LGLIAAAGTRYFDY | 14 | 1 | 1 | 4 | 1,4 | λ | 3-21*01 | 2*01 | QVWDSGSDHPV | 11 | 1 | 2 | 10 | 3,5 | 1 | 5,7 | 0,2 | 5,7 | ND | ND | ND | ND | ND | ND | ND | ND | ND | 82,8 | ND | 1304 | ND | 15,1 | ND | ND |
| ISG5.179 | d5 | 3-33*06 | 3-22*01 | 3*02 | y1 | VGSSESSGYYKPTEGWTDAFDI | 22 | 1 | 4 | 2 | 0,7 | κ | 1-5*03 | 1*01 | QQYNFLWT | 8 | 0 | 0 | 2 | 0,8 | 1 | 5,7 | 5,7 | 0,2 | 0,3313 | 3,29 | 1,003 | 10,88 | 10,9 | 13,13 | 13,27 | 13,29 | 13,3 | 9,7 | 17,9 | 1562 | 1399 | 52,0 | 9,2 | 5,3 |
| ISG5.184 | d5 | 3-48*01 | 6-13*01 | 4*02 | y1 | LGSSWTYYFDY | 11 | 0 | 1 | 8 | 2,7 | λ | 3-21*03 | 2*01 | QVWDTTSDLVV | 11 | 0 | 2 | 5 | 1,8 | 1 | 5,7 | 0,2 | 5,7 | ND | ND | ND | ND | ND | ND | ND | ND | ND | 81,2 | ND | 1200 | ND | 17,5 | ND | ND |
| ISG6.102 | d6 | 3-21*01 | 3-16*01 | 4*02 | y1 | DDLGVRGLGDYFDY | 14 | 1 | 4 | 7 | 2,4 | λ | 3-1*01 | 1*01 | QAWDSSTYV | 9 | 0 | 1 | 9 | 3,2 | 1 | 5,7 | 0,1 | 5,7 | ND | ND | ND | ND | ND | ND | ND | ND | ND | 81,7 | ND | 1369 | ND | 5,0 | ND | ND |
| ISG6.104 | d6 | 7-4-1*02 | 1-26*01 | 6*02 | y1 | VRWELLGYYYGMDV | 14 | 1 | 2 | 1 | 0,3 | κ | 4-1*01 | 4*01 | QQYYITPPLT | 10 | 0 | 0 | 2 | 0,7 | 1 | 5,7 | 5,7 | 0,2 | 0,3279 | 12,78 | 4,18 | 13,25 | 8,83 | 12,91 | 13,25 | 13,25 | 13,3 | 9,1 | 29,5 | 1456 | 1334 | 26,3 | 18,4 | 7,5 |
| ISG6.106 | d6 | 3-30*01 | 2-2*01 | 4*02 | y1 | GCSTSCVLGDRFDY | 14 | 1 | 2 | 0 | 0,0 | κ | 3-11*01 | 1*01 | QQHSNWPRT | 9 | 2 | 0 | 3 | 1,1 | 1 | 5,7 | 5,7 | 0,2 | 0,3276 | 10,91 | 9,425 | 12,99 | 4,367 | 11,13 | 12,91 | 13,02 | 13,2 | 9,5 | 19,3 | 1360 | 1291 | 18,7 | 11,1 | 1,8 |
| ISG6.111 | d6 | 3-48*04 | 3-3*01 | 4*02 | y1 | RIFGVVQYYFDY | 12 | 1 | 1 | 4 | 1,4 | λ | 3-1*01 | 1*01 | QAWDSRTAYV | 10 | 1 | 1 | 5 | 1,8 | 1 | 5,7 | 0,1 | 5,7 | ND | ND | ND | ND | ND | ND | ND | ND | ND | 83,9 | ND | 1297 | ND | 4,7 | ND | ND |
| ISG6.118 | d6 | 3-33*06 | 3-22*01 | 5*02 | y2 | DLRYFYDISDTGGFDP | 16 | 1 | 4 | 5 | 1,7 | λ | 1-40*01 | 1*01 | QSYDSRLRGYV | 11 | 2 | 1 | 4 | 1,3 | 1 | 5,7 | 5,7 | 0,7 | 0,4672 | 11 | 12,38 | 13,28 | 11,61 | 13,3 | 13,32 | 13,32 | 13,1 | 10,2 | 6,9 | 1575 | 1449 | 94,7 | 22,3 | 13,8 |
| ISG6.139 | d6 | 3-33*06 | 3/OR15-3a*01 | 4*02 | y1 | SASVFGHGYYFDY | 13 | 1 | 1 | 8 | 2,7 | κ | 3-11*01 | 1*01 | QQRNNWPPT | 9 | 1 | 0 | 5 | 1,9 | 1 | 5,7 | 5,7 | 0,2 | 0,3275 | 0,3352 | 0,4301 | 11,61 | 5,59 | 10,38 | 12,89 | 12,96 | 13,1 | 12,1 | 24,3 | 1308 | 1276 | 5,8 | 12,1 | 1,2 |
| ISG6.141 | d6 | 3-21*01 | 4-11*01 | 5*02 | y3 | RPGVTTTDNWFDP | 13 | 1 | 2 | 12 | 4,1 | λ | 3-21*03 | 1*01 | QVWDSSSDYV | 10 | 0 | 2 | 2 | 0,7 | 1 | 5,7 | 0,1 | 5,7 | ND | ND | ND | ND | ND | ND | ND | ND | ND | 82,9 | ND | 1296 | ND | 3,4 | ND | ND |
| ISG6.149 | d6 | 3-33*01 | 1-26*01 | 2*01 | y1 | PSLTWESSRYFDL | 13 | 1 | 2 | 4 | 1,4 | κ | 1-9*01 | 3*01 | QQLNSLFT | 8 | 0 | 0 | 4 | 1,6 | 2 | 5,7 | 5,7 | 0,2 | 0,3263 | 0,2827 | 0,315 | 11,6 | 10,3 | 10,78 | 12,92 | 13,06 | 13,2 | 11,4 | 5,3 | 1755 | 1402 | 92,5 | 43,7 | 9,9 |
| ISG6.152 | d6 | 3-21*01 | 4-23*01 | 2*01 | y1 | AGGLRWIDHWYFDL | 14 | 2 | 2 | 5 | 1,7 | λ | 1-51*02 | 3*02 | GTWDSSLSAGWV | 12 | 0 | 1 | 4 | 1,4 | 1 | 1,2 | 0,1 | 2,2 | ND | ND | ND | ND | ND | ND | ND | ND | ND | ND | ND | ND | ND | ND | ND | ND |
| ISG6.158 | d6 | 3-21*01 | 3-3*01 | 4*02 | y1 | VSGVVIRHFDY | 11 | 2 | 1 | 6 | 2,0 | λ | 3-21*03 | 2*01 | QVWDSSSDHHVV | 12 | 2 | 2 | 4 | 1,4 | 1 | 5,7 | 0,1 | 5,7 | ND | ND | ND | ND | ND | ND | ND | ND | ND | 78,6 | ND | 1305 | ND | 10,3 | ND | ND |
| ISG6.160 | d6 | 3-33*01 | N/A | 4*02 | y1 | PIGGHYFDY | 9 | 1 | 1 | 1 | 0,3 | κ | 2-29*03 | 1*01 | MQGIHLWT | 8 | 1 | 0 | 5 | 1,8 | 1 | 5,7 | 5,7 | 0,2 | 0,3278 | 1,635 | 9,523 | 13,02 | 7,331 | 12,1 | 13,15 | 13,21 | 13,0 | 8,3 | 11,6 | 1310 | 1229 | 37,3 | 22,2 | 1,2 |
| ISG6.163 | d6 | 3-48*04 | 1-7*01 | 4*02 | y1 | DGLYKWNYYFDY | 12 | 1 | 2 | 6 | 2,0 | λ | 3-21*03 | 3*02 | QVWDSSSDHRV | 11 | 2 | 2 | 1 | 0,4 | 1 | 5,7 | 0,1 | 5,7 | ND | ND | ND | ND | ND | ND | ND | ND | ND | 77,0 | ND | 1309 | ND | 0 | ND | ND |
| ISG6.165 | d6 | 3-33*01 | 6-6*01 | 4*02 | y1 | GVWAYSSASDY | 11 | 0 | 1 | 4 | 1,4 | κ | 1-9*01 | 2*01 | QQLNSYPYT | 9 | 0 | 0 | 6 | 2,2 | 1 | 5,5 | 0,1 | 5,6 | ND | ND | ND | ND | ND | ND | ND | ND | ND | 82,4 | ND | 1241 | ND | 11,9 | ND | ND |
| ISG6.166 | d6 | 1-8*01 | 3-3*01 | 6*02 | y1 | GYDFWSGYSRSRNYYYVMDV | 20 | 2 | 2 | 1 | 0,3 | κ | 3-20*01 | 5*01 | QQYGSSPPEIT | 11 | 0 | 1 | 2 | 0,7 | 1 | 5,7 | 0,1 | 5,3 | ND | ND | ND | ND | ND | ND | ND | ND | ND | 81,2 | ND | 1306 | ND | 10,6 | ND | ND |
| ISG6.167 | d6 | 3-33*01 | 1-26*01 | 3*02 | y1 | DRGRGSHGGAFDI | 13 | 3 | 2 | 5 | 1,7 | κ | 3-11*01 | 1*01 | QQHSNWWT | 8 | 1 | 0 | 5 | 1,9 | 2 | 5,7 | 5,7 | 0,2 | 0,397 | 5,742 | 12,16 | 13,16 | 11,91 | 12,71 | 13,24 | 13,27 | 13,3 | 10,3 | 20,6 | 1323 | 1460 | 14,4 | 1,0 | 1,8 |
| ISG6.169 | d6 | 3-21*01 | 2-21*02 | 4*02 | y1 | GGGGSDSGSYYFDY | 14 | 0 | 2 | 6 | 2,0 | κ | 3-11*01 | 4*01 | QQRSNWPPLT | 10 | 1 | 0 | 2 | 0,8 | 1 | 5,7 | 0,1 | 5,7 | ND | ND | ND | ND | ND | ND | ND | ND | ND | 84,0 | ND | 1288 | ND | 9,7 | ND | ND |
| ISG6.170 | d6 | 3-33*01 | 6-13*01 | 3*02 | y1 | VRNGYSSSWGDAFDI | 15 | 1 | 2 | 0 | 0,0 | κ | 1-5*03 | 1*01 | QQYNSLWT | 8 | 0 | 0 | 4 | 1,5 | 1 | 5,7 | 5,7 | 0,2 | 0,3212 | 0,2672 | 0,2686 | 1,144 | 5,448 | 8,209 | 11,24 | 11,98 | 13,0 | 24,9 | 74,4 | 1305 | 1262 | 16,4 | 0 | 1,7 |
| ISG6.181 | d6 | 3-21*01 | 6-19*01 | 3*02 | y3 | VSGGRGWFGAFDI | 13 | 1 | 1 | 7 | 2,4 | λ | 3-21*03 | 3*02 | QVWDNSDHPV | 10 | 1 | 2 | 9 | 3,2 | 1 | 5,7 | 0,1 | 5,7 | ND | ND | ND | ND | ND | ND | ND | ND | ND | 82,1 | ND | 1349 | ND | 15,8 | ND | ND |
| ISG6.188 | d6 | 3-21*01 | 6-19*01 | 4*02 | y1 | GWDRGWQRGVDY | 12 | 2 | 2 | 6 | 2,0 | λ | 1-51*01 | 2*01 | GTWDSSLSAVV | 11 | 0 | 1 | 5 | 1,7 | 1 | 5,7 | 0,1 | 5,7 | ND | ND | ND | ND | ND | ND | ND | ND | ND | 81,0 | ND | 1245 | ND | 17,5 | ND | ND |
| ISG6.193 | d6 | 3-15*01 | 1-26*01 | 5*02 | y3 | DYVYSGSYYWFDP | 13 | 0 | 2 | 0 | 0,0 | κ | 2-28*01 | 2*01 | MQALQTPHT | 9 | 1 | 0 | 0 | 0,0 | 1 | 3,2 | 1,7 | 0,2 | 0,3235 | 0,291 | 0,2863 | 0,3549 | 0,302 | 0,4027 | 0,3495 | 0,4427 | 7,9 | 61,6 | 100 | 1276 | 1262 | 15,1 | 2,0 | 1,2 |
| ISG6.195 | d6 | 3-48*04 | 4-23*01 | 4*02 | y1 | ATSTVVTLDLIDY | 13 | 0 | 2 | 10 | 3,4 | κ | 1-5*03 | 1*01 | QHYNSYPWT | 9 | 1 | 0 | 2 | 0,8 | 1 | 3,8 | 0,1 | 4,6 | ND | ND | ND | ND | ND | ND | ND | ND | ND | 80,8 | ND | 1326 | ND | 9,2 | ND | ND |
| ISG6.196 | d6 | 3-21*01 | 1-1*01 | 4*02 | y3 | VGGPWTYYFDY | 11 | 0 | 1 | 5 | 1,7 | λ | 3-21*03 | 2*01 | QVWDSSSNI | 9 | 0 | 1 | 4 | 1,4 | 1 | 5,7 | 0,1 | 5,7 | ND | ND | ND | ND | ND | ND | ND | ND | ND | 75,7 | ND | 1281 | ND | 6,6 | ND | ND |
| ISG8.106 | d8 | 4-39*01 | 1-26*01 | 3*02 | y1 | HIGGSHGAAFDI | 12 | 2 | 1 | 4 | 1,3 | κ | 1-33*01 | 2*01 | QQFDNLPYT | 9 | 0 | 1 | 5 | 2,3 | 1 | 4,4 | 0,1 | 5,4 | ND | ND | ND | ND | ND | ND | ND | ND | ND | 83,3 | ND | 1329 | ND | 10,3 | ND | ND |
| ISG8.108 | d8 | 3-21*01 | 3-3*01 | 5*02 | y1 | DLNDFWSGYQRSGWFDP | 17 | 1 | 3 | 7 | 2,4 | κ | 3-11*01 | 2*01 | QQRSNWSYT | 9 | 1 | 0 | 7 | 1,9 | 1 | 5,7 | 0,1 | 5,7 | ND | ND | ND | ND | ND | ND | ND | ND | ND | 81,4 | ND | 1241 | ND | 0 | ND | ND |
| ISG8.109 | d8 | 4-38-2*01 | 3-10*01 | 4*02 | y1 | HGLILRGGSGSGFDY | 15 | 2 | 1 | 9 | 3,0 | λ | 3-21*03 | 3*02 | QVWDSSSDHWV | 11 | 1 | 2 | 4 | 1,4 | 1 | 5,6 | 0,1 | 5,7 | ND | ND | ND | ND | ND | ND | ND | ND | ND | 81,0 | ND | 1230 | ND | 0 | ND | ND |
| ISG8.113 | d8 | 3-30*02 | 4-23*01 | 3*02 | y1 | NYVDNFDI | 8 | 0 | 2 | 7 | 2,4 | κ | 3-11*01 | 4*01 | HQRSNWPPT | 9 | 2 | 0 | 5 | 1,9 | 1 | 5,6 | 0,1 | 5,6 | ND | ND | ND | ND | ND | ND | ND | ND | ND | 74,4 | ND | 1263 | ND | 3,4 | ND | ND |
| ISG8.117 | d8 | 3-30*02 | 3-16*01 | 3*02 | y1 | NYVDAFDI | 8 | 0 | 2 | 6 | 2,0 | κ | 1-39*01 | 2*03 | QQSYSTPYS | 9 | 0 | 0 | 7 | 2,6 | 1 | 5,7 | 0,1 | 5,7 | ND | ND | ND | ND | ND | ND | ND | ND | ND | 76,9 | ND | 1328 | ND | 0 | ND | ND |
| ISG8.119 | d8 | 4-61*02 | 2-2*02 | 6*03 | y1 | ETVVPADIAVRGYYFYYYMDV | 21 | 1 | 3 | 3 | 1,0 | λ | 1-36*01 | 1*01 | AAWDDSLSGYV | 11 | 0 | 2 | 1 | 0,3 | 1 | 5,7 | 0,1 | 5,7 | ND | ND | ND | ND | ND | ND | ND | ND | ND | 81,6 | ND | 1232 | ND | 20,4 | ND | ND |
| ISG8.120 | d8 | 3-21*01 | 3-3*02 | 4*02 | y1 | RSGFSGAEDYHLGY | 14 | 2 | 2 | 2 | 0,7 | λ | 3-1*01 | 1*01 | QAWDSSTYV | 9 | 0 | 1 | 3 | 1,1 | 1 | 5,7 | 0,1 | 5,7 | ND | ND | ND | ND | ND | ND | ND | ND | ND | 70,3 | ND | 1256 | ND | 8,1 | ND | ND |
| ISG8.131 | d8 | 4-61*02 | 3-22*01 | 4*02 | y1 | AHYDSSYYSLGSDY | 14 | 1 | 2 | 6 | 2,0 | λ | 3-25*02 | 1*01 | QSADSSGTYYV | 11 | 0 | 1 | 4 | 1,4 | 1 | 5,7 | 5,7 | 0,2 | 12,2 | 7,592 | 11,88 | 13,07 | 8,572 | 12,48 | 13,13 | 13,2 | 13,1 | 11,1 | 17,5 | 1499 | 1384 | 38,9 | 6,8 | 10,0 |
| ISG8.132 | d8 | 3-49*04 | 1-1*01 | 4*02 | y1 | VGVHLWPLDY | 10 | 1 | 1 | 10 | 3,3 | κ | 3-20*01 | 1*01 | QQYGDSPWT | 9 | 0 | 1 | 10 | 3,7 | 1 | 5,7 | 5,7 | 0,2 | 0,3411 | 0,272 | 0,2829 | 2,013 | 1,015 | 3,162 | 10,85 | 12,67 | 13,3 | 28,0 | 28,1 | 1538 | 1379 | 18,4 | 0,0 | 2,7 |
| ISG8.133 | d8 | 3-21*01 | 2-2*01 | 6*03 | y3 | DGQLLRIHGLHYYYFYMDV | 19 | 3 | 2 | 0 | 0,0 | λ | 1-51*01 | 3*02 | GTWDSSLRGV | 10 | 1 | 1 | 3 | 1,0 | 1 | 5,7 | 0,1 | 5,6 | ND | ND | ND | ND | ND | ND | ND | ND | ND | 82,0 | ND | 1253 | ND | 15,0 | ND | ND |
| ISG8.137 | d8 | 3-23*04 | 2-2*02 | 5*02 | y3 | DGELCSRTNCYTGWFDP | 17 | 1 | 3 | 1 | 0,3 | λ | 4-69*01 | 3*02 | QTWGTGIRV | 9 | 1 | 0 | 6 | 2,0 | 1 | 5,7 | 0,1 | 5,7 | ND | ND | ND | ND | ND | ND | ND | ND | ND | 77,5 | ND | 1291 | ND | ND | ND | ND |
| ISG8.140 | d8 | 3-21*01 | 3-3*01 | 4*02 | y1 | EERFLESLSIDY | 12 | 1 | 4 | 10 | 3,4 | λ | 3-1*01 | 2*01 | QAWDSSTVV | 9 | 0 | 1 | 8 | 1,4 | 1 | 5,7 | 0,1 | 5,7 | ND | ND | ND | ND | ND | ND | ND | ND | ND | 83,6 | ND | 1302 | ND | 0 | ND | ND |
| ISG8.141 | d8 | 4-39*07 | 7-27*01 | 3*02 | y1 | SIGGLGMAFDI | 11 | 0 | 1 | 6 | 2,0 | κ | 1-33*01 | 4*01 | QHFDNLPLT | 9 | 1 | 1 | 4 | 1,6 | 1 | 5,7 | 0,1 | 5,6 | ND | ND | ND | ND | ND | ND | ND | ND | ND | 77,3 | ND | 1295 | ND | 0 | ND | ND |
| ISG8.143 | d8 | 3-30-3*01 | 5-12*01 | 4*02 | y1 | DGYSSSFFDY | 10 | 0 | 2 | 12 | 4,1 | κ | 1-5*03 | 4*01 | QHYNSYSGLT | 10 | 1 | 0 | 6 | 2,3 | 1 | 5,7 | 5,7 | 0,2 | 0,3127 | 0,2684 | 0,2803 | 0,357 | 0,4443 | 13,04 | 13,32 | 13,33 | 13,3 | 6,7 | 6,5 | 1366 | 1343 | 62,7 | 7,3 | 11,9 |
| ISG8.151 | d8 | 1-2*02 | 6-6*01 | 4*02 | y1 | LTTTARQRDY | 10 | 2 | 1 | 15 | 5,1 | κ | 2-28*01 | 5*01 | MQALQTPIT | 9 | 0 | 0 | 3 | 1,1 | 2 | 5,7 | 5,7 | 0,2 | 0,3274 | 0,3089 | 0,2921 | 8,17 | 9,399 | 12,88 | 13,31 | 13,33 | 13,4 | 7,4 | 7,8 | 1501 | 1464 | 93,9 | 16,7 | 13,5 |
| ISG8.154 | d8 | 3-21*01 | 1-26*01 | 4*02 | y1 | DSLFKWDHHFDY | 12 | 3 | 3 | 5 | 1,7 | λ | 1-44*01 | 1*01 | ATWDDSLNGQV | 11 | 0 | 2 | 4 | 1,4 | 1 | 5,7 | 0,1 | 5,7 | ND | ND | ND | ND | ND | ND | ND | ND | ND | 83,6 | ND | 1332 | ND | 0 | ND | ND |
| ISG9.101 | d9 | 4-61*02 | 2-2*02 | 6*03 | y1 | EEPVVPAAIRYYFYYYMDV | 19 | 1 | 3 | 1 | 0,3 | λ | 1-36*01 | 1*01 | AAWDDRLNAYV | 11 | 1 | 2 | 3 | 1,0 | 1 | 5,7 | 0,2 | 5,7 | ND | ND | ND | ND | ND | ND | ND | ND | ND | 82,7 | ND | 1299 | ND | 0 | ND | ND |
| ISG9.104 | d9 | 4-61*02 | 5-18*01 | 6*03 | y1 | ETADTAMVSYYFYYYMDV | 18 | 0 | 3 | 5 | 1,7 | λ | 1-44*01 | 1*01 | AAWDDSLNGYV | 11 | 0 | 2 | 2 | 0,7 | 1 | 5,7 | 0,1 | 5,7 | ND | ND | ND | ND | ND | ND | ND | ND | ND | 76,1 | ND | 1267 | ND | 1,3 | ND | ND |
| ISG9.105 | d9 | 4-61*02 | 3-22*01 | 6*03 | y1 | ERNYYDTSGYYSHYYYMDV | 19 | 2 | 3 | 4 | 1,3 | λ | 1-47*01 | 1*01 | AAWDDSLSGYV | 11 | 0 | 2 | 7 | 2,4 | 1 | 5,7 | 0,1 | 5,7 | ND | ND | ND | ND | ND | ND | ND | ND | ND | 75,4 | ND | 1267 | ND | 0 | ND | ND |
| ISG9.108 | d9 | 4-61*02 | 6-19*01 | 4*02 | y1 | AEYSSGWYSGGDDS | 14 | 0 | 3 | 9 | 3,0 | λ | 3-25*02 | 3*02 | QSADSSGIYWV | 11 | 0 | 1 | 10 | 3,5 | 2 | 5,7 | 5,7 | 4,2 | 11,66 | 12,2 | 12,53 | 13,13 | 11,89 | 13,12 | 13,25 | 13,29 | 13,1 | 17,2 | 42,2 | 1354 | 1268 | 24,1 | 12,8 | 4,5 |
| ISG9.110 | d9 | 4-61*10 | 1-26*01 | 4*02 | y1 | LVGRTFFDY | 9 | 1 | 1 | 5 | 1,7 | κ | 1-33*01 | 2*01 | QQYDNLPYT | 9 | 0 | 1 | 3 | 1,2 | 1 | 5,7 | 0,1 | 5,7 | ND | ND | ND | ND | ND | ND | ND | ND | ND | 81,6 | ND | 1260 | ND | 0,3 | ND | ND |
| ISG9.123 | d9 | 3-48*03 | 5-18*01 | 3*01 | y1 | MPGGYSFGFHAFDV | 14 | 1 | 1 | 12 | 4,1 | κ | 4-1*01 | 3*01 | QQYYSTPFT | 9 | 0 | 0 | 4 | 1,4 | 1 | 5,7 | 0,1 | 5,6 | ND | ND | ND | ND | ND | ND | ND | ND | ND | 83,0 | ND | 1273 | ND | 10,6 | ND | ND |
| ISG9.128 | d9 | 3-74*01 | 3-10*03 | 4*02 | y1 | SGLDY | 5 | 0 | 1 | 12 | 4,1 | κ | 3-11*01 | 3*01 | QQHSNWPFT | 9 | 1 | 0 | 9 | 3,4 | 1 | 5,7 | 0,1 | 5,7 | ND | ND | ND | ND | ND | ND | ND | ND | ND | 76,5 | ND | 1289 | ND | 11,6 | ND | ND |
| ISG9.129 | d9 | 3-48*03 | 5-18*01 | 3*02 | y1 | LPGGYSYGFHAFDI | 14 | 1 | 1 | 13 | 4,4 | κ | 4-1*01 | 3*01 | QQYYSTPFT | 9 | 0 | 0 | 8 | 2,8 | 1 | 5,7 | 0,2 | 5,7 | ND | ND | ND | ND | ND | ND | ND | ND | ND | 76,5 | ND | 1294 | ND | 3,4 | ND | ND |
| ISG9.145 | d9 | 3-30*02 | 3-9*01 | 4*02 | y1 | NLVDNFDL | 8 | 0 | 2 | 17 | 5,8 | κ | 3-11*01 | 1*01 | QQRSNWPPWT | 10 | 1 | 0 | 2 | 0,7 | 1 | 5,7 | 0,1 | 5,7 | ND | ND | ND | ND | ND | ND | ND | ND | ND | 83,5 | ND | 1307 | ND | 3,1 | ND | ND |

Sequence (aa): Red and Blue colors indicate positively and negatively charged amino acids, respectively, in the IgH and IgL complementary determining region (CDR3).
(+) an (-) indicate the numbers of positively and negatively charged amino acids, respectively, in the IgH and IgL complementary determining region (CDR3).
MUT, number of nucleotide somatic mutations.
EC50, half-maximal lethal effective concentration.
CSPR, circumsporozoite precipitation reaction.
ADCP, antibody-dependent cellular phagocytosis.
ND, not determined.

**Table S3**. SPR single-cycle kinetics (SCK) values of CRR antibodies to CSP18

| mAb | $k_{on}$ ($M^{-1}s^{-1}$) | $k_{off}$ ($s^{-1}$) [a] | $K_D$ (pM) [b] | Normalized binding[c] | $R_{max}$ (RU) [d] | $\chi^2$ ($RU^2$) [e] |
|---|---|---|---|---|---|---|
| ISG3.101 | 1,51E+07 | <0,00001 | < 0,66 | 0,0343 | 11,5 | 0,131 |
| ISG3.102 | 4,15E+06 | <0,00001 | < 2,40 | 0,0190 | 11,4 | 0,764 |
| ISG3.122 | 1,43E+07 | <0,00001 | < 0,70 | 0,0712 | 24,1 | 0,767 |
| ISG3.136 | 3,29E+07 | <0,00001 | < 0,30 | 0,0502 | 10,9 | 0,359 |
| ISG4.107 | 2,07E+07 | <0,00001 | < 0,48 | 0,0241 | 3,9 | 0,465 |
| ISG4.135 | 9,01E+07 | <0,00001 | < 0,11 | 0,0447 | 10,3 | 0,303 |
| ISG4.171 | 4,32E+07 | <0,00001 | < 0,23 | 0,0439 | 9,7 | 0,392 |
| ISG5.110 | 6,77E+06 | <0,00001 | < 1,50 | 0,0162 | 9,9 | 0,41 |
| ISG5.116 | 2,42E+07 | <0,00001 | < 0,41 | 0,0446 | 10,3 | 0,299 |
| ISG5.130 | 2,32E+07 | <0,00001 | < 0,43 | 0,0410 | 10,3 | 0,208 |
| ISG5.138 | 5,91E+07 | <0,00001 | < 0,17 | 0,0347 | 14,1 | 0,16 |
| ISG5.143 | 8,77E+07 | <0,00001 | < 0,11 | 0,0400 | 10,6 | 0,175 |
| ISG5.144 | 1,20E+07 | <0,00001 | < 0,83 | 0,0291 | 10,6 | 0,186 |
| ISG5.154 | 1,13E+07 | <0,00001 | < 0,88 | 0,0276 | 6,8 | 0,293 |
| ISG5.163 | 2,74E+07 | <0,00001 | < 0,36 | 0,0402 | 10,4 | 0,168 |
| ISG5.166 | 9,55E+06 | <0,00001 | < 1,00 | 0,0326 | 10,1 | 0,314 |
| ISG5.167 | 3,47E+07 | <0,00001 | < 0,29 | 0,0432 | 11,8 | 0,148 |
| ISG5.179 | 3,89E+06 | <0,00001 | < 2,60 | 0,0149 | 10,9 | 0,357 |
| ISG6.104 | 8,48E+06 | <0,00001 | < 1,20 | 0,0264 | 8,5 | 0,5 |
| ISG6.106 | 4,61E+06 | <0,00001 | < 2,20 | 0,0153 | 10,3 | 0,142 |
| ISG6.118 | 1,48E+07 | <0,00001 | < 0,68 | 0,0324 | 7,1 | 0,177 |
| ISG6.139 | 8,87E+06 | 1,87E-04 | 21,10 | 0,0396 | 23 | 0,754 |
| ISG6.149 | 1,22E+07 | <0,00001 | < 0,82 | 0,0615 | 22,3 | 0,943 |
| ISG6.160 | 2,61E+07 | <0,00001 | < 0,38 | 0,0665 | 23,3 | 1,07 |
| ISG6.167 | 5,86E+06 | <0,00001 | < 1,70 | 0,0260 | 24,1 | 0,625 |
| ISG6.170 | 4,49E+06 | <0,00001 | < 2,20 | 0,0196 | 11,6 | 0,563 |
| ISG8.131 | 4,43E+06 | <0,00001 | < 2,30 | 0,0300 | 16,9 | 0,751 |
| ISG8.132 | 5,78E+06 | <0,00001 | < 1,70 | 0,0435 | 31,9 | 0,762 |
| ISG8.143 | 1,36E+07 | <0,00001 | < 0,74 | 0,0743 | 28,2 | 0,687 |
| ISG8.151 | 1,20E+07 | <0,00001 | < 0,83 | 0,0667 | 22,6 | 0,613 |
| ISG9.108 | 4,62E+06 | <0,00001 | < 2,20 | 0,0184 | 10,3 | 0,782 |

[a] The slow dissociation rate precluded an accurate measurement. 0,00001 is the limit of detection of the equipment. [b] KD values expressed with < correspond to the upper limit of picomolar range. [c] The analyte binding level divided by the ligand capture level. This corrects for variations in target density to allow direct comparison of binding efficiency. [d] The maximum response or maximum binding capacity. [e] $\chi^2$ indicates the goodness of the overall modeling; the lower the value, the better is the fit.

**Table S4.** Affinities of ISG.5.143, ISG5.144, ISG5.163 and ISG6.149 antibodies and their fully germline-reverted variants

| | **CSP18** | | | | **$NANP_5$** | | | | **$NANP_3$** | | | |
|---|---|---|---|---|---|---|---|---|---|---|---|---|
| | $k_{on}$ $(M^{-1}s^{-1})$ | $k_{off}$ $(s^{-1})$ | $K_D$ $(M)$ | $\chi^2 (RU^2)$ | $k_{on}$ $(M^{-1}s^{-1})$ | $k_{off}$ $(s^{-1})$ | $K_D$ $(M)$ | $\chi^2$ $(RU^2)$ | $k_{on}$ $(M^{-1}s^{-1})$ | $k_{off}$ $(s^{-1})$ | $K_D$ $(M)$ | $\chi^2$ $(RU^2)$ |
| ISG5.143 | 3,7E+07 | 1,6E-03 | 4,3E-11 | 0,1 | 4,6E+06 | 4,0E-03 | 7,4E-10 | 4,3 | n.d | n.d | n.d | n.d |
| ISG5.143GL | n.d | n.d | n.d | n.d | n.d | n.d | n.d | n.d | n.d | n.d | n.d | n.d |
| ISG5.144 | 7,9E+06 | 6,6E-04 | 8,3E-11 | 0,1 | 9,6E+05 | 1,3E-03 | 1,2E-09 | 0,2 | 6,2E+05 | 4,3E-02 | 7,3E-08 | 0,9 |
| ISG5.144GL | 1,4E+07 | 3,5E-03 | 2,5E-10 | 0,077 | 3,4E+06 | 1,4E-02 | 4,1E-09 | 0,592 | n.d | n.d | n.d | n.d |
| ISG5.163 | 5,7E+07 | 3,5E-03 | 5,7E-11 | 0,3 | 9,2E+06 | 1,1E-02 | 1,5E-09 | 3,1 | 7,4E+05 | 9,5E-02 | 1,3E-07 | 0,2 |
| ISG5.163GL | 1,7E+07 | 3,7E-03 | 2,3E-10 | 0,0923 | 4,8E+06 | 5,5E-02 | 1,1E-08 | 3 | n.d | n.d | n.d | n.d |
| ISG6.149 | 1,8E+07 | 1,1E-03 | 6,3E-11 | 0,1 | 1,9E+06 | 1,2E-03 | 6,5E-10 | 0,7 | 9,0E+05 | 2,9E-02 | 4,5E-08 | 0,1 |
| ISG6.149GL | 2,6E+07 | 3,5E-03 | 1,3E-10 | 0,0563 | 1,0E+07 | 1,3E-02 | 1,3E-09 | 2,53 | n.d | n.d | n.d | n.d |

n.d.: not determined due to inaccurate fitting and/or unspecific binding detected by surface plasmon resonance. $\chi^2$ indicates the goodness of the overall modeling; the lower the value, the better is the fit.

**Tables S5**. CryoEM data collection and refinement statistics for the Fab-rsCSP complexes

| | ISG5.143 (EMDB-79206) (PDB 38XB ) | ISG5.144 (EMDB-79207) (PDB 38XC ) | ISG5.163 (EMDB-79205) (PDB 38XA ) | ISG6.149 (EMDB-79208) (PDB 38XD ) | ISG6.149GL_HC (EMDB-79210) (PDB 38XF ) | ISG6.149GL_LC (EMDB-79209) (PDB 38XE) |
|---|---|---|---|---|---|---|
| **Data collection and processing** | | | | | | |
| Magnification | 190x | 190x | 190x | 190x | 190x | 190x |
| Voltage (kV) | 200 | 200 | 200 | 200 | 200 | 200 |
| Electron exposure (e–/Å$^2$) | 60 | 60 | 60 | 60 | 60 | 60 |
| Defocus range (μm) | -0.8 to -1.8 | -0.8 to -1.8 | -0.8 to -1.8 | -0.8 to -1.8 | -0.8 to -1.8 | -0.8 to -1.8 |
| Pixel size (Å) | 0.725 | 0.725 | 0.725 | 0.725 | 0.718 | 0.718 |
| Symmetry imposed | C1 | C1 | C1 | C1 | C1 | C1 |
| Initial particle images (no.) | 725 876 | 1 007 757 | 736 123 | 858 516 | 1 305 259 | 1 302 979 |
| Final particle images (no.) | 361 978 | 75 841 | 51 617 | 492 414 | 138 580 | 409 848 |
| Map resolution (Å) | 3.13 | 3.55 | 3.39 | 2.87 | 3.17 | 2.90 |
| FSC threshold | 0.143 | 0.143 | 0.143 | 0.143 | 0.143 | 0.143 |
| **Refinement** | | | | | | |
| Map sharpening *B* factor (Å$^2$) | -100.4 | -85.9 | -65.3 | -75.8 | -66.4 | -87.2 |
| *Model composition* | | | | | | |
| Non-hydrogen atoms | 9200 | 5425 | 10790 | 10773 | 7112 | 8960 |
| Protein residues | 1191 | 701 | 1402 | 1396 | 921 | 1158 |
| *B factors (Å$^2$)* | | | | | | |
| Protein | 49.34 | 82.92 | 74.78 | 83.47 | 36.93 | 37.86 |
| *R.m.s. deviations* | | | | | | |
| Bond lengths (Å) | 0.004 | 0.004 | 0.004 | 0.003 | 0.004 | 0.004 |
| Bond angles (°) | 0.630 | 0.778 | 0.771 | 0.644 | 0.790 | 0.728 |
| *Validation* | | | | | | |
| MolProbity score | 1.66 | 1.61 | 1.71 | 1.67 | 1.83 | 1.58 |
| Clashscore | 6.85 | 4.42 | 5.48 | 6.25 | 7.38 | 5.87 |
| Poor rotamers (%) | 0 | 0 | 0 | 0 | 0 | 0 |
| *Ramachandran plot* | | | | | | |
| Favored (%) | 95.86 | 94.32 | 93.81 | 95.33 | 93.57 | 96.2 |
| Allowed (%) | 4.14 | 5.68 | 6.19 | 4.67 | 6.43 | 3.8 |
| Disallowed (%) | 0 | 0 | 0 | 0 | 0 | 0 |

**Table S6**. Helical parameters of Fab-rsCSP cryo-EM structures

| | Rise (Å) | | Helical turn (°) | |
|---|---|---|---|---|
| | Mean | SD | Mean | SD |
| ISG5.144 | 6.76825 | 2.38952595 | 101.5338 | 5.19850763 |
| ISG5.163 | 7.163525 | 3.76488567 | 103.57395 | 8.51480138 |
| ISG5.143 | 12.23145 | 0.22274644 | 85.877 | 0.54801801 |
| ISG6.149 | 14.26004 | 0.14077494 | 73.94268 | 0.21377582 |

**Table S7.** X-ray data collection and refinement statistics for matured Fabs

| | ISG6.149 | ISG5.144 | ISG5.163 |
|---|---|---|---|
| **Data collection** | | | |
| Beamline | SSRL 12-1 | SSRL 12-1 | SSRL 12-1 |
| Wavelength (Å) | 0.97946 | 0.97946 | 0.97946 |
| Resolution (Å) | 50.00-2.25 (2.29-2.25)[a] | 50.00-1.99 (2.03-1.99)[a] | 50.00-2.21 (2.25-2.21)[a] |
| Space group | $P2_1$ | $P2_12_12_1$ | $P2_12_12_1$ |
| Unit cell a, b, c (Å) | 90.64, 73.78, 106.16 | 47.48, 74.51, 135.95 | 56.95, 71.98, 134.04 |
| α, β, γ (°) | 90, 100.9, 90 | 90, 90, 90 | 90, 90, 90 |
| Unique reflections | 63,026 (3153)[a] | 33,416 (1606)[a] | 28,292 (1375)[a] |
| Redundancy | 3.5 (3.5)[a] | 5.4 (3.1)[a] | 6.3 (5.5)[a] |
| Completeness (%) | 96.1 (97.6)[a] | 99.2 (97.6)[a] | 99.5 (99.9)[a] |
| Mean I/sigma($\sigma_I$) | 18.2 (2.0)[a] | 10.8 (1.4)[a] | 27.4 (4.8)[a] |
| $R_{sym}$ (%)[b] | 5.5 (50.5)[a] | 16.2 (92.9)[a] | 11.4 (91.9)[a] |
| $R_{pim}$ (%)[b] | 4.1 (33.9)[a] | 9.2 (53.8)[a] | 5.6 (30.7)[a] |
| $CC_{1/2}$ (%)[c] | 99.3 (71.4)[a] | 98.1 (53.4)[a] | 99.1 (84.5)[a] |
| **Refinement statistics** | | | |
| Resolution (Å) | 47.54-2.25 | 44.83-1.99 | 49.05-2.21 |
| Reflections (work) | 62,736 | 33,273 | 28,206 |
| Reflections (test) | 1,999 | 1,998 | 2,000 |
| $R_{cryst}$[d] / $R_{free}$[e] (%) | 21.2/24.0 | 19.1/23.8 | 20.9/24.6 |
| *Number of atoms* | | | |
| Fab | 6,610 | 3,304 | 3,294 |
| Ligand | 138 | 69 | 70 |
| Water | 101 | 160 | 99 |
| *Average B-value (Å$^2$)* | | | |
| Fab | 44 | 24 | 32 |
| Water | 40 | 27 | 22 |
| Wilson B (Å$^2$) | 40 | 21 | 12 |
| *RMSD from ideal geometry* | | | |
| Bond angle (°) | 0.57 | 1.15 | 0.66 |
| Bond length (Å) | 0.002 | 0.011 | 0.004 |
| *Ramachandran statistics*[f] | | | |
| Favored (%) | 98.05 | 97.24 | 98.22 |
| Outliers (%) | 0.00 | 0.00 | 0.00 |
| **PDB Code** | 38VS | 38VP | 38VO |

[a] All numbers in parentheses refer to the highest resolution shell.
[b] Rsym = Σhkl Σi | Ihkl,i - | / Σhkl Σi Ihkl,i and $R_{pim}$ = Σhkl (1/(n-1))1/2 Σi | Ihkl,i - | / Σhkl Σi Ihkl,i, where Ihkl,i is the scaled intensity of the ith measurement of reflection h, k, l, is the average intensity for that reflection, and n is the redundancy.
[c] $CC_{1/2}$ = Pearson correlation coefficient between two random half datasets.
[d] $R_{cryst}$ = Σhkl | Fo - Fc | / Σhkl | Fo | x 100, where Fo and Fc are the observed and calculated structure factors, respectively.
[e] $R_{free}$ was calculated as for Rcryst, but on a test set comprising 5% of the data excluded from refinement.
[f] From MolProbity (Williams CJ, et al., (2018). Protein Sci 27:293-315)

**Table S8.** X-ray data collection and refinement statistics for germline-reverted Fabs

| | ISG6.149GL | ISG5.144GL | ISG5.163GL |
|---|---|---|---|
| **Data collection** | | | |
| Beamline | SSRL 12-1 | SSRL 12-1 | SSRL 12-1 |
| Wavelength (Å) | 0.97946 | 0.97946 | 0.97946 |
| Resolution (Å) | 30.00-1.99 (2.03-1.99)[a] | 40.00-1.98 (2.03-1.98)[a] | 40.00-2.29 (2.35-2.29)[a] |
| Space group | $P2_1$ | $P2_12_12_1$ | $P2_12_12_1$ |
| Unit cell a, b, c (Å) | 69.99, 123.00, 73.59 | 71.79, 84.04, 187.44 | 47.16, 74.83, 141.65 |
| α, β, γ (°) | 90, 93.4, 90 | 90, 90, 90 | 90, 90, 90 |
| Unique reflections | 81,965 (3923)[a] | 75,678 (3814)[a] | 22,506 (1138)[a] |
| Redundancy | 3.2 (2.8)[a] | 6.0 (6.6)[a] | 6.1 (6.5)[a] |
| Completeness (%) | 97.2 (93.5)[a] | 95.1 (97.2)[a] | 96.6 (99.6)[a] |
| Mean I/sigma($\sigma_I$) | 13.3 (1.1)[a] | 22.0 (3.9)[a] | 36.2 (17.5)[a] |
| $R_{sym}$ (%)[b] | 7.2 (58.2)[a] | 6.1 (24.4)[a] | 6.1 (11.5)[a] |
| $R_{pim}$ (%)[b] | 6.2 (41.1)[a] | 3.3 (13.2)[a] | 2.6 (4.7)[a] |
| $CC_{1/2}$ (%)[c] | 98.2 (63.7)[a] | 99.9 (97.7)[a] | 99.5 (99.0)[a] |
| **Refinement statistics** | | | |
| Resolution (Å) | 29.91-1.99 | 39.24-1.98 | 30.48-2.29 |
| Reflections (work) | 81,684 | 75,192 | 22,427 |
| Reflections (test) | 2,004 | 1,998 | 2,000 |
| $R_{cryst}$[d] / $R_{free}$[e] (%) | 20.2/22.6 | 20.5/24.8 | 20.8/25.6 |
| *Number of atoms* | | | |
| Fab | 6,566 | 6,636 | 3,290 |
| Ligand | 138 | 138 | 61 |
| Water | 322 | 333 | 154 |
| *Average B-value (Å$^2$)* | | | |
| Fab | 34 | 31 | 18 |
| Water | 35 | 32 | 20 |
| Wilson B (Å$^2$) | 32 | 27 | 17 |
| *RMSD from ideal geometry* | | | |
| Bond angle (°) | 0.75 | 0.83 | 0.64 |
| Bond length (Å) | 0.005 | 0.006 | 0.003 |
| *Ramachandran statistics*[f] | | | |
| Favored (%) | 97.56 | 97.94 | 98.15 |
| Outliers (%) | 0.00 | 0.00 | 0.00 |
| **PDB Code** | 38VR | 38VQ | 38VN |

[a] All numbers in parentheses refer to the highest resolution shell.
[b] Rsym = Σhkl Σi | Ihkl,i - | / Σhkl Σi Ihkl,i and $R_{pim}$ = Σhkl (1/(n-1))1/2 Σi | Ihkl,i - | / Σhkl Σi Ihkl,i, where Ihkl,i is the scaled intensity of the ith measurement of reflection h, k, l, is the average intensity for that reflection, and n is the redundancy.
[c] $CC_{1/2}$ = Pearson correlation coefficient between two random half datasets.
[d] $R_{cryst}$ = Σhkl | Fo - Fc | / Σhkl | Fo | x 100, where Fo and Fc are the observed and calculated structure factors, respectively.
[e] $R_{free}$ was calculated as for Rcryst, but on a test set comprising 5% of the data excluded from refinement.
[f] From MolProbity (Williams CJ, et al., (2018). Protein Sci 27:293-315).
